# Modality Independent or Modality Specific? Common Computations Underlie Confidence Judgements in Visual and Auditory Decisions

**DOI:** 10.1101/2022.10.31.514447

**Authors:** Rebecca K West, William J Harrison, Natasha Matthews, Jason B Mattingley, David K Sewell

**Affiliations:** School of Psychology, University of Queensland, Australia; Queensland Brain Institute, University of Queensland, Australia; Canadian Institute for Advanced Research, Canada

## Abstract

Humans possess the ability to evaluate their confidence in a range of different decisions. In this study, we investigated the computational processes that underlie confidence judgements and the extent to which these computations are the same for perceptual decisions in the visual and auditory modalities. Participants completed two versions of a categorisation task with visual or auditory stimuli and made confidence judgements about their category decisions. In each modality, we varied both evidence strength, (i.e., the strength of the evidence for a particular category) and sensory uncertainty (i.e., the intensity of the sensory signal). We evaluated several classes of models which formalise the mapping of evidence strength and sensory uncertainty to confidence in different ways: 1) unscaled evidence strength models, 2) scaled evidence strength models, and 3) Bayesian models. Our model comparison results showed that across tasks and modalities, participants take evidence strength and sensory uncertainty into account in a way that is consistent with the scaled evidence strength class. Notably, the Bayesian class provided a relatively poor account of the data across modalities, particularly in the more complex categorisation task. Our findings suggest that a common process is used for evaluating confidence in perceptual decisions across domains, but that the parameter settings governing the process are tuned differently in each modality. Overall, our results highlight the impact of sensory uncertainty on confidence and the unity of metacognitive processing across sensory modalities.

**Author Summary:** In this study, we investigated the computational processes that describe how people derive a sense of confidence in their decisions. In particular, we determined whether the computations that underlie the evaluation of confidence for a visual decision are the same as those for an auditory decision. We tested a range of different models from 3 distinct classes which make different predictions about the computations that are used. We found that a single class of models provided the best account of confidence, suggesting a common process for evaluating confidence across sensory modalities. Even though these computations are governed by the same general process, our results suggest that the process is still fine-tuned within each modality.

## Introduction

Humans possess a remarkable ability to flexibly and reliably evaluate their uncertainty in a range of decisions. Whether reflecting on knowledge (e.g., How sure am I that the capital of Australia is Canberra?), perception (e.g., Is that a cyclist on the road ahead?), or value judgements (e.g., Do I want to buy the latest smart phone?), people have an awareness of the quality of their decisions, even in the absence of explicit feedback (Deroy et al., 2016; Lempert et al., 2015). This awareness arises from a self-monitoring reflective process, termed *metacognition*, which allows people to assess the quality of their internal cognitive operations (Aitchison et al., 2015).

Metacognition has been quantified experimentally using retrospective confidence judgements of performance. In a typical paradigm, an observer makes a task response, typically referred to as the *Type 1* or *first-order* decision, and then reports their level of confidence in that response, referred to as the *Type 2* or *second-order* decision (Fitzgerald et al., 2017). Using confidence judgements, several studies have reported a high degree of association between observers’ task accuracy and their confidence. These accurate metacognitive evaluations of performance have been reported in a range of visual, auditory, tactile, memory, arithmetic, reading, spelling and general-knowledge tasks (Annevirta et al., 2007; Bellon et al., 2020; Faivre et al., 2018; Fleck et al., 2006; Heereman et al., 2015; Koriat, 1993; McCurdy et al., 2013; Rademaker et al., 2012; Rinne & Mazzocco, 2014; Samaha & Postle, 2017; Sanders et al., 2016; Song et al., 2011). In many of these contexts, observers evaluate their uncertainty in tasks which contain just a single stream of information. In the natural environment, however, we typically evaluate information from several different sensory and cognitive domains at any one time. For example, when deciding whether to buy a particular pineapple at the grocery store, one might evaluate their confidence that the pineapple’s colour, smell and texture indicate that it’s ripe.

Currently, little is known about the computations that support the diversity and flexibility with which people monitor their confidence. In particular, it is unclear whether the computations used to evaluate confidence are functionally and algorithmically related across different sensory and cognitive domains. To address this question, we compared metacognitive judgements across two sensory modalities, vision and audition. We chose to compare confidence across modalities as it allowed us to keep the decision context the same but vary the domain of information presentation. Thus, we could isolate any similarities (*modality-independent* processes) or differences (*modality-specific* processes) in the computations used to generate confidence.

### Modality-Independent or Modality-Specific Metacognition?

There are theoretical justifications for expecting metacognitive processes to be either modality-specific or modality-independent. Sensory information in different modalities has different features, and the different sensory systems respond selectively to those features. Modality-specific metacognitive processes can capitalise on this specialised encoding and maximise the accuracy of uncertainty monitoring within a modality. In addition, we do not rely on all of our senses equally (Alais & Burr, 2004; Ernst & Banks, 2002; Hillis et al., 2002; Treisman, 1998). We may use optimal but computationally expensive processes for the senses we use most often, such as vision (Knill & Richards, 1996; Körding, 2007; Ma & Jazayeri, 2014), and have more heuristic, efficient metacognitive processes for other modalities, for example audition. Modality independent processes, on the other hand, may be equally advantageous. The sensory environment is inherently multi-modal in nature. In the real world we consistently filter out irrelevant information from certain modalities, make comparative judgements and integrate information across modalities. Transforming modality-specific sensory input into a modality-independent confidence variable would create a common currency for flexible comparison and integration of uncertainty across domains (de Gardelle et al., 2016; de Gardelle & Mamassian, 2014).

To compare these competing accounts, we evaluated a number of existing models which formalise the computations involved in generating choice and confidence judgements, and compared the ability of such models to account for human performance in visual and auditory categorisation tasks. We consider three broad classes of existing confidence models: 1) unscaled evidence strength models, 2) scaled evidence strength models, and 3) Bayesian models. We provide an overview of these confidence models below. To illustrate the similarities and differences between models, in the following sections we focus on their implementation in a two-alternative force choice task (2AFC) where an observer decides if a Gabor patch is rotated clockwise or counter-clockwise relative to horizontal (a category decision; see two example gratings in Fig 1A) and evaluates their confidence in that decision.

**Fig 1.**
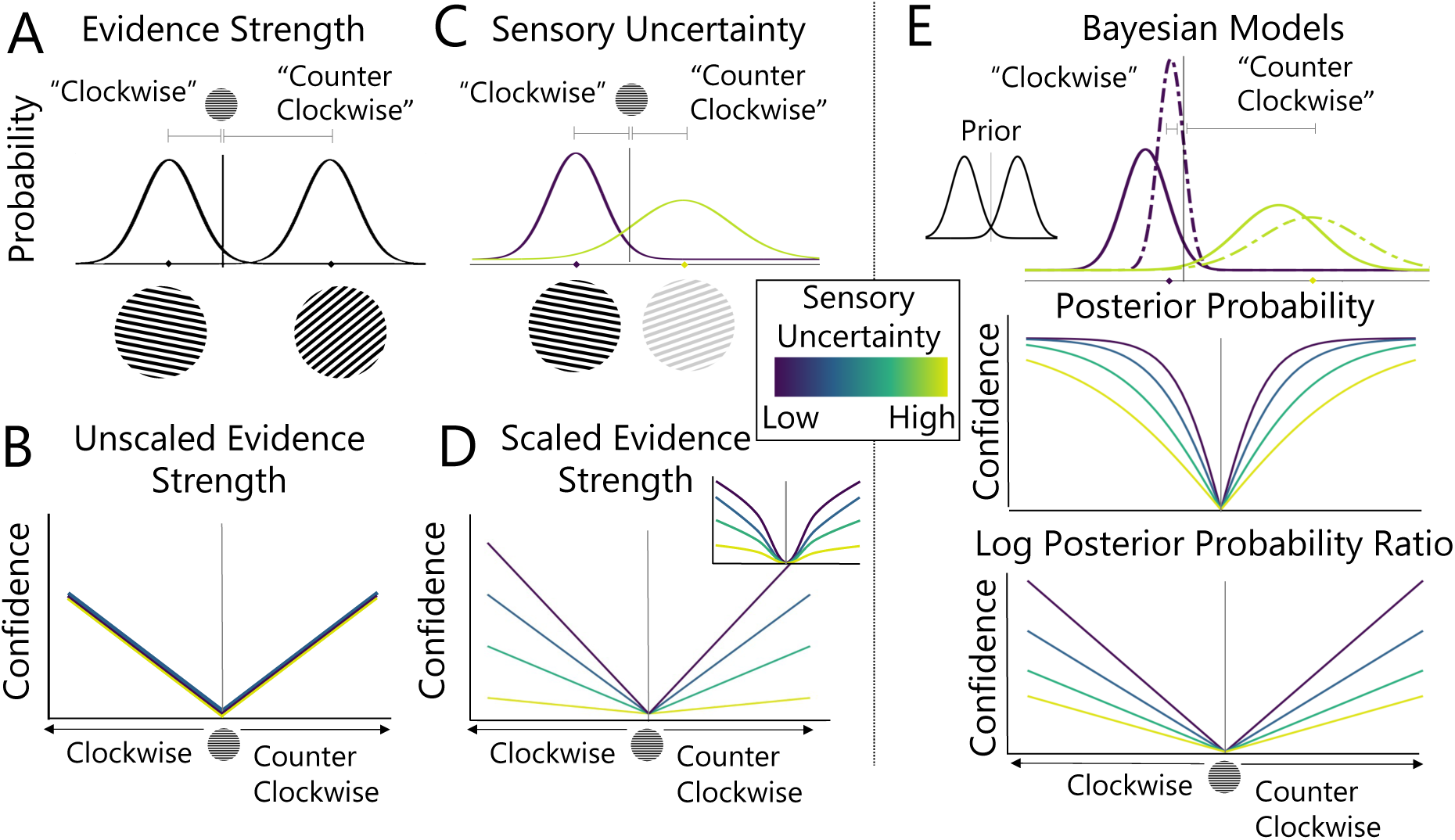
Confidence Models Tested in the Current Study. Illustration of (A) evidence strength and (C) sensory uncertainty concepts for confidence models. (B) Unscaled Evidence Strength models only factor in evidence strength. (D) Scaled Evidence and (E) Bayesian models factor in both evidence strength and sensory uncertainty using different principles. Scaled Evidence strength models assume a parametric relationship between confidence and uncertainty. Linear and quadratic (inset in D) examples are shown here. Bayesian models describe confidence in terms of the posterior probability or the log posterior probability ratio of competing outcomes.

### Confidence Models

#### Evidence Strength Models

Evidence strength models posit that an observer’s confidence depends on the distance between the sensory measurement and the decision criterion, whereby increasing distance from the criterion would result in increasing confidence. To illustrate this, see Fig 1A which shows two noisy measurements of the orientations of two Gabor patches with different distances from the category decision criterion. For numerical confidence ratings with *N* levels, *N*-1 criteria can be used to construct confidence bins that delineate the mapping between distance and confidence ratings. Evidence strength models can be further differentiated based on whether the positioning of confidence criteria takes sensory uncertainty, the amount of noise in the measurement of the stimulus (which may vary, for example, with the contrast of the Gabor patch, see Fig 1C) into account.

##### Unscaled Evidence Strength

According to unscaled evidence strength models, confidence reports depend only on the unsigned distance between the sensory measurement and the decision criterion (Kepecs et al., 2008; Komura et al., 2013; Rahnev et al., 2011). These models do not independently quantify sensory uncertainty, such that a given distance will always be associated with the same level of confidence, regardless of other properties of the stimulus. These are perhaps the simplest models to reject, because any empirical difference in observers’ confidence judgments across levels of sensory uncertainty cannot be accounted for by the model.

##### Scaled Evidence Strength

Scaled evidence strength models allow a point estimate of sensory uncertainty to scale the confidence criteria used to delineate distance (Baranski & Petrusic, 1994; Kiani, Corthell, & Shadlen, 2014; Barthelme & Mamassian, 2010; Adler & Ma, 2018; Locke et al., 2021). In effect this means that under conditions of high sensory uncertainty (light green line in Fig 1C, D), greater evidence strength is required to produce the same level of confidence as under conditions of low sensory uncertainty (purple line in Fig 1C, D). Several models fall into this class where different scaling rules can be applied. Adler and Ma (2018), for example, tested models where category and confidence response criteria were estimated as linear (depicted in main plot of Fig 1D) or quadratic (depicted in inset of Fig 1D) functions of sensory uncertainty. Here we consider category and confidence criteria that are linear and quadratic (and other) monotonic functions of sensory uncertainty, but we do not draw any theoretical distinctions between the different scaling rules.

#### Bayesian Models

Bayesian confidence models constitute a statistically optimal way of mapping both task and sensory information to confidence. In Bayesian models, observers combine knowledge about the statistical structure of the task (the “prior”) with the present sensory input (the “likelihood”) to compute a posterior probability distribution over possible states of the stimulus (Drugowitsch et al., 2014b; Hangya et al., 2016; Kepecs and Mainen, 2012; Meyniel et al., 2015; Pouget et al., 2016). Bayesian models therefore differ from evidence strength models because they involve the computation of a probability distribution of all possible states of the stimulus. Confidence judgements are then derived from the probability metric. For the clockwise vs. counter clockwise decision illustrated in Fig 1, where *x* is the measurement of the orientation of the Gabor patch, a Bayesian observer would report clockwise whenever *p*(*clockwise* | *x*) is above 0.5. The more extreme the posterior probability, the more confident the observer can be in their response. An alternative way of mapping posterior probability to confidence is to use the log posterior probability ratio, 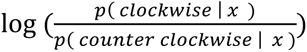, which is unbounded and allows for greater differences in confidence at different levels of sensory uncertainty (see bottom panel of Fig 1E; Adler & Ma, 2018; Locke et al., 2022).

#### Model Comparison

To date, formal comparisons of these different classes of confidence models have focused on the visual domain and have yielded conflicting results with some studies finding that visual confidence judgements are most consistent with Bayesian models (Aitchison et al., 2015; Hangya et al., 2016; Li & Ma, 2020; Navajas et al., 2017; Sanders et al., 2016) and others finding evidence for non-Bayesian models (Adler & Ma, 2018; Aitchison et al., 2015; Bertana et al., 2021; Lisi et al., 2021; Locke et al., 2021). Thus, not only are there are demonstrated inconsistencies in research findings within the visual domain and but it is also currently unclear how well the existing set of confidence models can account for confidence judgements across modalities. A formal investigation of the computations used to generate confidence across modalities and tasks is needed.

### The Current Study

The aim of the present study was to investigate the computations used to generate confidence judgements across sensory modalities by comparing the fits of different confidence models to data from different modalities. We had participants undertake two versions of a categorisation task, one requiring decisions on visual stimuli and the other, decisions on auditory stimuli. We equated task demands across modalities to allow us to distinguish differences in metacognitive processes from overall changes in task performance. We varied both evidence strength (i.e., the strength of the evidence for a particular category) and sensory uncertainty (i.e., the intensity of the sensory signal) in each modality. To preview our results, we found that a single class of models, namely scaled evidence strength models, provided the best account of confidence across modalities and tasks. However, while a single class of models provided the best account across modalities and tasks, we found it is unlikely that this metacognitive process operates under the same parameter settings across modalities. We tested several levels of flexibility in the parameter settings of a model fit to data from both modalities and found that the best model was one in which all the parameters were estimated separately in each modality. We propose that a scaled evidence strength algorithm tuned to the specific information in each sensory modality provides the best account of metacognition across domains.

## Method

### Overview

Participants completed two categorisation tasks in each of two sensory modalities, vision and audition. On each trial, participants made a two-alternative forced choice category decision, followed by a 4-point confidence rating, ranging from low confidence (1) to high confidence (4). The categorisation tasks were based on those used by Adler and Ma (2018). Category membership was determined by a single stimulus attribute: orientation for visual stimuli (drifting Gabor patches) and pitch (frequency) for auditory stimuli (pure tones). A normal distribution of the relevant stimulus attribute defined each category and the parameters of these category distributions differed across two versions of the task: a *different means* task and a *different standard deviations* (SDs) task. In the different means task, category distributions had different means and the same standard deviation. In the different SDs task, category distributions had the same mean and different standard deviations. The category distributions are described in detail below. In both tasks, sensory uncertainty was also varied for each modality, with stimuli presented at 4 different intensities. In the visual tasks, intensity depended on the contrast of the Gabor patch (3.3%, 5%, 6.7%, 13.5%). In the auditory tasks, intensity depended on the loudness of the tone (4, 9, 26, 55 phon; phons are the sound pressure level in decibels of a 1000 Hz pure tone that has been subjectively matched in loudness to the target tone).

### Participants

12 participants (*Mage* = 24.83, *SDage* = 4.35) were recruited through The University of Queensland’s research participation scheme and by word-of-mouth. Participants were reimbursed for their time ($20 per hour in cash or gift cards).

Inclusion criteria included normal (or corrected-to-normal) vision and normal hearing, both assessed by self-report.

### Categorisation Tasks

#### Different Means Task

As shown in Fig 2, category distributions in the different means task had different means and the same standard deviation. In the visual modality, where 0 degrees represented a horizontal Gabor patch, orientations that were rotated counter clockwise relative to horizontal (i.e., negative orientations) were more likely to be sampled from category 1 (*µ_cat_*_1_ = −4°, σ*_cat_*_1_ = 5°). Orientations that were rotated clockwise relative to horizontal (i.e., positive orientations) were more likely to be sampled from category 2 (*µ_cat_*_2_ = 4°, σ*_cat_*_2_ = 5°). In the auditory modality, the category distributions had the same properties, but parameter values were shifted into the frequency domain. Lower frequency tones were more likely to be sampled from category 1 (*µ_cat_*_1_ = 2300 *HZ*, σ*_cat_*_1_ = 475 *HZ*) and higher frequency tones were more likely to be sampled from category 2 (*µ_cat_*_2_ = 3100 *HZ*, σ*_cat_*_2_ = 475 *HZ*).

**Fig 2.**
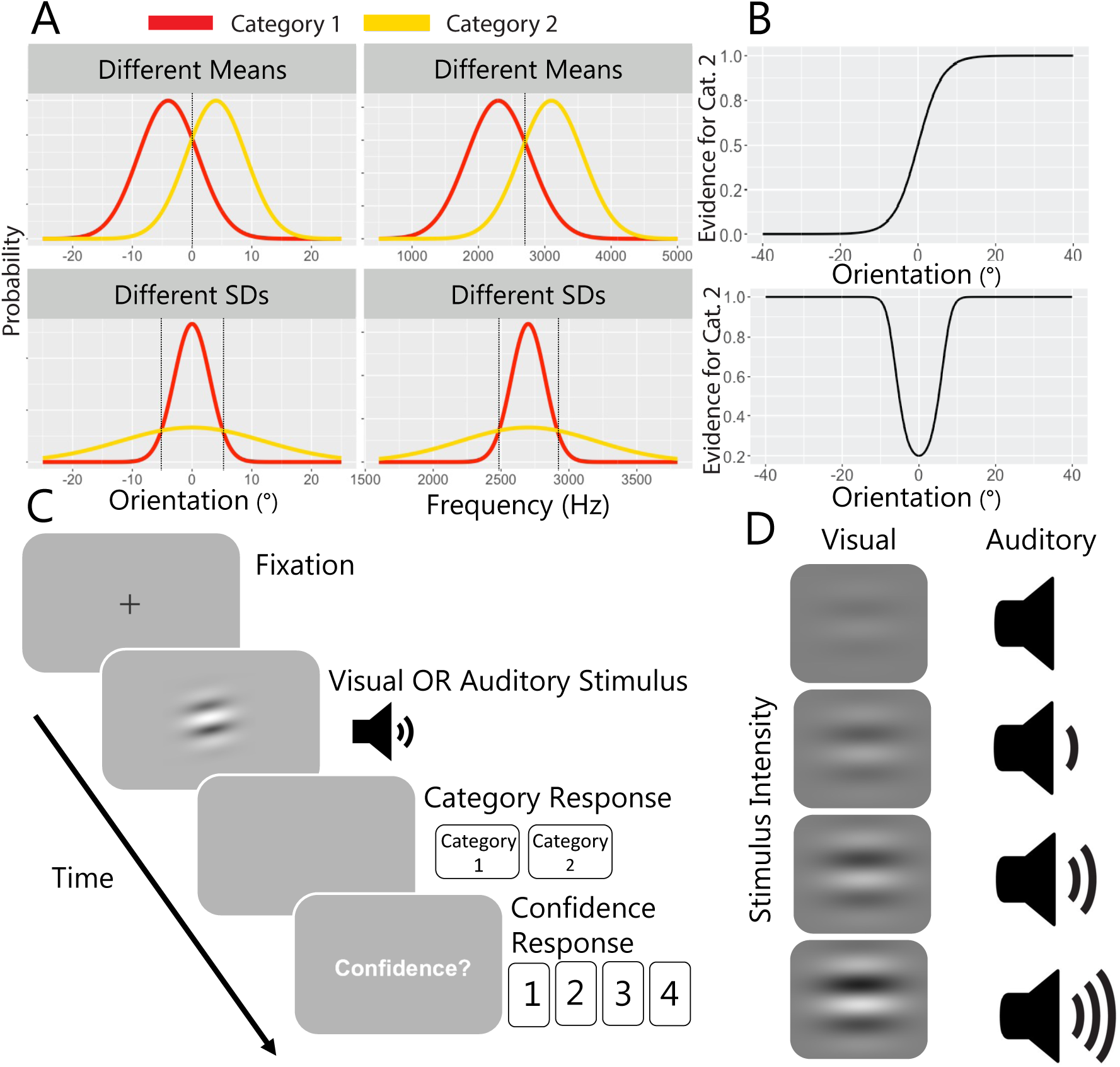
Experimental Task. (A) Category distributions for all conditions. (B) Changes in evidence for category 2 as a function of stimulus value in the different means task (top panel) and the different SDs task (bottom panel). (C) Trial sequence for test trials. (D) Levels of sensory uncertainty across modalities, where Gabor contrast varied in the visual tasks and tone intensity varied in the auditory tasks.

Stimuli with orientation/frequency values equal to where the two category distributions intersected, at 0 degrees for orientations and 2700 Hz for frequencies, were equally likely to be sampled from either category 1 or category 2. Because the relative likelihood of category membership changed monotonically around this point, to maximise correct responses, an ideal observer would report that any stimulus value below this point was sampled from category 1 and any stimulus value above this point was sampled from category 2.

#### Different SDs Task

In the different SDs task, category distributions had the same mean and different standard deviations. In the visual modality, larger rotations clockwise or counter clockwise relative to horizontal were more likely to be sampled from category 2 (*µ_cat_*_2_ = 0°, σ*_cat_*_2_ = 12°) whereas smaller rotations relative to horizontal were more likely to be sampled from category 1 (*µ_cat_*_1_ = 0°, σ*_cat_*_1_ = 3°). In the auditory modality, lower and higher frequency tones were more likely to be sampled from category 2 (*µ_cat_*_2_ = 2700 *HZ*, σ*_cat_*_2_ = 500 *HZ*), whereas intermediate frequency tones were more likely to be sampled from category 1 (*µ_cat_*_1_ = 2700 *HZ*, σ*_cat_*_1_ = 125 *HZ*).

To maximise correct responses, an ideal observer would use the two points where the category distributions intersect, at -5 and 5 degrees for orientations and 2485 Hz and 2915 Hz for frequencies, reporting that stimulus values contained within these intervals were sampled from category 1 and stimulus values outside these intervals were sampled from category 2.

### Stimuli

#### Visual Stimuli

The visual stimuli were drifting Gabors which had a spatial frequency of 0.5 cycles per degrees of visual angle (dva), a speed of 6 cycles per second, a Gaussian envelope with a standard deviation of 1.2 dva, and a randomized starting phase.

Visual stimuli appeared at fixation for 50 ms.

#### Auditory Stimuli

The auditory stimuli were tones of varying frequency (including 5 ms linear onset and offset amplitude ramps to eliminate onset and offset clicks). Stimuli were synthesized with an ASIO4ALL sound driver with a sampling rate of 28 kHz. Tones were matched for loudness according to the International Standard ISO 226:2003: Acoustics- Normal Equal-Loudness-Level-Contours (International Standardization Organisation, 2003) and extensive piloting was undertaken to select intensity levels. Auditory stimuli were presented in stereo via headphones (Sennheiser HD 202) for 50 ms.

### Procedure

All participants completed all combinations of tasks in four separate testing sessions (i.e., visual different means task, visual different SDs task, auditory different means task, auditory different SDs task). Participants completed the *different means* and the *different SDs* task of the same modality on the same day (with at least one hour between sessions) and completed the visual and auditory tasks on separate days. The order of task modality was counterbalanced across participants and task type (different means vs. different SDs) was counterbalanced within modalities and across participants.

Participants were seated in a dark room, at a viewing distance of 57 cm from the screen, and their head was stabilised with a chinrest. Stimuli were presented on a gamma-corrected 60 Hz 1920-by-1080 display. The computer display (ASUS VG248QE Monitor) was connected to a Dell Precision T1700, calibrated with a ColorCal MKII (Cambridge Research Systems). Stimuli were generated and presented using custom code and the Psychophysics Toolbox extensions (Brainard, 1997; Pelli, 1997) for MATLAB.

In each session, participants received instructions for that session’s task, which included an explanation about how the stimuli were generated from the relevant category distributions. To further illustrate these distributions, participants were then shown 36 stimuli randomly sampled from each category. They then completed the category training and testing (detailed below). Each session took approximately 1 hour. Combining all sessions and tasks, participants completed 1440 training trials and 2880 testing trials. Data from training trials were not included in any analyses.

#### Category Training

At the start of each training trial, participants fixated on a central cross for 1 s. The fixation cross was then extinguished, and a stimulus was presented. A stimulus value (i.e., orientation for visual stimuli and frequency for auditory stimuli) was sampled randomly from the relevant category distribution and the stimulus was presented (50 ms duration for auditory stimuli and 300 ms duration for visual stimuli). Immediately after the offset of the stimulus, participants were able to respond category 1 or category 2 by pressing the F or J key on a standard keyboard with their left or right index finger, respectively. No confidence ratings were collected during training. After participants made their response, corrective feedback (i.e., the word “correct” in green or “incorrect” in red) was displayed for 1.1 s. The inter-trial interval was 1 s, after which the fixation cross reappeared. Within a training block, stimuli were sampled equally from each category and the order of categories randomised across trials.

Participants completed 3 training blocks (120 trials per block, in total 360 trials) per session. During training, only the highest intensity level was used for stimulus presentation.

#### Test

The trial procedure in testing blocks was the same as in training blocks, except that trial-to-trial feedback was withheld, stimuli were presented at four different intensity levels (different contrast values for the visual stimuli and different loudness values for the auditory stimuli) and stimuli were presented for 50 ms, regardless of modality. Participants then made a category response, followed by a confidence report, using the 1-4 number keys to indicate their confidence. At the end of each block, participants were required to take at least a 30 s break. During the break, they were shown the percentage of trials they had correctly categorized in the most recent block. Participants were also shown a list of the top 10 block scores (across all participants, indicated by initials) for the task they had just completed. This was intended to motivate participants to maintain a high level of performance.

Participants completed 6 testing blocks per session (120 trials per block, a total of 720 trials per session). Within a testing block, an equal number of test stimuli were sampled from each category and intensity level (i.e., 15 trials per cell of the design). The order of both category and intensity level were randomised.

### Model Specification

#### All Models

##### Measurement Noise

In all models, we assume that the observer’s internal representation of the stimulus, *x*, is a noisy measurement. We approximate this measurement noise using a Gaussian distribution (referred to as the ‘measurement distribution’) in both modalities. Although orientation is circular and traditionally modelled using a von Mises distribution, we used a Gaussian distribution because we only used a small range of orientations in both the visual tasks (Adler & Ma, 2018). The Gaussian distribution was centred on the true stimulus value presented on trial i, *s*), with standard deviation, *σ*. We assume that *σ* depended on stimulus intensity, where greater stimulus intensity was associated with less measurement noise. In contrast to Adler and Ma (2018), we did not enforce a power law relationship between intensity and measurement noise because we were agnostic about the functional form of this relationship. Therefore, the standard deviation of the measurement distribution was estimated separately at each intensity level with a monotonicity constraint,

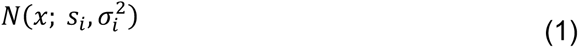

where *N* denotes the normal density function. The standard deviation of the measurement distribution, therefore, approximated the level of sensory uncertainty in the observer’s perception of the stimulus.

##### Orientation Dependent Noise

Following Adler and Ma (2018), for modelling performance in the visual task, we also tested a variation of some models which assume additive orientation-dependent noise in the form of a rectified 2-cycle sinusoid (Girshick et al., 2011).

In these models, the standard deviation of the measurement distribution was

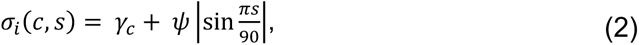

where *c* represents the intensity level, *γ_c_* is a free parameter representing the baseline amount of measurement noise at a given intensity and *ψ* scales the amount of additive measurement noise across stimulus orientations. We did not fit a frequency dependent noise parameter in the auditory tasks as each frequency was corrected for equal loudness prior to adjusting overall intensity.

##### Boundaries

In all models, we assume that participants use a set of *boundaries* to make a response. The boundaries partition the stimulus space into discrete category and confidence response regions so that if a stimulus falls within the defined region, response *r* is given. Given four confidence levels for each of the two categories, we required eight response regions, *r* ∈ {1, 2, 3, …, 8}, each corresponding to a unique category and confidence combination. Given the generating category distributions (see Fig 2), the positioning of these boundaries differs for the different means and the different SDs tasks, and is described below.

#### Different Means Task Boundaries

In the different means task, we assumed that one of the boundaries, *b_1_*, divides the stimulus range into the two different category response regions. That is, where the observed stimulus value is below this boundary, a category 1 response is given and where the observed stimulus value is above this boundary, a category 2 response is given. Confidence associated with the category response is determined by a further three boundaries, *b_5_*, *b_6,_b_8_*, which created four different confidence response regions within each category. Within a category response region, we assumed that the confidence boundaries increased monotonically across successive boundaries such that higher confidence response regions were aligned with more extreme category evidence (see Fig 3B). We also assumed that the confidence boundaries were symmetrically positioned across categories, because two measurements that are equidistant from the intersection of the category response regions (and therefore have the same evidence strength) should logically have the same confidence associated with them. Due to symmetry, confidence regions on equidistant but opposite sides of the category boundary, *b*1, were treated as equivalent by the model. The boundaries in the different means task were ordered such that: −*b_4_* < −*b_3_* < −*b_2_* < *b_1_* < *b_2_* < *b_3_* < *b_4_*.

**Fig 3.**
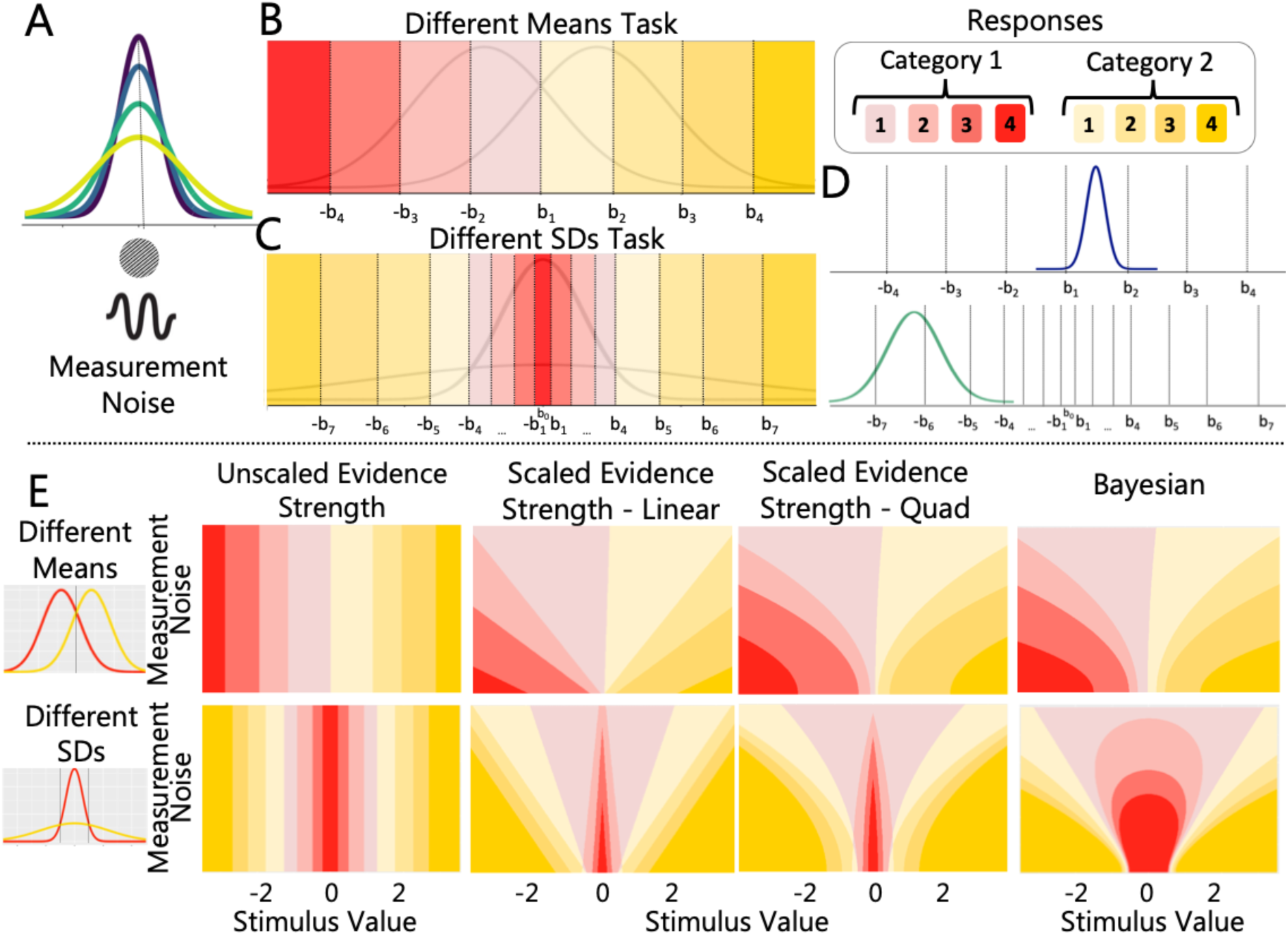
Computational Models. (A) Pictorial representation of the measurement distribution showing how measurement noise varies as a function of stimulus intensity. Illustration of boundary placement and response regions for (B) the different means task and (C) the different SDs task. (D) The measurement distribution is compared to the boundaries. (E) Predicted response regions for the different means (top panel) and different SDs (bottom panel) tasks across different levels of sensory uncertainty and standardised stimulus values for the different models. These predictions were generated using parameter values similar to those found after fitting participant data.

#### Different SDs Task Boundaries

In the different SDs task, two boundaries, −*b_8_* and *b_8_*, divided the stimulus range into two different category response regions. Where a stimulus value falls within the −*b_8_* and *b_8_* region, a category 1 response is given. Where a stimulus value falls beyond −*b_8_* or *b_8_*, a category 2 response is given. To make a confidence response, the observer uses 4 symmetrical boundaries for each category: *b_4_*– *b_8_* for category 1 responses and *b_8_* - *b_9_* for category 2 responses, as shown in Fig 3B. In all models, we assumed that the boundaries increased monotonically across successive boundaries such that higher confidence response regions were aligned with more extreme category evidence (see Fig 3B). We also assumed that the boundaries are symmetrically positioned around the means of the category distributions. For ease of exposition, we refer to this centre point as *b*_&_. We assumed the boundaries are symmetrical in this task because two measurements that are equidistant from the middle of the category structure but on opposite sides of *b*0 should logically have the same confidence associated with them. Due to symmetry, confidence regions on equidistant but opposite sides of *b*0 were treated as equivalent by the model. The boundaries in the different SDs task were ordered such that: −*b_7_* < −*b_6_* < −*b_5_* < −*b_4_* < −*b_3_* < −*b_2_* < −*b_1_* < *b_1_* < *b_2_* < *b_3_* < *b_4_* < *b_5_* < *b_6_* < *b_7_*.

#### Model Descriptions

The distinguishing factor among candidate models concerns how the category- confidence boundaries were positioned in either perceptual space (Distance, Linear, Quadratic & Free-exponent) or probability space (Bayesian; discussed in more detail below) as a function of sensory uncertainty. For models using perceptual space, the resulting boundaries were in units of the stimulus (degrees or frequency) and, therefore, the observer’s internal representation of the stimulus could be compared directly with the boundaries. For models using probability space, the observer’s internal representation of the stimulus was transformed into a probability metric which was then compared with the boundaries. To illustrate, for the different means task in the visual domain, for a non-Bayesian model with category-confidence boundaries in perceptual space, a Category 2 Confidence 1 response might be made for any stimulus perceived as having an orientation between 0 and 5 degrees. For a Bayesian model, where the boundaries are in probability space, the same Category 2 Confidence 1 response might be made for any stimulus perceived as having a log posterior probability ratio between 0 and 0.5.

Assumptions shared across all model classes are detailed above. A description of the distinguishing elements of each model class is provided below. All models were fit to each task and modality separately (except where specified otherwise).

#### Unscaled Evidence Strength Models

##### Distance Model

**I**n the distance model, the boundary positions were not dependent on (estimated) values of sensory uncertainty (*σ*) and therefore occupied fixed positions across intensity levels. The position of each boundary in perceptual space was described by a free parameter, *k*, where perceptual boundaries *b_r_* = *k_r_*.

#### Scaled Evidence Strength Models

##### Linear, Quadratic and Free-Exponent Models

In the linear and quadratic models, the positions of the boundaries were linear or quadratic functions of *σ*. Each boundary position depended on 3 free parameters: *k*, *m* and *σ*. *k,* represented the base position of each boundary which was offset by a scaled value of sigma, ±*mσ* (linear model), or by a scaled value of sigma squared, ±*mσ^5^* (quadratic model), where *m* was fixed for each boundary. The resulting boundaries in perceptual space were given by *b_r_*(*σ*) = *k_r_*+ *m_r_σ* (Linear) and *b_r_*(*σ*) = *k_r_*+ *m_r_σ^5^* (Quadratic). In the free-exponent model, we estimated the exponent on σ, *a*, as a free parameter, *b_r_*(*σ*) = *k_r_*+ *m_r_σ^3^*. The additional free parameter allowed for the changes in boundary position across intensities to occupy values between and beyond linear or quadratic values. We constrained *a* to be greater than 1.

##### Parameter Settings of the Free-Exponent Models Across Modalities

While the primary focus of this study was to determine whether a single class of metacognitive models could account for the way participants make confidence judgements across modalities, we also wanted to determine the extent to which this process was sensitive to different parameter settings. Accordingly, rather than fit the model to each modality separately, we fit the free-exponent model to data from both modalities simultaneously. We formulated several levels of flexibility in the parameter settings, with more flexible versions allowing the parameter values to be different across modalities and less flexible versions constraining parameter values to be the same across modalities. We chose the free-exponent model to test the flexibility in parameter settings, as it was the most flexible of the scaled evidence strength models and includes the linear and quadratic models as special cases.

##### Common Settings Model

For the common settings model, both modalities had the same parameter settings. We assumed that sensory uncertainty (*σ*) did not differ across modalities. Thus, we fit *σ* as a free parameter for each intensity level and used the same value of *σ* across modalities. We also assumed that the same boundary positions were used across modalities so that all perceptual boundaries were given by *b_r_*(*σ*) = *k_r_*+ *m_r_σ^3^*.

##### Common Boundary Settings (Scaled Noise) Model

For the common boundary settings (scaled) model, the settings of the boundary parameters (*k* and *m*) were fixed across modalities. An additional free parameter was then used to scale the sigma parameters in the auditory modality such that *σ*(*c*)*_auditory_* = *wσ*(*c*)*_visual_*, where *w* is the non-negative free parameter and *c* denotes the intensity level. Perceptual boundaries were given by *b_r_*(*σ*) = *k_r_*+ *m_r_σ^3^*.

##### Common Boundary Settings (Offset Noise) Model

For the common boundary settings (offset) model, an additional free parameter was used to add a constant to the sigma parameters in the auditory modality such that *σ*(*c*)*_auditory_* = *p* + *σ*(*c*)*_visual_*, where *p* is the free parameter and *c* denotes the intensity level. Perceptual boundaries were given by *b_r_*(*σ*) = *k_r_*+ *m_r_σ^3^* where *k* and *m* were fixed across modalities.

##### Different Noise Settings Model

For the different noise settings model, sigma was a free parameter for each intensity level in each modality. *k* and *m* were fixed across modalities, assuming that only measurement noise changed across modalities. The perceptual boundaries were given by *b_r_*(*σ*, *t*) = *k_r_*+ *m_r_σ*(*c*, *t*)*^3^*, where *c* is the intensity level and *t* is the sensory modality.

##### Flexible Settings Model

In the flexible settings model, all parameters in the standard free-exponent model were freely estimated in each modality.

#### Bayesian Models

A Bayesian observer optimally combines prior beliefs of the generative model (e.g., the task-specific category distributions) with observed stimulus information to form posterior beliefs about the nature of the stimulus (e.g., category membership). We tested several versions of Bayesian confidence models which make different assumptions, as described below.

##### Log Posterior Probability Ratio

When the perceived value of a stimulus on a given trial is *x*, we assume that the observer uses the log posterior probability ratio, 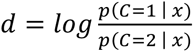, to make a response. Category and confidence reports depend on the internal representation of the stimulus *x* only via *d.* On a given trial, we assume that the observer chooses a response by comparing *d* to a set of boundaries *k* = (*k_4_*, *k_5_* … *k_9_*) that partitions *d* space into eight response regions, each representing a unique category and confidence combination. The positions of these boundaries in *d* units were free parameters. Following Adler and Ma (2018), we used the following task-specific derivations of *d* for the different means task:

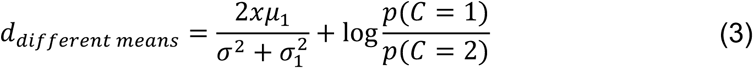

and the different SDs task:

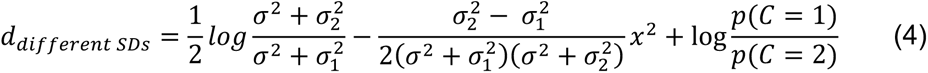

Where *μ_1_* and *σ_1_* are the mean and standard deviation of the category 1 distribution and *μ_2_* and *σ_2_* are the mean and standard deviation of the category 2 distribution. In terms of the generative model, in the different means task the observer needs knowledge of *μ_4_* and *σ_1_* to make an optimal decision and in the different SDs task the observer needs knowledge of *σ_1_* and *σ_2_*to make an optimal decision. Because we sampled equally from each category, we assumed that *p*(*C*=1)=*p*(*C*=2)=0.5 such that 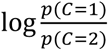 is 0 in both Equation 3 and Equation 4.

##### Log Posterior Probability Ratio with Decision Noise

For the log posterior probability ratio model with decision noise, we assumed that there was an added Gaussian noise term on *d*, *σ_<_*. This allowed for the observer’s calculation of *d* to be noisy, in addition to sensory noise in the internal representation of the stimulus, *σ_)_*(Adler & Ma, 2018).

##### Log Posterior Probability Ratio with Free Category Distribution Parameters

The log posterior probability ratio model assumes that the observer knows the true values of the means and standard deviations of the category distributions, *μ_4_* and *σ_1_*in Equation 3 and *σ_1_* and *σ_2_*in Equation 4. We also tested a version of the model in which the observer had imperfect knowledge of the parameters of the generative model, and estimated these values as free parameters instead.

#### Fitting the Models to Data

To compare parameter values and model fits across modalities, we standardised stimulus values before fitting models to each individual’s raw data. To standardise stimulus values, we subtracted the mean and divided by the standard deviation of all stimulus values for each task in each modality. Model parameters were estimated for each individual using maximum likelihood estimation. Below we describe the process for calculating the likelihood of the dataset, given a model with parameters *θ*, for the non-Bayesian and Bayesian models, as they require different transformations.

##### Non-Bayesian Models

For all non-Bayesian models, the boundaries were positioned in perceptual space and no transformation of boundary positions was required. For each trial, we determined the predicted response probability for each combination of category and confidence level by computing the probability mass of the measurement distribution between adjacent boundaries, given the parameters of the model and the participant’s response on that trial, *r_)_* (cf. Luce, 1959; see Fig 3D). For the different means task, this quantity is:

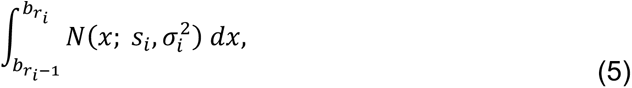

Where *b_5_* = ∞ and −*b_5_* = −∞.

For the different SDs task, this quantity is:

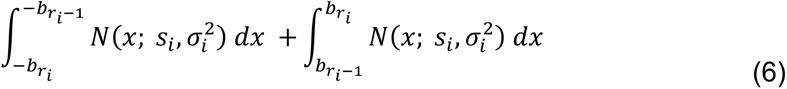

Where *b_0_* = 0, *b_8_* = ∞ and − *b*_8_ = −∞.

##### Bayesian Models

In the Bayesian models, boundaries are positioned in *d*- space (log posterior probability ratio space). To evaluate the probability mass of the measurement distribution, we transformed the boundary positions into perceptual space, *b*(*σ*), using parameters *k* as the left-hand side of Equation (3) or Equation (4) and solved for *x* at the fitted levels of *σ*. Solving Equation (4) for *x* requires calculating the square root of *d*, which is undefined for negative values. We therefore used the absolute value of *d* to solve Equation (4) for *x* and applied the appropriate sign to the result to arrive at a final standardised orientation/frequency value for *x*. Where the category distribution means and standard deviations are free parameters, we used the estimated values in place of the true mean and standard deviations (*μ_4_* and *σ_1_* for the different means task and *σ_1_* and *σ_2_* for the different SDs task). We then calculated the probability mass of the measurement distribution between adjacent perceptual boundaries using Equation (5) or Equation (6), given the task and participant’s response for that trial.

##### Log Posterior Probability Ratio with Decision Noise

For the log posterior probability ratio model with decision noise, we took 101 evenly spaced draws from a normal distribution (spanning the 1^st^ to 99^th^ distribution percentiles) for each boundary, with *k* as the mean and the standard deviation as a free parameter, *σ_<_*. Each of the 101 draws was then converted into perceptual space using Equation (3) or Equation (4) to solve for *x* at the fitted levels of *σ*. We then calculated the probability mass of the measurement distribution between corresponding draws of the relevant boundaries, given the participant’s response for that trial. This means that for a single trial we had 101 response probabilities. We calculated the weighted sum of these probabilities using the normalised densities for each draw of *k*.

##### Log Likelihood of the Dataset

To obtain the log likelihood of the dataset, we computed the sum of the log probability for every trial *i,* where *t* is the total number of trials.

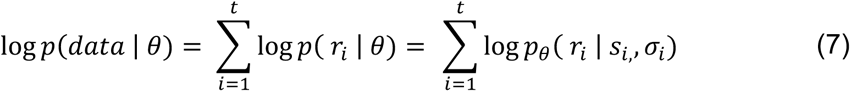

#### Model Selection

To compare candidate models, we used Akaike information criterion (*AIC*) and Bayesian information criterion (*BIC*), summed across participants, for model selection. For a single participant, *AIC* is defined as:

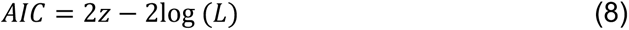

Where *z* is the number of model parameters and log(*L*) is the log likelihood of the dataset. To quantify support for each model across all participants, we summed AIC values across participants, *b,*

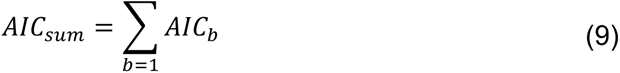

For a single participant, *BIC* is defined as:

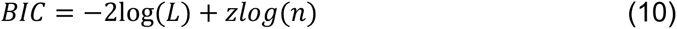

Where *z* is the number of model parameters, log(*L*) is the log likelihood of the dataset and *n* is the number of trials in the dataset. Because *BIC* is sensitive to the total sample size used to compute the likelihood, calculating the summed *BIC* for each model requires that the sample size accounts for the total number of trials across all participants. We therefore summed the likelihood terms across participants, *b*, and added to that the total number of free parameters across participants, *zB*, multiplied by the log transformed total number of trials across participants, *nB*.

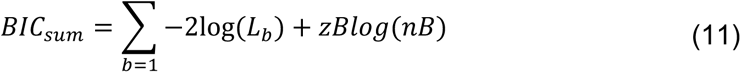

Where *B* is the total number of participants.

#### Visualisation of Model Fits

##### Non-Bayesian Models

To visualise the fit of the non-Bayesian models, we generated a synthetic dataset of model responses using the best fitting parameters for each individual. Using the experimental data for that individual, for each trial we took 100 samples from a normal distribution with mean *s_i_* (the true stimulus value) and standard deviation *σ_c_* (estimated value of *σ* for a given intensity *c*). We then compared each sample to the boundary parameters (*b_1_* − *b_3_* for the different means task and *b_1_* − *b_7_* for the different SDs task) to generate 100 category/confidence responses for that trial. We took the mean of the generated category/confidence responses to produce a single average response for that trial.

##### Bayesian Models

As with non-Bayesian models, for each trial, we took 100 samples from a normal distribution with mean *s_i_* and standard deviation *σ_$_*. We transformed boundary parameters from probability space into perceptual space (using the relevant equation from the model specification section) and compared each stimulus sample to the resulting boundaries to generate a predicted response. This approach generated 100 category/confidence responses for each trial, and we took the mean of the generated category/confidence responses.

##### Bayesian Models with Decision Noise

For Bayesian models with decision noise, we compared each stimulus sample to 100 random draws of the category/confidence boundaries. We sampled from a normal distribution with standard deviation *σ_d_* and mean *k*. Each draw was converted into perceptual space using the relevant equation and *σ_c_*. We generated 100 × 100 category/confidence responses for each trial and took the mean of the generated category/confidence responses to produce a single average response for that trial.

##### Generating Figures

For plots with stimulus value (or a transformation of stimulus value) on the horizontal axis, stimulus values were binned so that each bin consisted of the same number of trials. For visualisation, we calculated the mean category/confidence response for both the experimental data and model predictions for each bin.

## Results

### Participant Exclusion

To ensure that participants understood the task and the underlying category distributions, we excluded data from individuals whose categorisation accuracy was not significantly above chance (50%) in the highest intensity condition, using a binomial test of significance. This test was applied to each task and modality separately and included testing trials only (see Supplementary Table 1). We considered a category response as correct if it was the most probable generating category, given the stimulus value and category distributions for that task. One of the 12 participants was excluded using this criterion. A second participant was excluded for reported confusion about the task instructions, a third was excluded for not completing all sessions (2/4 sessions) and a fourth was excluded for using the highest confidence rating on every trial in one session. The target sample size (after exclusions; *N* = 8) was preregistered and was determined a priori (https://osf.io/d3y5n), which allowed full counterbalancing of experimental conditions.

### Confirming the Effect of the Chosen Stimulus Manipulations

Although our primary theoretical interest was in the model-based analysis of the data, we first checked that our chosen stimulus manipulations had the intended effects on category and confidence responses across modalities. As described below, we fit generalised linear mixed models (GLMMs) to predict category and confidence responses from the relevant stimulus features for each modality and task. In Panel A of Fig 4, we show the raw data (without GLMM predictions) for each modality and task (columns). Stimulus values, orientation for the visual tasks and frequency for the auditory tasks, are binned on the x axis so that each bin contains the same number of values. Responses, on the y axis, are coded such that value indicates confidence (1, 2, 3, 4) and sign indicates category, where category 1 responses are negative and category 2 responses are positive.

**Fig 4.**
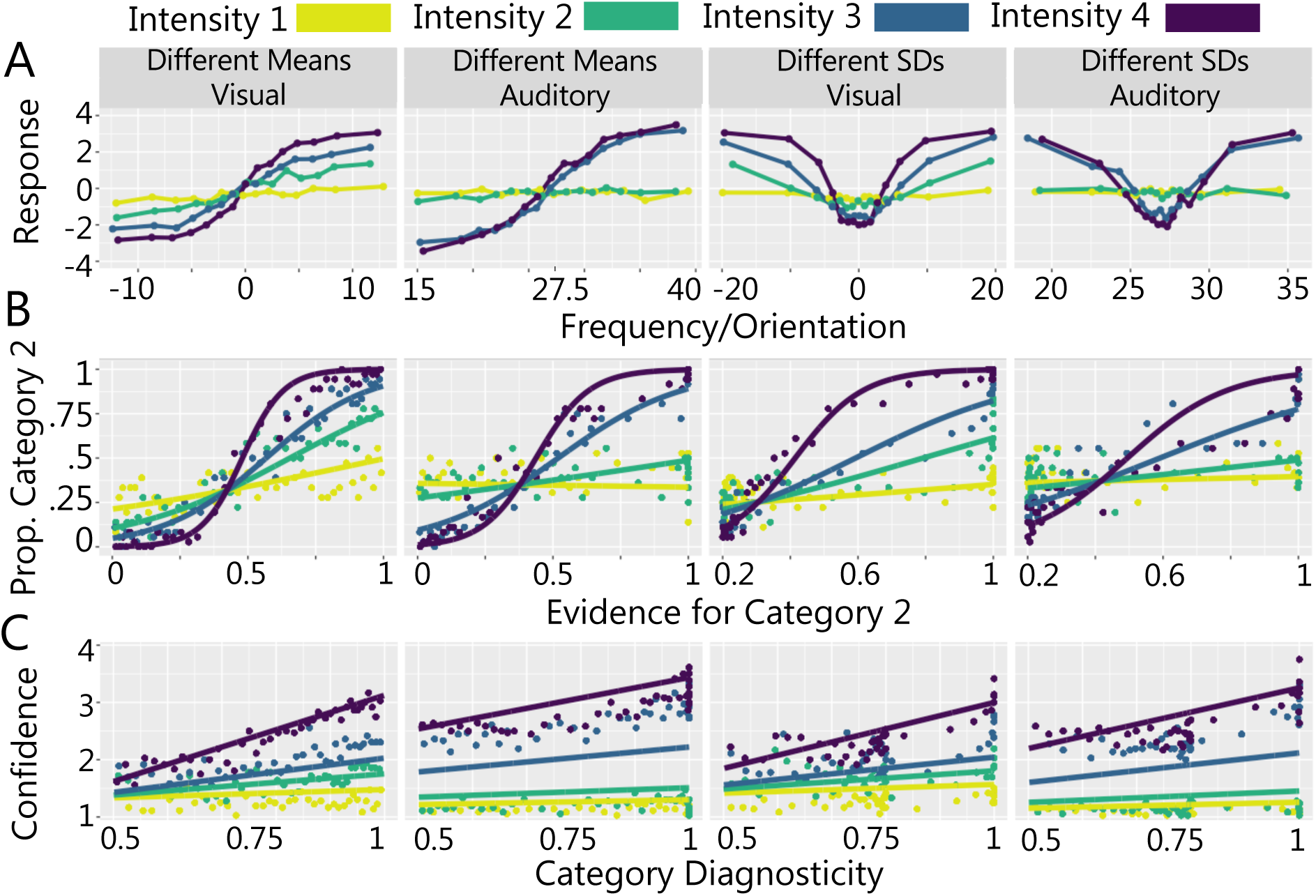
Logistic and Linear Mixed Models. (A) Mean response as a function of binned stimulus value. (B) Category decisions as a function of category evidence. Data points show binned evidence values and solid lines show logistic model predictions for the fixed effect of evidence at each intensity level. (C) Confidence as a function of category diagnosticity. Data points show binned evidence values and solid lines show linear model predictions for the fixed effect of category diagnosticity at each intensity level.

Because the expected and observed pattern of responses differed across the different means and different SDs task (see Fig 2A), we had to transform the stimulus values (see Fig 4B,C) to fit the GLMMs and compare the results across the different versions of the task.

### Confirming that the Stimulus Manipulations Affected Category Responses

To determine the effect of the stimulus manipulations on category responses, we transformed stimulus values to ‘evidence for category 2’. For each task and modality, we computed the evidence for category 2 by evaluating the probability density of the presented stimulus value for the category 2 distribution, normalised by the total probability density for both category distributions for that stimulus value,

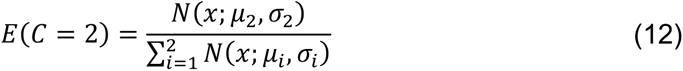

where *N* denotes the normal density function.

The result indicated evidence for category 2 for each stimulus value, which could range from 0 (stimulus could be from category 1 only) to 1 (stimulus could be from category 2 only). Note, however, that for the different SDs task evidence for category 2 never reaches 0 because the category 1 distribution falls entirely within the range of the category 2 distribution. The minimum evidence value for this task is 0.2.

We fit linear mixed models using a logistic link function to predict category response (1 or 2) from stimulus intensity and evidence for category 2, for each task and modality. The models included an intercept term, intensity, evidence and the interaction term between intensity and evidence as random effects. For each regression analysis, we report the fixed effect and 95% confidence intervals in Table 1. As this analysis was intended to provide a descriptive summary of the effect of the stimulus manipulations on responses across modalities, as opposed to a hypothesis testing exercise, we report other statistical metrics and follow-ups in Supplementary Table 2.

**Table 1.** GLMM with Stimulus Manipulations as Predictors

| GLMM Predictor |  | Tasks |  |  |  |
| --- | --- | --- | --- | --- | --- |
|  |  | Visual<br>Different<br>Means | Auditory<br>Different<br>Means | Visual<br>Different SDs | Auditory<br>Different SDs |
|  |  | Category |  |  |  |
| Intercept | Mean | -0.31 | -0.03 | -0.27 | -0.30 |
| | $CI_{lower}$ | -0.74 | -0.43 | -0.49 | -0.69 |
| | $CI_{upper}$ | 0.12 | 0.36 | -0.05 | 0.09 |
| Evidence for Category 2 | $\beta$ | 1.85 | 1.42 | 1.34 | 0.87 |
| | $CI_{lower}$ | 1.40 | 1.09 | 1.11 | 0.50 |
| | $CI_{upper}$ | 2.30 | 1.76 | 1.52 | 1.24 |
| Stimulus Intensity | $\beta$ | 0.35 | 0.61 | 0.69 | 0.22 |
| | $CI_{lower}$ | 0.12 | 0.01 | 0.37 | -0.38 |
| | $CI_{upper}$ | 0.59 | 1.23 | 0.89 | 0.82 |
| Evidence × Intensity<br>Interaction | $\beta$ | 1.45 | 1.49 | 1.13 | 0.83 |
| | $CI_{lower}$ | 0.99 | 1.07 | 0.95 | 0.47 |
| | $CI_{upper}$ | 1.88 | 1.92 | 1.29 | 1.20 |
|  |  | Confidence |  |  |  |
| Intercept | Mean | 1.82 | 1.98 | 1.88 | 1.82 |
| | $CI_{lower}$ | 1.45 | 1.81 | 1.46 | 1.66 |
| | $CI_{upper}$ | 2.18 | 2.15 | 2.30 | 1.98 |
| Category Diagnosticity | $\beta$ | 0.19 | 0.13 | 0.15 | 0.13 |
| | $CI_{lower}$ | 0.13 | 0.08 | 0.11 | 0.07 |
| | $CI_{upper}$ | 0.25 | 0.19 | 0.18 | 0.19 |
| Stimulus Intensity | $\beta$ | 0.40 | 0.72 | 0.38 | 0.62 |
| | $CI_{lower}$ | 0.25 | 0.53 | 0.26 | 0.46 |
| | $CI_{upper}$ | 0.56 | 0.92 | 0.51 | 0.79 |
| Diagnosticity × Intensity<br>Interaction | $\beta$ | 0.15 | 0.11 | 0.10 | 0.10 |
| | $CI_{lower}$ | 0.09 | 0.06 | 0.07 | 0.05 |
| | $CI_{upper}$ | 0.21 | 0.15 | 0.14 | 0.15 |

Note. CI terms refer to 95% Confidence Intervals

As expected, stimulus values that indicated increasing evidence for category 2 were positively predictive of category 2 responses in all tasks and modalities (see ‘Evidence for Category 2’ in top panel of Table 1 for coefficients). Increasing stimulus intensity was also positively predictive of category 2 responses for all tasks except the auditory different SDs task (see ‘Stimulus Intensity’ in top panel of Table 1 for coefficients). These main effects were qualified by significant two-way interactions between category 2 evidence and intensity, for all tasks and modalities (see ‘Evidence × Intensity Interaction’ in top panel of Table 1 for coefficients). As illustrated in Fig 4B, the effect of increasing category 2 evidence on category 2 responses was strongest for the highest intensity stimuli and decreased with stimulus intensity (see Supplementary Table 2 for follow up tests).

These findings demonstrate that participants understood the category distributions across tasks and modalities. Specifically, when there was an increase in the relative probability that a stimulus was sampled from category 2, participants were more likely to report that that stimulus had been sampled from category 2. This was true regardless of the specific category structure (i.e., for both the different means and different SDs tasks) and stimulus modality. The effect of category 2 evidence was modulated by stimulus intensity, specifically, contrast for the visual tasks and loudness for the auditory tasks, such that as the intensity of the stimulus was reduced, category responses were more random and less influenced by the category distributions. This effect was observed across modalities and confirmed that the chosen stimulus manipulations had the intended effect on participants’ category responses in both the visual and auditory domains. Because we were interested in quantifying the computations underlying metacognitive judgements across modalities, this analysis also provided evidence for similarities in the perceptual task requirements, allowing us to isolate any differences in metacognition itself (Lehmann et al., 2022).

### Confirming that the Stimulus Manipulations Affected Confidence

As above, the expected pattern of confidence responses differed across the tasks. To compare the effect of the stimulus manipulations on confidence across the different versions of the tasks we transformed stimulus values to ‘category diagnosticity’ prior to fitting the GLMMs. For each task and modality, we computed category diagnosticity by evaluating the probability density of the presented stimulus value for the most probable category distribution, normalised by the total probability density for both category distributions:

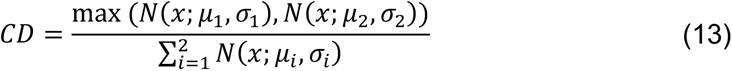

where *N* denotes the normal density function.

The result indicated category diagnosticity for each stimulus value, which could range from 0.5 (both categories are equally likely) to 1 (stimulus could be from the most probable category only). We fit linear mixed models to predict confidence from stimulus intensity and category diagnosticity, for each task and modality. The models included an intercept term, intensity, diagnosticity and the interaction term between intensity and diagnosticity as random effects. For each regression analysis, we report the fixed effect and 95% confidence intervals in Table 1.

Increasing category diagnosticity (see ‘Category Diagnosticity’ in bottom panel of Table 1 for coefficients) and stimulus intensity (see ‘Stimulus Intensity’ in bottom panel of Table 1 for coefficients) were predictive of increasing confidence in all tasks. These main effects were qualified by significant two-way interactions between category diagnosticity and intensity for all tasks (see ‘Diagnosticity × Intensity Interaction’ in bottom panel of Table 1 for coefficients). Increasing category diagnosticity was associated with increasing confidence but the strength of this effect was modulated by stimulus intensity. As illustrated in Fig 4C, the effect of increasing category diagnosticity was strongest for the highest intensity stimuli and decreased with lower stimulus intensities (see Supplementary Table 2 for follow up tests).

These results indicate that as the relative evidence for a given category increased, participants’ confidence in their chosen category increased. Taken with the results of the category GLMMs, showing that participants’ category responses corresponded with the most probable category given the value of the stimulus, these findings suggest that participants understood the category distributions and used them to accurately guide their confidence responses. The effect of the category distributions on confidence was modulated by stimulus intensity, such that as the intensity of the stimulus decreased, confidence responses were uniformly low. This effect was observed across both modalities.

### Computational Modelling

The GLMMs showed that our chosen stimulus manipulations had the intended effects on category and confidence responses across modalities. We next sought to characterise the processes underlying the category and confidence judgements using computational modelling. We evaluated the performance of popular confidence models from three classes: 1) unscaled evidence strength models, 2) scaled evidence strength models and 3) Bayesian models. To assess the similarity in the underlying computational processes, we sought to determine whether a single class of models provided the best account of responses across modalities and tasks (visual different means, auditory different means, visual different SDs, auditory different SDs). An overview of the computational mechanisms in each class and the performance of each model is provided in detail below.

#### Unscaled Evidence Strength Models: The Distance model

According to unscaled evidence strength models, confidence reports depend on the distance between the sensory measurement and the category decision criterion. These models do not independently quantify sensory uncertainty. We tested a single variation of this class of models: the distance model. In the distance model, the observer compares their internal representation of a stimulus to a set of category/confidence boundaries that do not change with stimulus intensity.

To visualise the fit of the distance model, in Fig 5 we show the model’s predicted responses (solid lines) against experimental data (square plotting symbols) as a function of standardised stimulus values (see Supplementary Fig 13 for alternative visualisations of model fits). The experimental data show a clear effect of stimulus intensity (denoted by different colours) on category and confidence responses. The distance model, however, has no mechanism to account for modulation by stimulus intensity. Unlike scaled evidence strength models, the boundary positions are not dependent on the estimate of measurement noise for a given intensity, so the model is forced to settle on a set of boundaries that best captures responses across all intensities. As a result, the model overpredicts confidence for low intensities and underpredicts confidence for high intensities. This pattern of over- and under- prediction is seen across modalities and tasks. Overall, the distance model did not perform well, suggesting that a model in which confidence depends entirely on the distance from the category decision criterion provides a poor account of observed responses in both the visual and auditory tasks.

**Fig 5.**
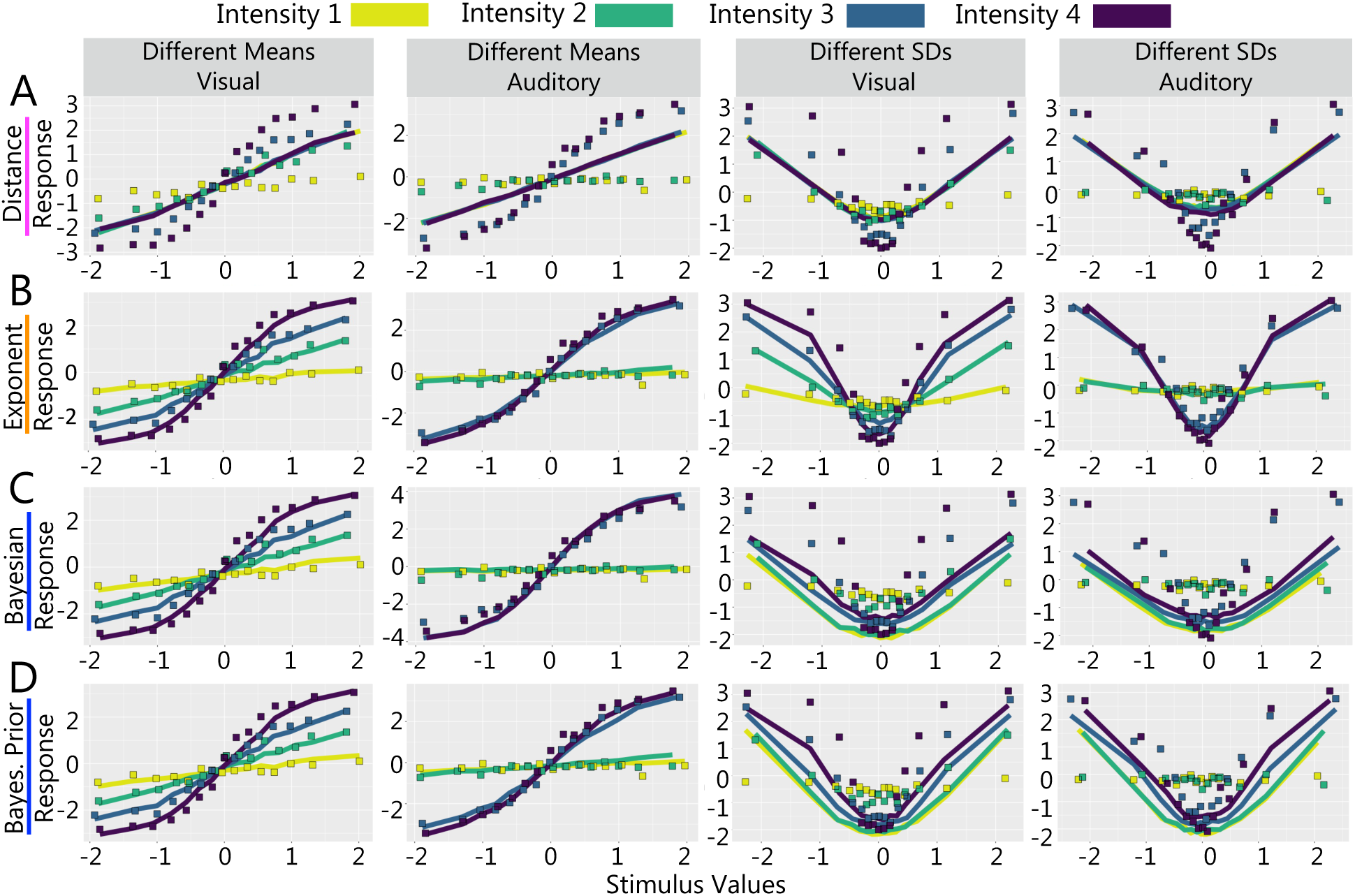
Model Predictions and Experimental Data. Columns show different versions of the task. Rows show model fits for (A) the distance model (unscaled evidence strength class; pink), (B) the free-exponent model (scaled evidence strength class; orange), (C) the log posterior probability ratio model (Bayesian class; blue) and (D) the log posterior probability ratio model with free category distribution parameters (Bayesian class; blue).

#### Scaled Evidence Strength Models: Linear, Quadratic & Exponent Models

According to scaled evidence strength models, confidence reports depend on the distance between the observer’s internal representation of the stimulus and sensory uncertainty dependent choice and confidence boundaries. We tested three variations of this class of models which used different scaling rules: the linear model, the quadratic model and the free-exponent model.

Each model allows for a point estimate of measurement noise to expand/contract the category/confidence boundaries and thereby account for the changes in responding across stimulus intensities seen in the experimental data. We show the predictions for the free exponent model in Fig 5B. See Supplementary Fig 14 for linear model predictions and Supplementary Fig 15 for quadratic model predictions. All members of this model class fit the data well and the differences in the predictions across models are subtle.

The additional free parameter in the free-exponent model allowed for the changes in boundary positions across intensities to occupy values between and beyond linearly- or quadratically-spaced values. Thus, the free-exponent model can account for a wider range of sensory uncertainty effects across participants. This additional flexibility was best able to capture the experimental data on average and thus, the model has the lowest summed *AIC* and *BIC* scores across tasks and modalities (with the exception of summed *BIC* for the linear model in the auditory different means task; see Table 2). See Supplementary Tables 9 and 10 for best fitting parameters.

**Table 2.** Model Fits

| Model Class | Model | Visual |  | Preferred Model<sup>a</sup> | Auditory |  | Preferred Model<sup>a</sup> |
| --- | --- | --- | --- | --- | --- | --- | --- |
| | | $AIC_{sum}$ | $BIC_{sum}$ | | $AIC_{sum}$ | $BIC_{sum}$ | |
| Different Means Task |  |  |  |  |  |  |  |
| Unscaled ES | Distance | 16666.53 | 17092.68 | 0 | 18909.84 | 19336.00 | 0 |
| Scaled ES | Linear | 14183.77 | 14823.01 | 1 | 13011.35 | **13650.59** | 5 |
| Scaled ES | Quadratic | 14200.41 | 14839.64 | 1 | 13033.60 | 13672.84 | 3 |
| Scaled ES | Free-Exponent | 14144.17 | 14836.67 | 0 | **13001.97** | 13694.47 | 0 |
| Scaled ES | Free-Exponent + ODN | **13745.56** | **14687.33** | 4 | - | - | - |
| Bayesian | LPPR | 14382.99 | 14809.15 | 1 | 14054.42 | 14480.57 | 0 |
| Bayesian | LPPR + ODN | 14344.52 | 14823.95 | 1 | - | - | - |
| Bayesian | LPPR + D noise | 14373.15 | 14852.58 | 0 | 13995.73 | 14475.16 | 0 |
| Bayesian | LPPR: Free Prior | 14385.80 | 14918.50 | 0 | 14097.59 | 14630.28 | 0 |
| Different SDs Task |  |  |  |  |  |  |  |
| Unscaled ES | Distance | 17125.22 | 17711.18 | 0 | 18414.05 | 19000.02 | 0 |
| Scaled ES | Linear | 14661.52 | 15620.37 | 5 | 13400.11 | 14358.96 | 1 |
| Scaled ES | Quadratic | 14737.16 | 15696.01 | 1 | 13370.00 | 14328.85 | 1 |
| Scaled ES | Free-Exponent | 14592.14 | **15604.27** | 2 | **13016.11** | **14028.23** | 6 |
| Scaled ES | Free-Exponent + ODN | **14478.89** | 15824.28 | 0 | - | - | - |
| Bayesian | LPPR | 18638.13 | 19224.10 | 0 | 20096.12 | 20682.09 | 0 |
| Bayesian | LPPR + ODN | 18761.58 | 19400.81 | 0 | - | - | - |
| Bayesian | LPPR + D noise | 17123.45 | 17762.68 | 0 | 17817.43 | 18456.66 | 0 |
| Bayesian | LPPR: Free Prior | 18509.15 | 19201.65 | 1 | 21209.72 | 21902.22 | 0 |

*Note.* ES in model class labels refers to Evidence Strength. LPPR refers to the log posterior probability ratio models, ODN refers to models which include an orientation dependent noise parameter and D noise refers to models with a decision noise parameter. <sup>a</sup>The number of participants for whom the model was the best fitting model. Where there was inconsistency between $AIC$ and $BIC$ for the best fitting model, we chose the model with the lowest $BIC$ .

As shown in Table 2, adding an orientation-dependent noise parameter for modelling performance in the visual task provides an improvement in the performance of the free-exponent model in the different means task and a marginal improvement in the different SDs task. Including the orientation-dependent noise parameter, however, did not qualitatively change the visualisation of the model predictions or the conclusions (that is, which class of model was the best performing across modalities and tasks).

Overall, the scaled evidence strength models were on average the best performing models across tasks and modalities. This finding suggests that a model in which participants use sensory uncertainty dependent choice and confidence boundaries provides a good account of observed responses in both the visual and auditory tasks.

#### Bayesian Models

According to Bayesian models of confidence, observers optimally combine sensory information with prior knowledge of the generative model to form a posterior belief about the nature of the stimulus. We tested several variants of this class of models.

The Bayesian models have fewer parameters than the scaled evidence strength models because the boundaries have the same positions in probability space across intensity levels. The estimates of measurement noise at each intensity level allow these boundaries to shift in perceptual space. As shown in Fig 5C/D, this allows the model to account for different patterns of responding across intensities. This mechanism appears sufficient for the different means task where overall the Bayesian models fit the data well in both modalities—though not as well as the scaled evidence strength models (see Fig 6). In the different SDs task, however, the Bayesian models produced qualitative mispredictions of category and confidence data. We discuss the performance of each model in the Bayesian class in turn below (with emphasis on its performance in the different SDs task).

**Fig 6.**
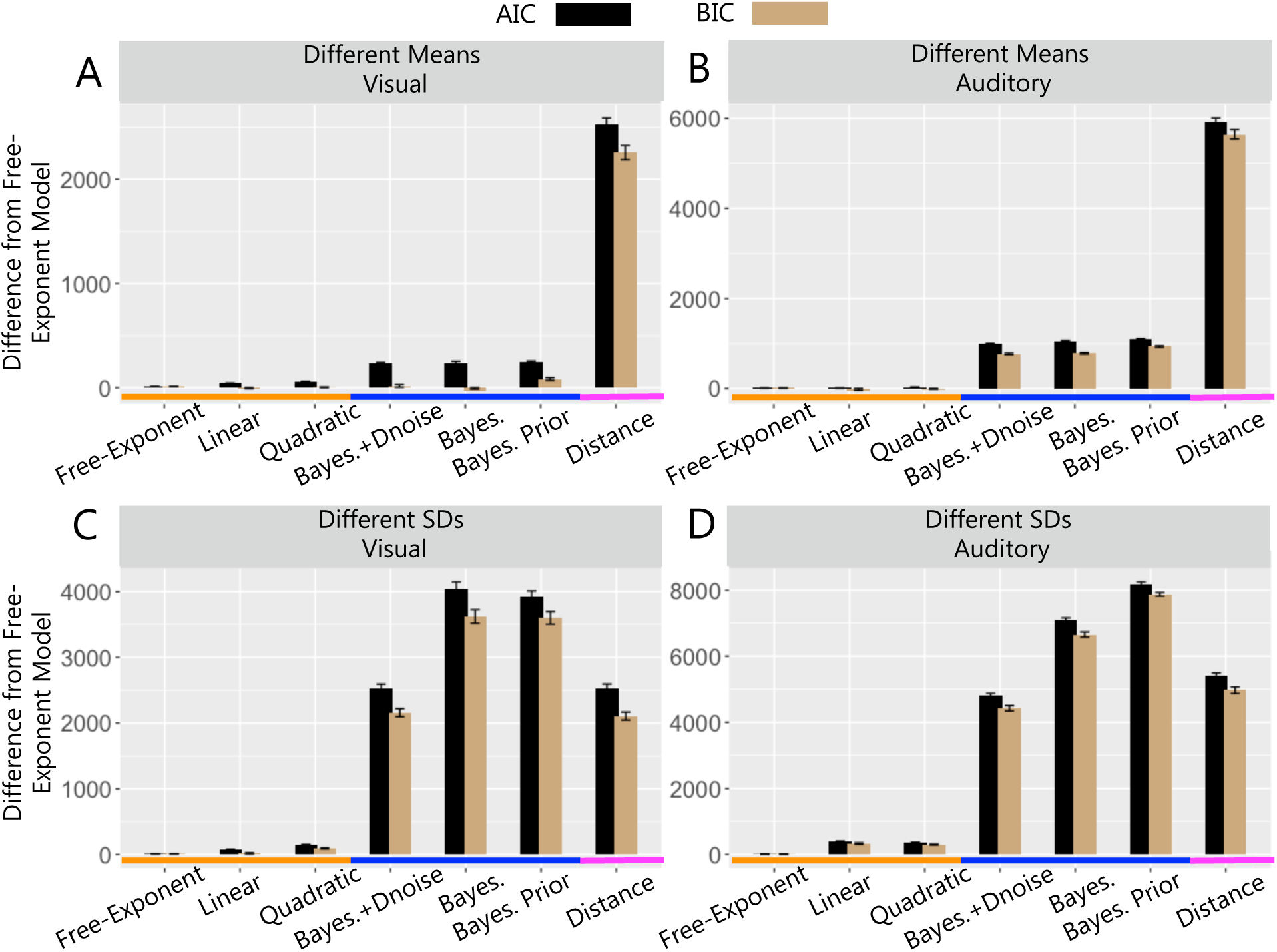
Model Comparison. Difference in summed *AIC* (black) and *BIC* (brown) between the free-exponent model and other models from 3 classes: unscaled evidence strength model (pink; the distance model), scaled evidence strength models (orange; free-exponent, linear and quadratic models) and Bayesian models (blue; the log posterior probability ratio model labelled as ‘Bayes.’; the log posterior probability ratio model with decision noise labelled as ‘Bayes.+Dnoise’; and the log posterior probability ratio model with free category distribution parameters labelled as ‘Bayes. Prior’). (A) Different means visual task, (B) different means auditory task, (C) different SDs visual task and (D) different SDs auditory task.

##### Log Posterior Probability Ratio

In the log posterior probability ratio model, we assume that the observer compares the log posterior probability ratio of the stimulus measurement to a set of category and confidence boundaries in log posterior probability ratio space. In the different SDs task, the main failure of this model is that it overpredicts confidence at lower intensity levels across modalities (see Supplementary Fig 17). Even though the boundary positions are sensitive to the estimates of sigma and the model is able to generate different predictions at each intensity level, the boundary parameters themselves are shared across all intensity levels such that the fit to low-intensity trials is constrained by high-intensity trials. This constraint does not appear to allow sufficient flexibility in the model to account for the distinct patterns of responses across intensity levels. We tested several variations of the log posterior probability ratio model which relaxed the assumptions of the Bayesian framework.

##### Log Posterior Probability Ratio with Orientation Dependent Noise

Adding an orientation-dependent noise parameter improves the fit of the log posterior probability ratio model to visual modality data in the different means task but not in the different SDs task. The predictions of this model still show the same systematic deviations from the data as the log posterior probability ratio model in that it overpredicts confidence at lower intensity levels across modalities (see Supplementary Fig 18).

##### Log Posterior Probability Ratio with Decision Noise

Adding a Gaussian noise term on the calculation of the log posterior probability ratio improved the fit of the log posterior probability ratio model in both tasks and modalities. Including the decision noise parameter shifts the best-fitting boundary parameters away from the means of the category distributions. As a result, the model predicts more low confidence, category 1 responses at the lower intensity levels for all stimulus values (see Supplementary Fig 19). This aligns with the data better in terms of confidence but overpredicts category 1 responses as the experimental data show roughly equal proportions of category 1 and category 2 responses for a wide range of stimulus values at low intensities (i.e., participants are likely *guessing* a category more often). As a result of the shift in boundary positions away from the centre of the category distributions across all intensity levels, the model underpredicts confidence for stimulus values close to the mean of the category distributions and underpredicts category 2 responses for extreme stimulus values at the higher intensity levels. In the different SDs task, the log posterior probability ratio model with decision noise is the best performing of the Bayesian models in both modalities. In the different means task, it is also the best performing Bayesian model in the auditory modality. In the visual modality, the log posterior probability ratio model with decision noise is outperformed by the log posterior probability ratio model with orientation-dependent noise.

##### Log Posterior Probability Ratio with Free Category Distribution Parameters

The log posterior probability ratio model assumes that the observer knows the true values of the means and standard deviations of the category distributions. We tested a model in which the observer has imperfect knowledge of the parameters of the generative model, and we estimated these values as free parameters instead. Like the standard log posterior probability ratio model, this model overpredicts confidence at lower intensities and underpredicts confidence for stimulus values at the centre of the distributions at the highest intensity (see Supplementary Fig 20). This misprediction is seen across modalities. The model does, however, do a better job of matching confidence and proportion of category 2 responses at the more extreme values at higher intensities (relative to the log posterior probability ratio model with and without decision noise).

In the different means task, the best-fitting category distribution parameters were similar to the true category distribution parameters (see Supplementary Table 9 for parameters). In the different SDs task, the best-fitting category distribution parameters had a consistently smaller standard deviation for the category 1 distribution and a larger standard deviation for the category 2 distribution in both modalities (see Supplementary Table 10 for parameters). The performance of this model demonstrates that even with a data-informed subjective prior, the Bayesian model class still produces qualitative mispredictions of category and confidence data in both the visual and auditory tasks. Overall, it appears that regardless of the flexibility of the model, observed responses in the visual and auditory tasks are not fully consistent with a Bayesian framework in either modality.

#### Consistency Across Participants

##### Consistency in Best Performing Model Class

The analyses described so far characterise how well each model captured the general patterns of responding across participants. As models were fit to data from each individual, however, we were also able to examine the best-fitting model for each participant. If a single class of models reliably characterises the computational processes underlying participants’ category and confidence decisions collectively, we would expect that data from individual participants should also be best characterised by the same class of models across modalities and tasks. If, however, a single class of models was not able to reliably characterise the underlying computational processes, we would expect to see variation in the performance of the model classes across individuals. Although our goal was not to investigate individual differences per se, which would require a larger sample size, we can nonetheless examine the consistency in model fits across participants.

As shown in Table 2, in the auditory modality the scaled evidence strength models performed best for all participants. In the visual modality, the scaled evidence strength models were the best performing for 75% of participants in the different means task (the Bayesian class was the best performing model for the remaining 25%) and for 87.5% in the different SDs task (the Bayesian class was the best performing model for the remaining 12.5%). This analysis reveals that even at the individual level, there was a preference for the scaled evidence strength class across modalities and tasks, consistent with the summed *AIC* and *BIC* scores which are the lowest for the scaled evidence strength models, as described above.

##### Consistency in Best Performing Model

While the main motivation of the model-based comparison was to focus the analysis at the level of model classes, to further understand the specific implementation of the scaled evidence strength models, we also examined differences at the individual model level. We compared the best performing model for each participant and found that within the scaled evidence strength class, there were notable differences in model performance.

In the auditory different means task, either the linear model (62.5% of participants) or quadratic model (37.5% of participants) performed best for individuals, although the free-exponent model was the best performing model on average. For the visual different means task, by contrast, either the free-exponent model (with orientation dependent noise; 50% of participants), the linear model (12.5% of participants) or the quadratic model (12.5% of participants) was the best performing model for participants best characterised by a scaled evidence strength class model. In the different SDs task in the visual modality, either the linear (62.5%), quadratic (12.5%) or free-exponent (12.5%) performed best and in the auditory modality, the free-exponent model was preferred (87.5%; remaining 12.5% for quadratic and 12.5% for linear). Therefore, although the scaled evidence strength class consistently outperformed the other model classes, a specific scaling rule within that class (linear, quadratic or other exponent) does not seem to apply consistently across different versions of the task, across modalities or across participants.

### Parameter Settings Across Modalities

In the analyses described thus far we sought to determine whether a single class of metacognitive models could account for choice and confidence judgements across the visual and auditory modalities. Consistent with this, we found that the scaled evidence strength models were the best performing across both the different means and different SDs task in both modalities. We found consistent evidence for this class of models both at the group level, where the models were best able to capture the main patterns of responding across participants, and at the individual participant level, where data from most individual participants were best accounted for by the same class of models across tasks and modalities. Given the consistency in evidence for a single class of models, we next sought to determine the extent to which this scaled evidence strength algorithm required different parameter settings to accommodate responding across modalities.

If the scaled evidence strength algorithm operates in a “one size fits all” configuration and does not require tuning/parameterisation specific to each modality, a single set of parameters would be sufficient to account for category and confidence responses across modalities. Alternatively, if the algorithm is tuned/parameterised specifically in each modality, different sets of parameters would be required to account for responding across modalities. To address this question, rather than fit the model to each modality separately we fit the free-exponent model to data from both modalities simultaneously and formulated several levels of flexibility in the parameter settings of the free exponent model. To better visualise the fit of these models, in Fig 7 we show model predictions (solid lines and circular plotting symbols) and experimental data (square data plotting symbols) as a function of standardised stimulus value (x axes) and confidence (y axes) for the least and most flexible models.

**Fig 7.**
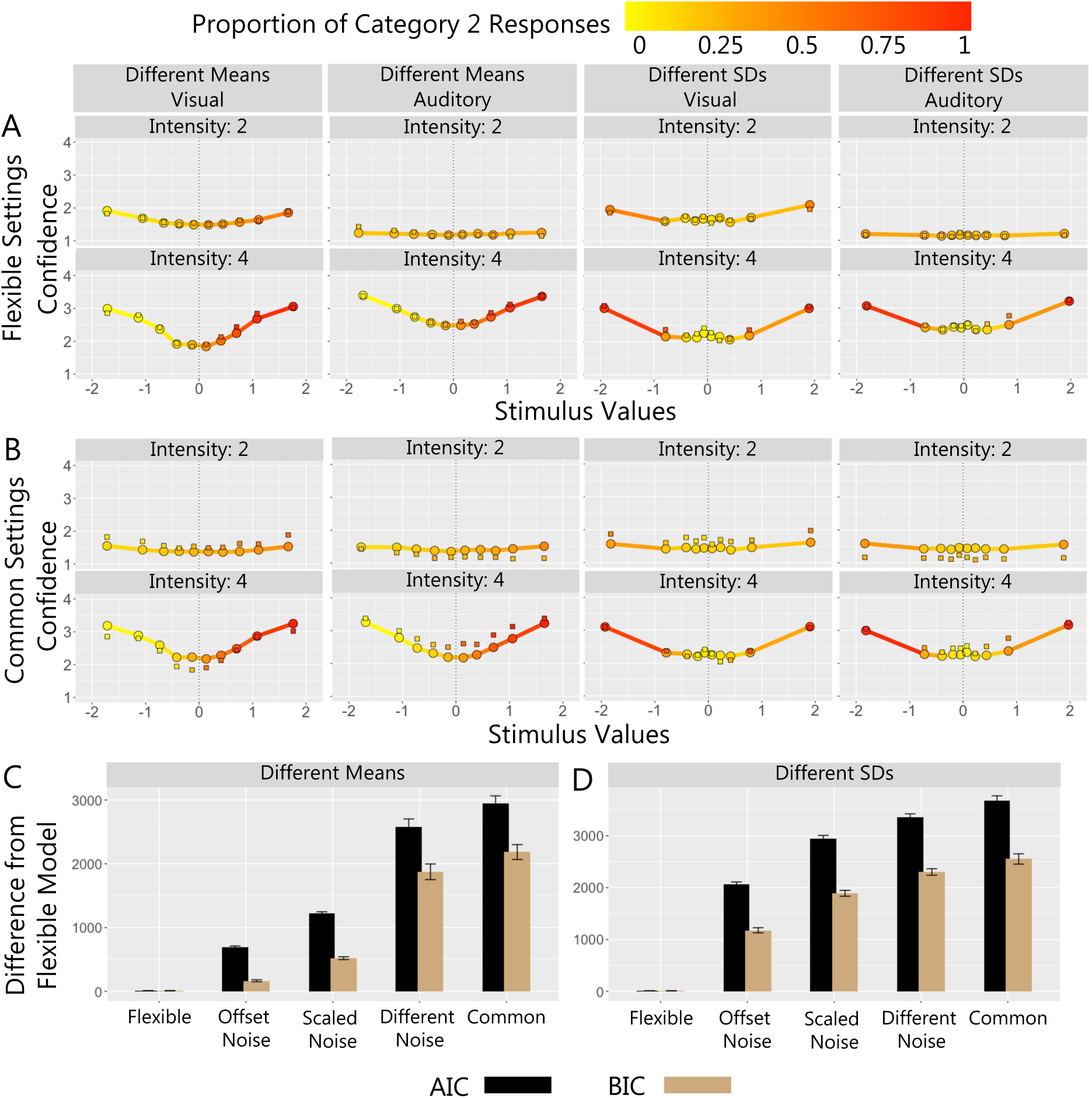
Parameter Settings Across Modalities. (A) and (B) show mean confidence and proportion of category 2 responses (colour scale) for binned stimulus values. Square plotting symbols show means for experimental data and solid lines and circular plotting symbols show means for (A) flexible settings model predictions and (B) common settings model predictions. Columns show different versions of the task and rows show different intensity levels. For clarity the plots include results for intensity levels 2 and 4 only. Difference in summed *AIC* (black) and *BIC* (brown) between the flexible settings model and the other models with different parameter settings in (C) the different means task and (D) different SDs task.

Fig 7B shows the common settings model, the least flexible of the models, in which a single set of parameters was fit to data from both modalities. The model predictions capture both the category and confidence data well. The common settings model, however, was outperformed by the more flexible models (see Table 3). The common boundary settings (scaled noise) model was marginally more flexible such that an additional free parameter was used to scale the *σ* parameters in the auditory modality relative to the visual modality. By comparison, the common boundary settings (offset noise) model had the same number of parameters, but the additional free parameter added a constant to the sigma parameters in the auditory modality relative to the visual modality. The offset noise model performed worse than the scaled noise model (Fig 7C, D). The different noise settings model, where sigma was a free parameter for each intensity level in each modality, performed better than either of the common boundary settings models. As shown in Table 3, the flexible settings model, where all parameters in the standard free-exponent model were freely estimated in each modality, was the best performing model in both the different means (see Fig 7C) and the different SDs tasks (see Fig 7D). This was true at the individual participant level and in terms of summed *AIC* and *BIC* for the group. These results show that adding flexibility in the parameter settings across modalities consistently improved the performance of the model (see Fig 7C, D). This finding suggests that although the scaled evidence strength algorithm seems to provide the best account of confidence judgements across modalities, the process is tuned specifically in each modality and does not operate in a “one size fits all” configuration.

**Table 3.**
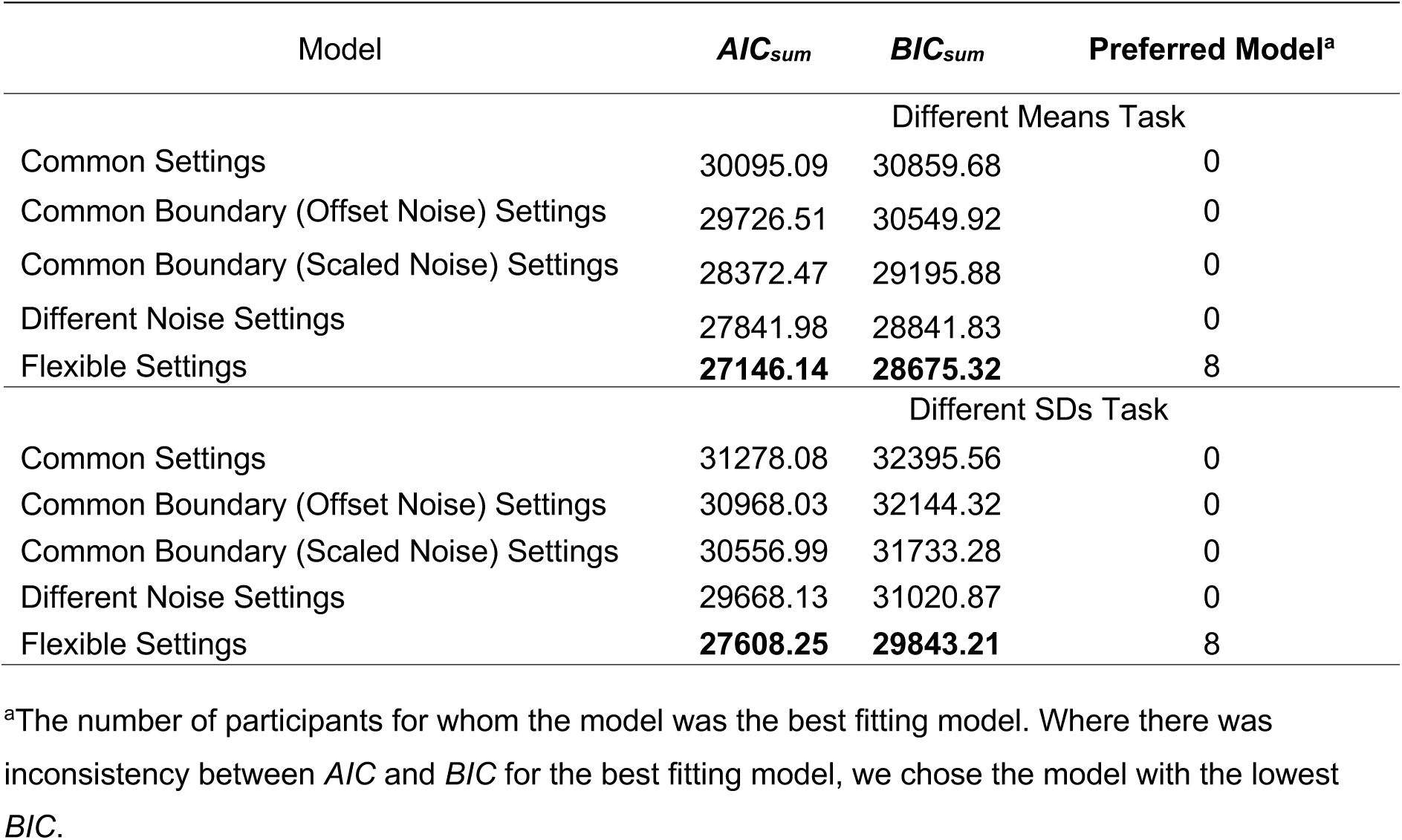
Parameter Settings Across Modalities

## Discussion

This study investigated whether the computations involved in metacognitive judgements generalise across sensory modalities. Participants completed auditory and visual versions of a categorisation task where evidence strength and sensory uncertainty were varied in each modality. We used computational modelling to quantify the mapping of evidence strength and sensory uncertainty to confidence, evaluating the performance of three distinct classes of confidence models. We found that a single class of models provided the best account of category and confidence judgements in both the visual and auditory domain but that the parameters governing this process are tuned to different values across modalities. Here we discuss how our model comparison results provide insight into the computations underlying metacognitive judgements across modalities and the extent to which these computations operate in a modality-independent or modality-specific manner.

### The Computations Underlying Metacognitive Judgements Across Modalities

#### The Importance of Accounting for Sensory Uncertainty

The unscaled evidence strength models, in which confidence depends entirely on the distance between the most likely value of the stimulus and the category decision criterion, performed consistently worse than the other model classes across both the visual and auditory categorisation tasks. Unlike the other model classes, the unscaled evidence strength class does not independently quantify sensory uncertainty, such that a given distance from the category criterion is always associated with the same level of confidence (Kepecs et al., 2008; Komura et al., 2013; Rahnev et al., 2011). The poor performance of the unscaled evidence strength class suggests that in mixed difficulty designs, observers modulate their confidence by the amount of sensory uncertainty in their internal representation of the stimulus in both the visual and auditory domain. The role of sensory uncertainty in adjusting response criteria has been demonstrated previously in the visual modality (Adler & Ma, 2018; Denison et al., 2018; Locke et al., 2022; Qamar et al., 2013) but has not been demonstrated in the auditory modality. In addition to providing evidence that observers take sensory uncertainty into account across modalities, our study also provides insight into the extent to which these processes are consistent with Bayesian or non-Bayesian accounts of confidence.

#### Implications for Bayesian Models of Confidence

Although observers take sensory uncertainty into account, they do not appear to do so in a way that is consistent with Bayesian models of confidence in either modality. The Bayesian model class performed poorly relative to the scaled evidence strength class, particularly in the different SDs task. The Bayesian class of models assumes that observers use knowledge of the level of sensory uncertainty associated with the measurement of the stimulus and (sometimes veridical) knowledge of the generating category distributions to compute a log posterior probability ratio for category membership. This value is then compared with a set of thresholds in posterior probability ratio space to determine confidence. Although a small proportion of participants’ responses were best accounted for by this model—but only in the visual modality—we found that this class of models was outperformed by the scaled evidence strength class, both for the group as a whole and for the majority of individual participants.

The relatively poor performance of the Bayesian models does not seem to be due to lack of flexibility. The Bayesian models continued to perform poorly even after allowing for a noise term in the calculation of the log posterior probability ratio or inaccurate subjective beliefs about the generative model (i.e., freely estimated category distribution parameters). Therefore, we think it is unlikely that confidence judgments in the visual and auditory modalities are generated by a Bayesian computational process.

While the poor performance of the Bayesian models is in disagreement with some studies (Aitchison et al., 2015; Hangya et al., 2016; Li & Ma, 2020; Navajas et al., 2017; Sanders et al., 2016), it is consistent with others (Adler & Ma, 2018; Aitchison et al., 2015; Bertana et al., 2021; Denison et al., 2018; Lisi et al., 2021; Locke et al., 2022). Our study extends existing research on Bayesian models of confidence in two ways. First, most previous studies have focused exclusively on the visual domain. Here we compared Bayesian and non-Bayesian accounts of confidence in both the visual and auditory domains, thus showing converging evidence for non-Bayesian metacognitive computations in a broader range of sensory contexts. Second, most previous studies have used more simple task structures (Qamar et al., 2013). Here we found that the Bayesian class did not perform as poorly when there was a single clearly defined category boundary, as in the different means task. In the different SDs task, however, where the categories overlapped to a greater extent and optimal categorisation required observers to consider the variability of both categories, the Bayesian models performed much worse. This suggests that the support for Bayesian computations in previous research may be partly attributed to low task complexity. Because the computations involved in confidence generation become more complex as the task structure becomes more complex (i.e., tasks where there is a high overlap of exemplars from different categories), deviations between human behaviour and predictions from Bayesian principles may become more pronounced. Future research evaluating Bayesian models should rely more on complex category structures to assess human behaviour relative to Bayesian optimality.

#### Non-Bayesian Sensory Uncertainty Dependent Computations

The scaled evidence strength class of models, in which positioning of the confidence and choice criteria depend on a point estimate of sensory uncertainty but does not involve the computation of posterior category probabilities, provided the best account of category and confidence responses across modalities (Adler & Ma, 2018; Baranski & Petrusic, 1994; Barthelme & Mamassian, 2010; Locke et al., 2022). We found consistent evidence for this class of models both at the group level, where the models were best able to capture the main patterns of behaviour across participants in both modalities and tasks, and at the individual participant level, where the same class of models was preferred for the majority of individuals across tasks and modalities. Using response criteria that depend on an estimate of sensory uncertainty allows observers to avoid assigning high confidence to highly variable stimuli (Locke et al., 2022; Zylberberg et al., 2014) but does so using algorithms that are less computationally expensive than Bayesian alternatives (Adler & Ma, 2018).

##### The Use of a Common Scaling Factor for Sensory Uncertainty Across Tasks and Modalities

Although we found that observers use sensory uncertainty-dependent boundaries for category and confidence judgements, the specific way sensory uncertainty scales the boundaries was not universal across modalities or tasks. We found individual differences in preferences for the different models within the scaled evidence strength class (i.e., the linear, quadratic, and free-exponent models) across modalities, tasks and individuals. These results are consistent with previous work in the visual domain which reports heterogeneity in confidence modelling, suggesting that there is no universal formula for taking sensory uncertainty into account when judging confidence (Adler & Ma, 2018; Aitchison et al., 2015; Bertana et al., 2021; Li & Ma, 2020; Lisi et al., 2021; Locke et al., 2022). It is likely that the scaling rule used to update confidence criteria reflects more contextual/task-specific factors, such as task difficulty or stimulus familiarity, rather than reflecting qualitative differences in the computational processes used by individuals within or across modalities. From a measurement perspective, this supports the case for freely estimating the scale factor, as it potentially provides a better assessment of how people’s confidence judgments are affected by sensory uncertainty. Future research should use this approach to investigate what contextual factors moderate different strategies (and scale factors) for adjusting response criteria by an estimate of sensory uncertainty.

### Modality-Independent or Modality-Specific Metacognition

Our results suggest that the general algorithm used for relating choice and confidence is the same across modalities. We did not find, however, that this process was completely ‘modality-independent’. We sought to determine the extent to which the scaled evidence strength algorithm operated in a “one size fits all” configuration or whether it required different parameter settings to accommodate responding across modalities. We fit the free-exponent model to data from both modalities simultaneously and formulated several levels of flexibility in the parameter settings of the model. The most flexible version of the model allowed the parameter values to vary across modalities, and the less flexible versions constrained parameters to be the same across modalities. We found that the most flexible model, in which all parameters were estimated independently for each modality, was the best performing model both at the individual level and the group level. Therefore, although the scaled evidence strength algorithm provided the best account of category and confidence judgements across both tasks and modalities, it is likely this process is tuned/parameterised specifically in each sensory domain. The existence of a general confidence process which may be tuned specifically to task requirements is consistent with studies finding both ‘domain general’ and ‘domain specific’ components of metacognition (Faivre et al., 2018; Lehmann et al., 2022; Masset et al., 2020; Mazancieux et al., 2020; Morales et al., 2018; Rouault et al., 2018). These studies suggest that generic confidence signals are combined with domain-specific information to fine-tune decision making (Donoso et al., 2014; Morales et al., 2018).

The existence of a general set of computations for confidence which are fine-tuned within a domain may be particularly useful when making comparative judgements across modalities or making decisions about multidimensional stimuli. For example, where an observer has to compare their confidence in a visual decision to an auditory decision or an observer has to integrate uncertainty from both the visual and auditory dimensions of a stimulus to assess their confidence. Having a common currency for confidence facilitates the integration and comparison of information across dimensions, which might maximise processing efficiency (de Gardelle et al., 2016; de Gardelle & Mamassian, 2014). We did not require participants to combine sensory information across modalities in our tasks. Future research should investigate the underlying metacognitive computations when participants are required to do so.

## Conclusions

Our study provides strong evidence that of the existing set of popular confidence models, the scaled evidence strength class provides the best account of confidence in both our visual and auditory categorisation tasks. Our model comparison results provide insight into the metacognitive computations used across modalities and the extent to which these processes operate in a modality-independent configuration. Our study paves the way for investigating how these computations might differ depending on other contextual factors or for tasks in which participants are required to explicitly compare or integrate sensory information across modalities.

## Acknowledgements

We would like to thank Dr David Lloyd and Associate Professor Wayne Wilson for their assistance with headphone calibration. We would also like to thank Dr William Adler for helpful comments and feedback about the model fitting. JBM was supported by a National Health and Medical Research Council (Australia) Investigator Grant (GNT2010141). WJH was supported by an Australian Research Council Discovery Early Career Researcher Award (DE190100136).

## Contributions

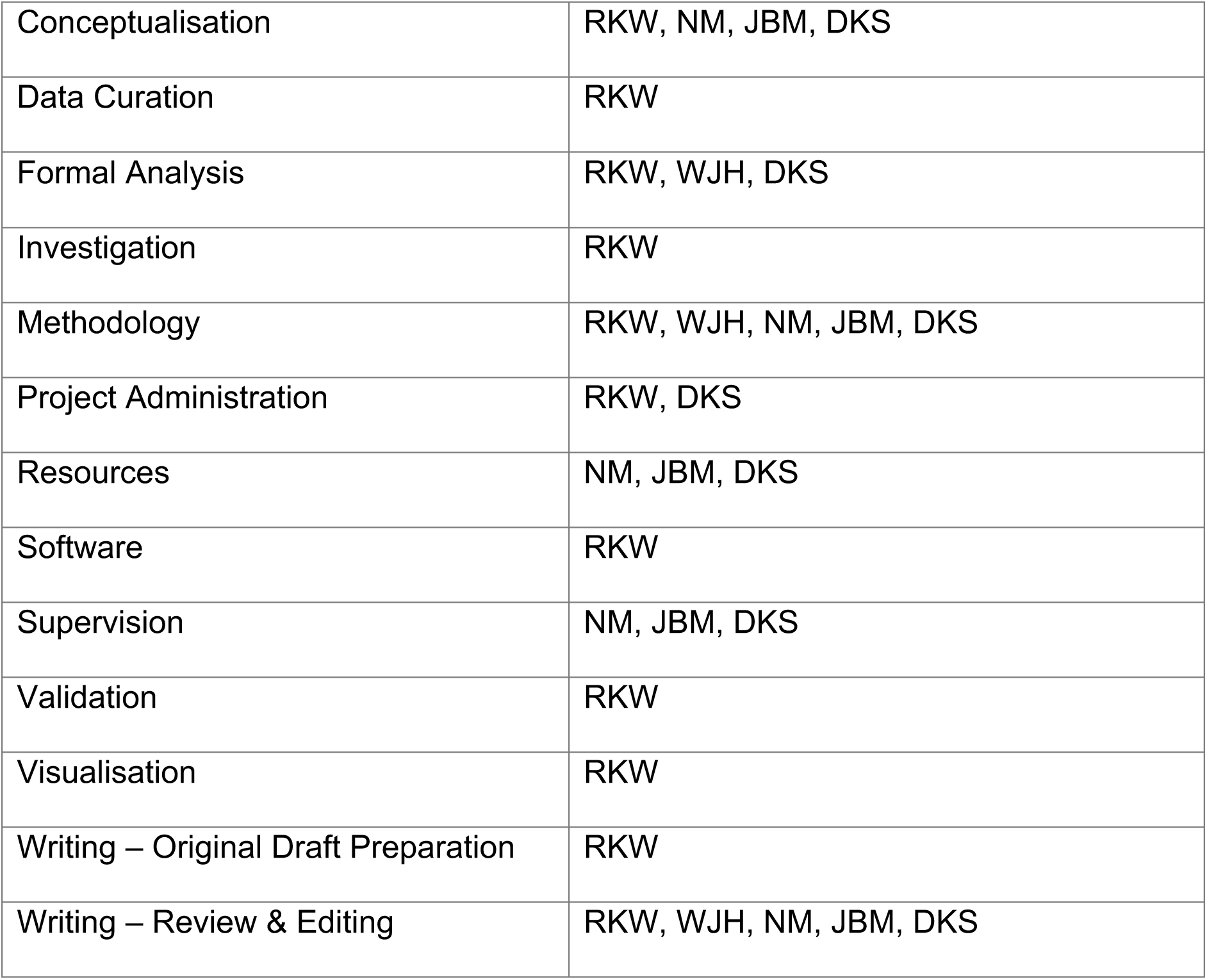

## Supplementary Information

### Headphone Calibration

All auditory calibration was done using Brüel & Kjær equipment (Brüel & Kjær, Denmark). Pure tones were generated and adjusted using Psychophysics Toolbox extensions for MATLAB on the Dell Precision T1700 with an ASIO4ALL sound driver and a sampling rate of 28 kHz to present a flat frequency response between 0.5 and 5 kHz. The flat frequency response was set by coupling the earphone of the Sennheiser HD 202 unit to a Type 5152 artificial ear with a DB0843 adaptor. The artificial ear contained a type 4144 1-inch pressure-field microphone connected to a Type 2250B handheld analyzer (a Class 1 sound level meter under AS/NZS IEC 61672-1 [Standards Australia, 2019]). The analyzer was calibrated yearly by the Brüel & Kjær laboratories in Sydney, Australia, and its status was checked before and after use by coupling it to a Type 4231 sound level calibrator. The adjustments required to present a flat frequency response output between 0.5 and 5 kHz were determined for each earphone of the Sennheiser HD 202 by presenting a 0.5 kHz tone at 85.0 dB SPL, as measured by the system described above. Tones at 0.75 kHz, 1 kHz, 1.5 kHz, 2 kHz, 3 kHz, 4 kHz and 5 kHz were then played in turn and the required adjustments in output level were determined so that each tone presented within the range of 84.0 to 86.0 dB SPL (within 1.0 dB of 85 dB SPL). The consistency of these adjustments were then checked for output levels at half the sound pressure (an output reduction of 6 dB SPL) and one quarter the sound pressure (an output reduction of 12 dB SPL).

**Supplementary Table 1.**
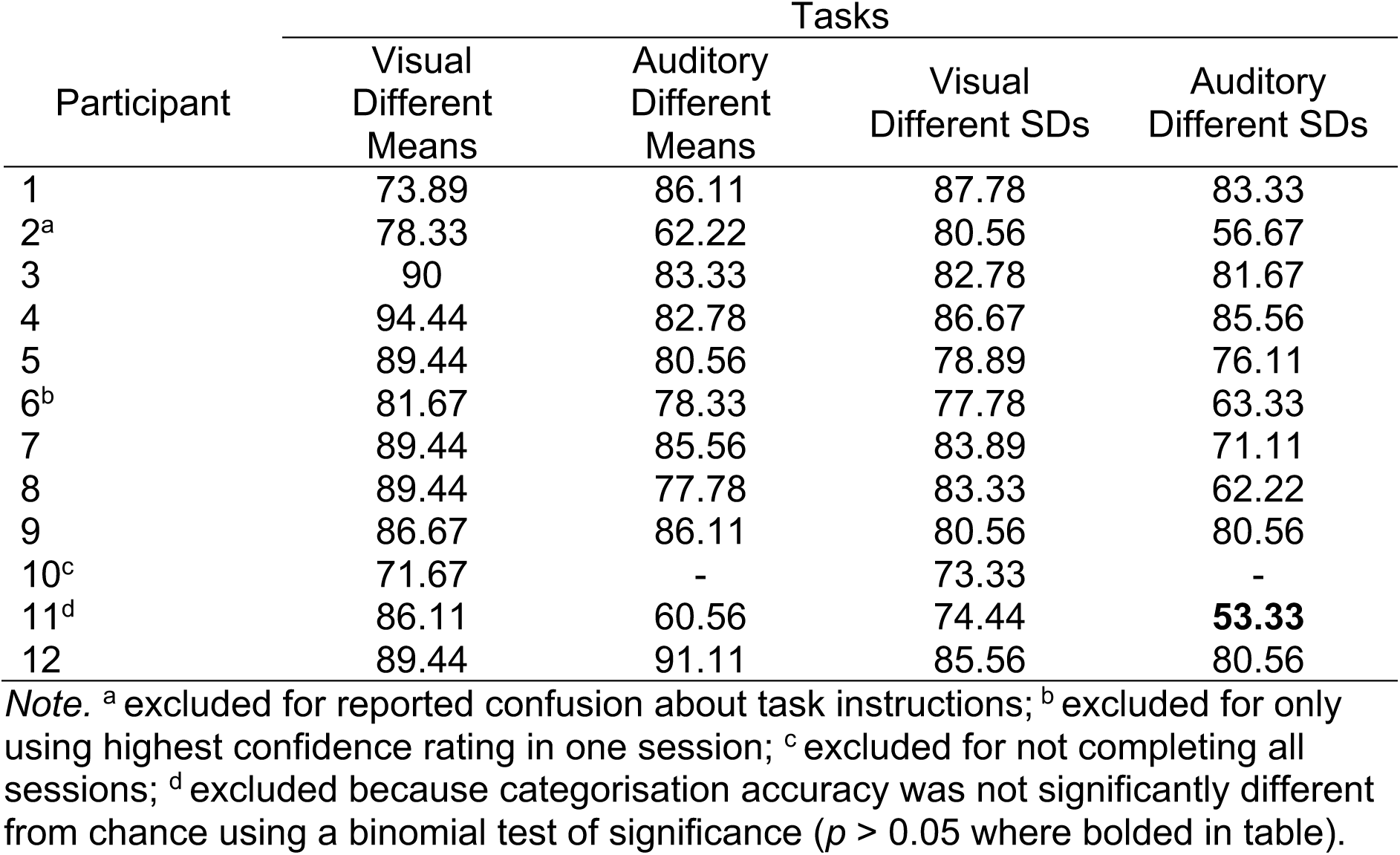
Categorisation Accuracy for Highest Intensity Stimuli and Exclusions

**Supplementary Table 2.** GLMM Follow-Up Tests: The Effect of the Evidence Predictor at all Intensity Levels

|  |  | Tasks |  |  |  |
| --- | --- | --- | --- | --- | --- |
|  |  | Visual<br>Different<br>Means | Auditory<br>Different<br>Means | Visual<br>Different<br>SDs | Auditory<br>Different<br>SDs |
|  |  | Category |  |  |  |
| Intensity 1 | $\beta$ | 0.27 | 0.00 | 0.17 | 0.00 |
| | $CI_{lower}$ | 0.13 | -0.16 | 0.04 | -0.17 |
| | $CI_{upper}$ | 0.42 | 0.16 | 0.29 | 0.16 |
| Intensity 2 | $\beta$ | 1.12 | 0.12 | 0.64 | 0.09 |
| | $CI_{lower}$ | 0.84 | -0.05 | 0.37 | -0.11 |
| | $CI_{upper}$ | 1.39 | 0.30 | 0.91 | 0.28 |
| Intensity 3 | $\beta$ | 1.99 | 2.36 | 1.45 | 1.40 |
| | $CI_{lower}$ | 1.56 | 1.78 | 1.19 | 0.90 |
| | $CI_{upper}$ | 2.42 | 2.95 | 1.71 | 1.89 |
| Intensity 4 | $\beta$ | 3.42 | 2.97 | 2.73 | 1.74 |
| | $CI_{lower}$ | 2.54 | 2.23 | 2.33 | 1.11 |
| | $CI_{upper}$ | 4.31 | 3.72 | 3.14 | 2.37 |
|  |  | Confidence |  |  |  |
| Intensity 1 | $\beta$ | 0.01 | 0.01 | 0.03 | 0.00 |
| | $CI_{lower}$ | -0.03 | -0.01 | -0.01 | -0.02 |
| | $CI_{upper}$ | 0.04 | 0.03 | 0.07 | 0.01 |
| Intensity 2 | $\beta$ | 0.11 | 0.02 | 0.05 | 0.01 |
| | $CI_{lower}$ | 0.06 | -0.03 | 0.01 | -0.01 |
| | $CI_{upper}$ | 0.16 | 0.07 | 0.09 | 0.03 |
| Intensity 3 | $\beta$ | 0.23 | 0.25 | 0.19 | 0.25 |
| | $CI_{lower}$ | 0.15 | 0.17 | 0.13 | 0.14 |
| | $CI_{upper}$ | 0.31 | 0.34 | 0.25 | 0.36 |
| Intensity 4 | $\beta$ | 0.43 | 0.26 | 0.30 | 0.25 |
| | $CI_{lower}$ | 0.29 | 0.16 | 0.23 | 0.13 |
| | $CI_{upper}$ | 0.56 | 0.36 | 0.37 | 0.37 |

### Results

#### Model Fitting

Models were initialised with random starting values sampled from a normal distribution which spanned a reasonable and sufficiently large range, given the parameters of each model. To mitigate against the possibility of finding parameter estimates that corresponded to a local minimum in the parameter space, we used between 10-20 random starting values for each model and confirmed convergence of different starting values to similar best-fitting parameter estimates. The same set of starting values were used for each subject in each modality, where possible.

#### Parameter Recovery for Different SDs Task Models

We explored the parameter recovery properties of the core models: distance, linear, quadratic, free-exponent, free-exponent with orientation dependent noise, log posterior probability ratio, log posterior probability ratio with free category distribution parameters, the log posterior probability ratio with decision noise and the log posterior probability ratio with orientation dependent noise. Data were simulated using a set of parameters for a given model. We then fit the simulated data with the same model and assessed whether we were able to recover the data-generating parameter values from the fitting procedure. We used stimuli that were similar to those used in the different SDs task, resampling stimulus values from the category distributions. To examine the effect of sample size on parameter recovery, we simulated data sets with sample sizes of: a) 360 trials (half the number of trials per participant in the main experiment), b) 720 trials (the same number of trials per participant), c) 1440 trials and d) 7200 trials. As seen in Supplementary Figures 1 – 9, as the sample size increased, the parameters were recovered better, as expected. However, even at smaller sample sizes, the parameters were recovered quite well for all models. Of note, some of the more extreme boundary parameters (boundary 6 and boundary 7) did not recover as well, especially at low sample sizes. This was particularly noticeable for the Bayesian models (Supplementary Figures 2 – 5). We suspected that this was because, by definition, the majority of stimulus values were sampled from around the means of the category distributions and thus, there were fewer simulated responses to constrain estimates of the boundaries further away from the means. We investigated this by sampling stimulus values that were equally spaced between a reasonable minimum and maximum value of size 720 trials. We simulated responses according to the fixed, free-exponent and Bayesian models and then determined whether the recovery of the generating parameters had improved. As shown in Supplementary Figure 10, the same parameters, particularly boundary 6 and boundary 7, were recovered better with the uniformly spaced data (compare to column 2 in Figure 2)

**Supplementary Figure 1.**
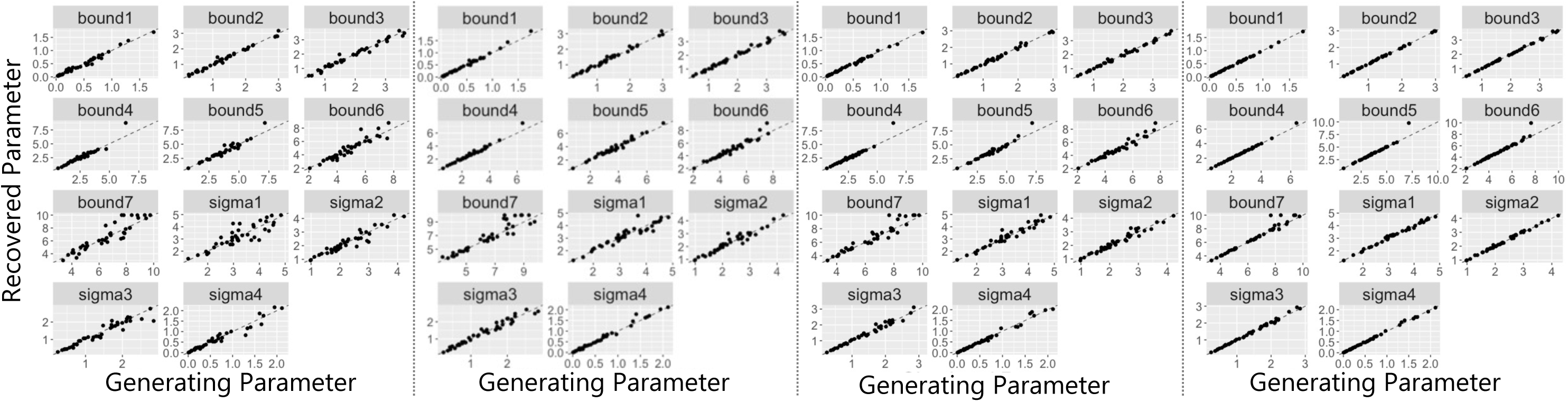
Parameter Recovery for Distance Model. We randomly generated stimulus values and simulated category and confidence responses using a set of parameters (generating parameters plotted on the x axis). We then fit the model to these simulated responses to determine whether we could recover the generating parameters (recovered parameters plotted on the y axis). The simulated datasets consisted of either 360 (first panel from left), 720 (second panel), 1440 (third panel) and 7200 trials (fourth panel). Bound 1 – 7 refer to the category/confidence boundaries and sigma 1 – 4 refer to the estimated noise parameters at each intensity level (the perceived level of sensory uncertainty in the observer’s perception of the stimulus). Dotted diagonal lines indicate perfect recovery of the data-generating parameter values.

**Supplementary Table 3.** Correlations Between Generating and Recovered Parameters for Unscaled Evidence Strength Models

| Parameter | Size of Simulated Dataset (Trials) |  |  |  |
| --- | --- | --- | --- | --- |
|  | 360 | 720 | 1440 | 7200 |
| Distance Model |  |  |  |  |
| Bound 1 | 0.99 | 1.00 | 1.00 | 1.00 |
| Bound 2 | 0.99 | 0.99 | 1.00 | 1.00 |
| Bound 3 | 0.99 | 0.99 | 1.00 | 1.00 |
| Bound 4 | 0.96 | 0.99 | 0.96 | 1.00 |
| Bound 5 | 0.96 | 0.99 | 0.98 | 0.97 |
| Bound 6 | 0.94 | 0.97 | 0.98 | 0.98 |
| Bound 7 | 0.92 | 0.93 | 0.95 | 0.98 |
| Sigma 1 | 0.87 | 0.95 | 0.97 | 1.00 |
| Sigma 2 | 0.95 | 0.96 | 0.98 | 1.00 |
| Sigma 3 | 0.96 | 0.98 | 0.99 | 1.00 |
| Sigma 4 | 0.98 | 0.99 | 0.99 | 1.00 |

**Supplementary Figure 2.**
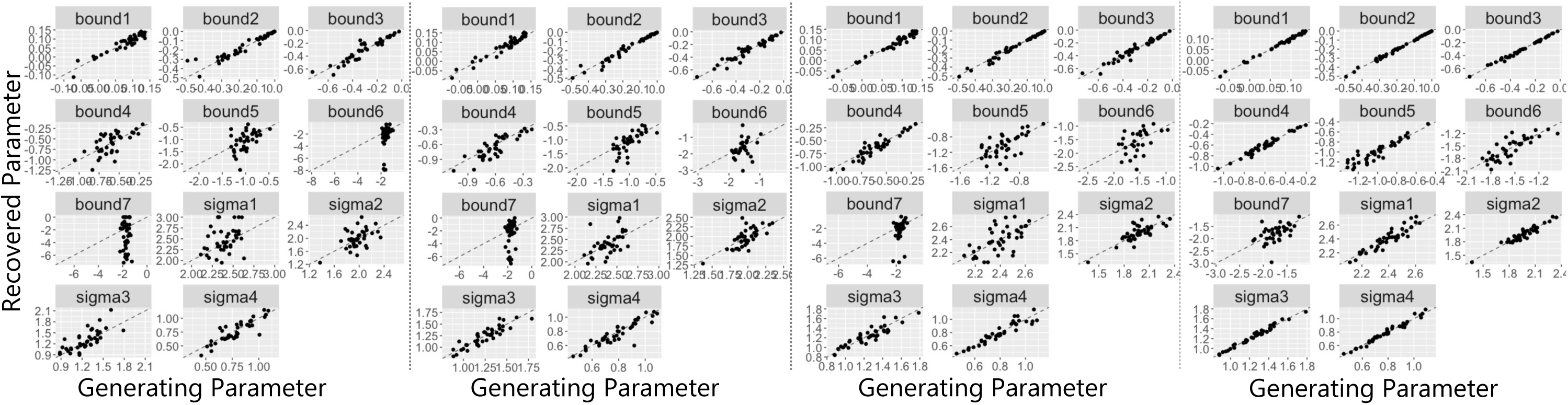
Parameter Recovery for Standard Bayesian Model. We randomly generated stimulus values and simulated category and confidence responses using a set of parameters (generating parameters plotted on the x axis) according to the standard Bayesian model (log posterior probability ratio model). We then fit the model to these simulated responses to determine whether we could recover the generating parameters (recovered parameters plotted on the y axis). The simulated datasets consisted of either 360 (first panel from left), 720 (second panel), 1440 (third panel) and 7200 trials (fourth panel). Bound 1 – 7 refer to the category/confidence boundaries in *d* units (log posterior probability ratio space) and sigma 1 – 4 refer to the estimated noise parameters at each intensity level (the perceived level of sensory uncertainty in the observer’s perception of the stimulus). Dotted diagonal lines indicate perfect recovery of the data-generating parameter values.

**Supplementary Figure 3.**
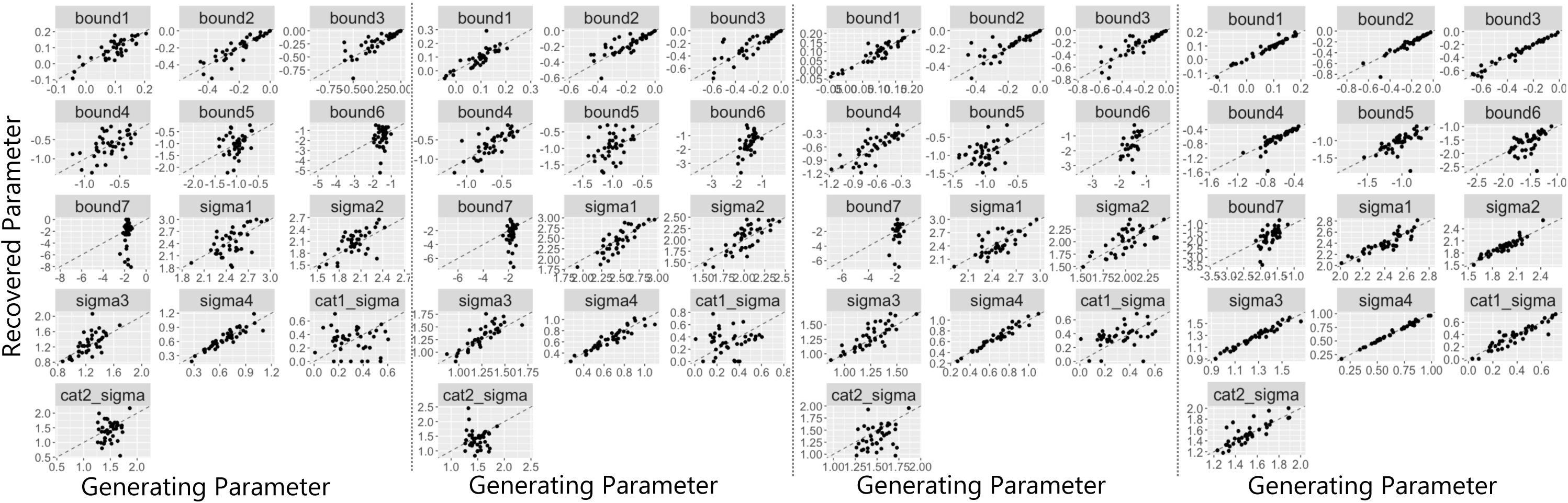
Parameter Recovery for Bayesian Model with Free Category Distribution Parameters. We randomly generated stimulus values and simulated category and confidence responses using a set of parameters (generating parameters plotted on the x axis) according to the Bayesian model with free category distribution parameters. We then fit the model to these simulated responses to determine whether we could recover the generating parameters (recovered parameters plotted on the y axis). The simulated datasets consisted of either 360 (first panel from left), 720 (second panel), 1440 (third panel) and 7200 trials (fourth panel). Bound 1 – 7 refer to the category/confidence boundaries in *d* units (log posterior probability ratio space), sigma 1 – 4 refer to the estimated noise parameters at each intensity level (the perceived level of sensory uncertainty in the observer’s perception of the stimulus), ‘cat1_sigma’ refers to the estimated standard deviation of the category 1 distribution and ‘cat2_sigma’ refers to the estimated standard deviation of the category 2 distribution. Dotted diagonal lines indicate perfect recovery of the data-generating parameter values.

**Supplementary Figure 4.**
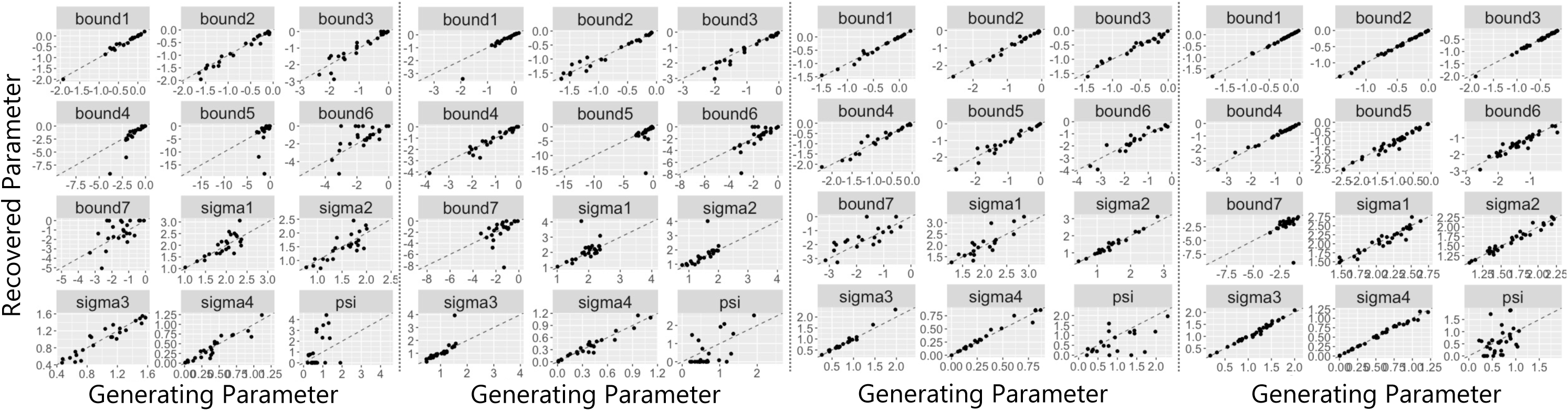
Parameter Recovery for Bayesian Model with Orientation Dependent Noise. We randomly generated stimulus values and simulated category and confidence responses using a set of parameters (generating parameters plotted on the x axis) according to the Bayesian model with orientation dependent noise. We then fit the model to these simulated responses to determine whether we could recover the generating parameters (recovered parameters plotted on the y axis). The simulated datasets consisted of either 360 (first panel from left), 720 (second panel), 1440 (third panel) and 7200 trials (fourth panel). Bound 1 – 7 refer to the category/confidence boundaries in *d* units (log posterior probability ratio space), sigma 1 – 4 refer to the estimated noise parameters at each intensity level (the perceived level of sensory uncertainty in the observer’s perception of the stimulus) and ‘psi’ refers to the orientation dependent noise parameter (see Equation 2). Dotted diagonal lines indicate perfect recovery of the data-generating parameter values.

**Supplementary Figure 5.**
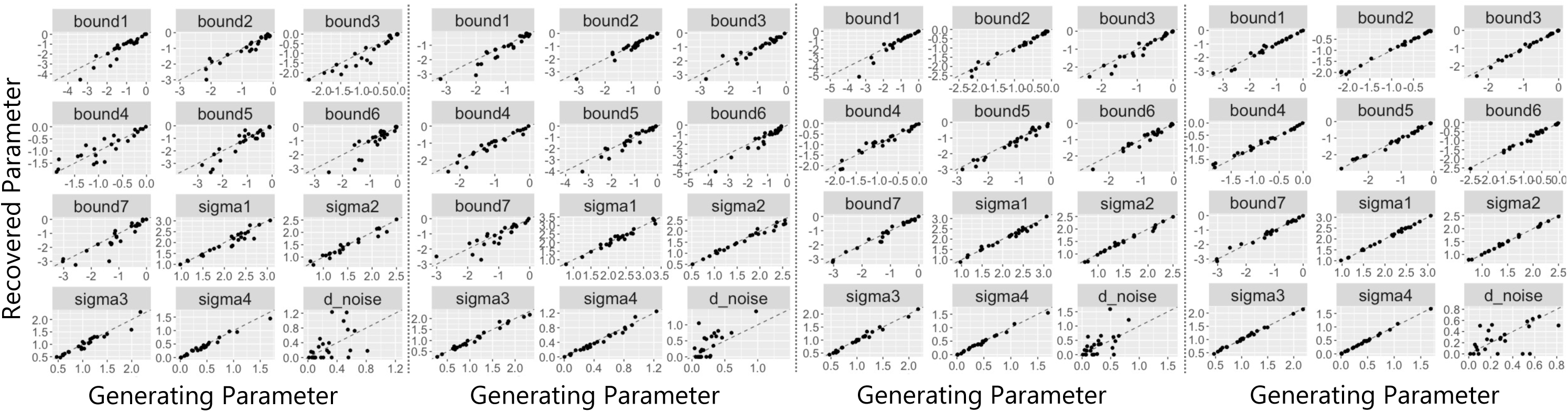
Parameter Recovery for Bayesian Model with Decision Noise. We randomly generated stimulus values and simulated category and confidence responses using a set of parameters (generating parameters plotted on the x axis) according to the Bayesian model with decision noise. We then fit the model to these simulated responses to determine whether we could recover the generating parameters (recovered parameters plotted on the y axis). The simulated datasets consisted of either 360 (first panel from left), 720 (second panel), 1440 (third panel) and 7200 trials (fourth panel). Bound 1 – 7 refer to the category/confidence boundaries in *d* units (log posterior probability ratio space), sigma 1 – 4 refer to the estimated noise parameters at each intensity level (the perceived level of sensory uncertainty in the observer’s perception of the stimulus), and ‘d_noise’ refers to the decision noise parameter (a Gaussian noise term on *d*). Dotted diagonal lines indicate perfect recovery of the data-generating parameter values.

**Supplementary Table 4.** Correlations Between Generating and Recovered Parameters for Bayesian Models

| Parameter | Size of Simulated Dataset (Trials) |  |  |  |
| --- | --- | --- | --- | --- |
|  | 360 | 720 | 1440 | 7200 |
| Standard Bayesian Model |  |  |  |  |
| Bound 1 | 0.95 | 0.96 | 0.98 | 1.00 |
| Bound 2 | 0.95 | 0.98 | 0.99 | 1.00 |
| Bound 3 | 0.94 | 0.96 | 0.97 | 1.00 |
| Bound 4 | 0.67 | 0.85 | 0.93 | 0.98 |
| Bound 5 | 0.50 | 0.71 | 0.62 | 0.92 |
| Bound 6 | 0.38 | 0.31 | 0.46 | 0.81 |
| Bound 7 | -0.02 | 0.22 | 0.29 | 0.51 |
| Sigma 1 | 0.34 | 0.54 | 0.62 | 0.93 |
| Sigma 2 | 0.50 | 0.72 | 0.78 | 0.95 |
| Sigma 3 | 0.73 | 0.90 | 0.91 | 0.99 |
| Sigma 4 | 0.87 | 0.91 | 0.94 | 0.99 |
| Bayesian Model with Free Category Distribution Parameters |  |  |  |  |
| Bound 1 | 0.86 | 0.85 | 0.92 | 0.98 |
| Bound 2 | 0.90 | 0.91 | 0.81 | 0.96 |
| Bound 3 | 0.87 | 0.91 | 0.84 | 0.97 |
| Bound 4 | 0.67 | 0.70 | 0.77 | 0.88 |
| Bound 5 | 0.17 | 0.36 | 0.34 | 0.71 |
| Bound 6 | 0.10 | 0.34 | 0.49 | 0.74 |
| Bound 7 | 0.04 | 0.03 | 0.34 | 0.59 |
| Cat. 1 Sigma | 0.24 | 0.50 | 0.39 | 0.17 |
| Cat. 2 Sigma | 0.70 | 0.67 | 0.34 | 0.58 |
| Sigma 1 | 0.65 | 0.46 | 0.69 | 0.93 |
| Sigma 2 | 0.60 | 0.64 | 0.68 | 0.88 |
| Sigma 3 | 0.51 | 0.76 | 0.82 | 0.90 |
| Sigma 4 | 0.90 | 0.95 | 0.97 | 0.98 |
| Bayesian Model with Orientation Dependent Noise |  |  |  |  |
| Bound 1 | 0.99 | 0.96 | 0.99 | 1.00 |
| Bound 2 | 0.98 | 0.98 | 0.99 | 1.00 |
| Bound 3 | 0.94 | 0.96 | 0.98 | 1.00 |
| Bound 4 | 0.89 | 0.96 | 0.97 | 0.99 |
| Bound 5 | 0.35 | 0.29 | 0.96 | 0.97 |
| Bound 6 | 0.61 | 0.73 | 0.94 | 0.94 |
| Bound 7 | 0.56 | 0.40 | 0.77 | 0.15 |
| Psi | 0.43 | 0.60 | 0.57 | 0.40 |
| Sigma 1 | 0.80 | 0.53 | 0.85 | 0.95 |
| Sigma 2 | 0.81 | 0.64 | 0.98 | 0.98 |
| Sigma 3 | 0.94 | 0.73 | 0.99 | 0.99 |
| Sigma 4 | 0.93 | 0.96 | 0.98 | 0.99 |

| Bayesian Model with Decision Noise |  |  |  |  |
| --- | --- | --- | --- | --- |
| Bound 1 | 0.93 | 0.95 | 0.96 | 0.99 |
| Bound 2 | 0.95 | 0.97 | 0.99 | 1.00 |
| Bound 3 | 0.95 | 0.96 | 0.97 | 1.00 |
| Bound 4 | 0.89 | 0.96 | 0.97 | 0.99 |
| Bound 5 | 0.92 | 0.96 | 0.98 | 0.99 |
| Bound 6 | 0.89 | 0.97 | 0.97 | 0.99 |
| Bound 7 | 0.92 | 0.87 | 0.98 | 0.99 |
| Dnoise | 0.45 | 0.72 | 0.60 | 0.54 |
| Sigma 1 | 0.96 | 0.98 | 0.99 | 1.00 |
| Sigma 2 | 0.97 | 0.99 | 0.99 | 1.00 |
| Sigma 3 | 0.97 | 0.99 | 0.99 | 1.00 |
| Sigma 4 | 0.99 | 0.99 | 0.99 | 1.00 |

**Supplementary Figure 6.**
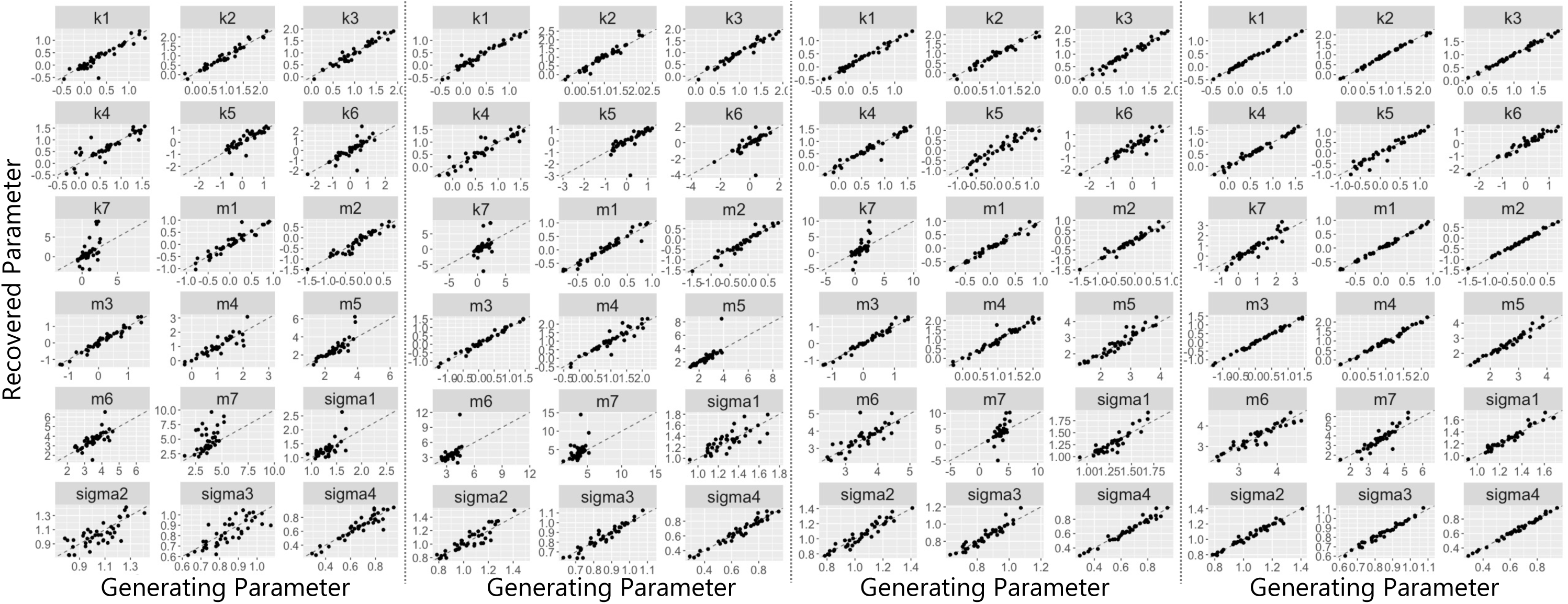
Parameter Recovery for Linear Model. We randomly generated stimulus values and simulated category and confidence responses using a set of parameters (generating parameters plotted on the x axis) according to the linear model. We then fit the model to these simulated responses to determine whether we could recover the generating parameters (recovered parameters plotted on the y axis). The simulated datasets consisted of either 360 (first panel from left), 720 (second panel), 1440 (third panel) and 7200 trials (fourth panel). Sigma 1 – 4 refer to the estimated noise parameters at each intensity level (the perceived level of sensory uncertainty in the observer’s perception of the stimulus). Parameters *k* 1 – 7 and *m* 1 – 7 are used to estimate the category/confidence boundaries according to *b* = *k* + *mσ* (see description for scaled evidence strength models for more details). Dotted diagonal lines indicate perfect recovery of the data-generating parameter values.

**Supplementary Figure 7.**
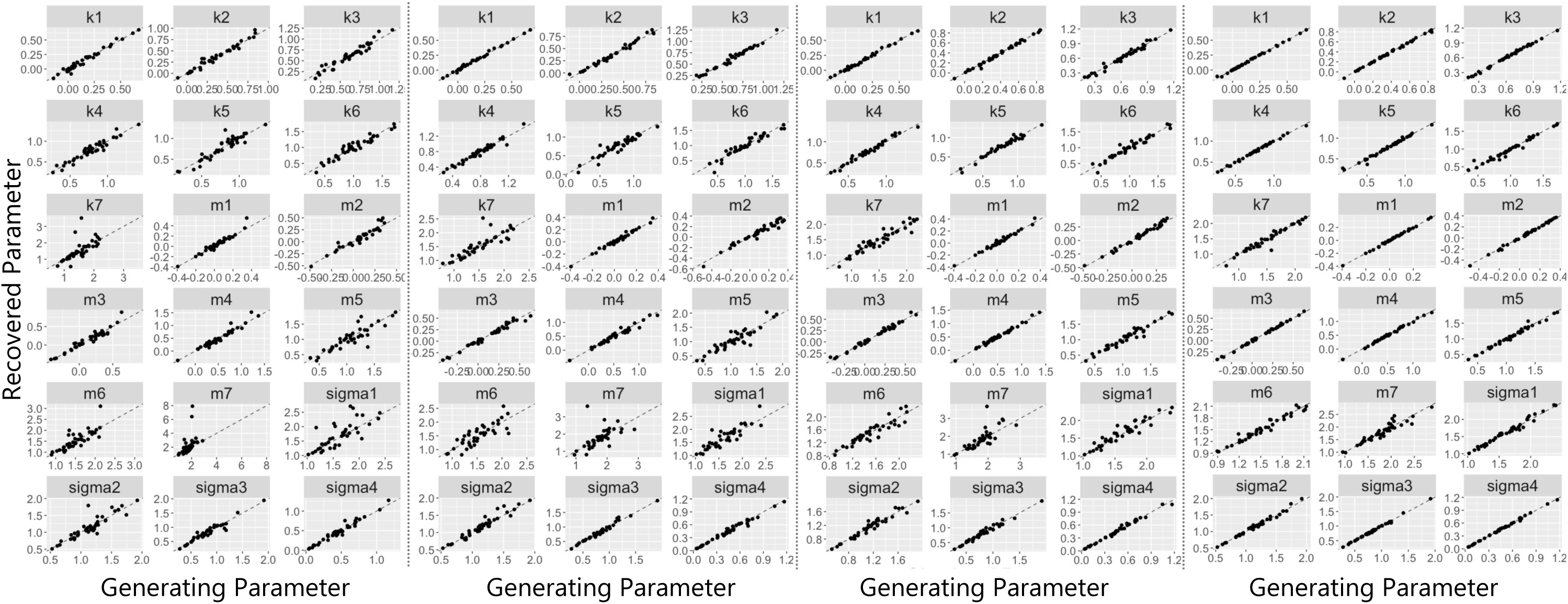
Parameter Recovery for Quadratic Model. We randomly generated stimulus values and simulated category and confidence responses using a set of parameters (generating parameters plotted on the x axis) according to the quadratic model. We then fit the model to these simulated responses to determine whether we could recover the generating parameters (recovered parameters plotted on the y axis). The simulated datasets consisted of either 360 (first panel from left), 720 (second panel), 1440 (third panel) and 7200 trials (fourth panel). Sigma 1 – 4 refer to the estimated noise parameters at each intensity level (the perceived level of sensory uncertainty in the observer’s perception of the stimulus). Parameters *k* 1 – 7 and *m* 1 – 7 are used to estimate the category/confidence boundaries according to *b* = *k* + *mσ^5^* (see description for scaled evidence strength models). Dotted diagonal lines indicate perfect recovery of the data-generating parameter values.

**Supplementary Figure 8.**
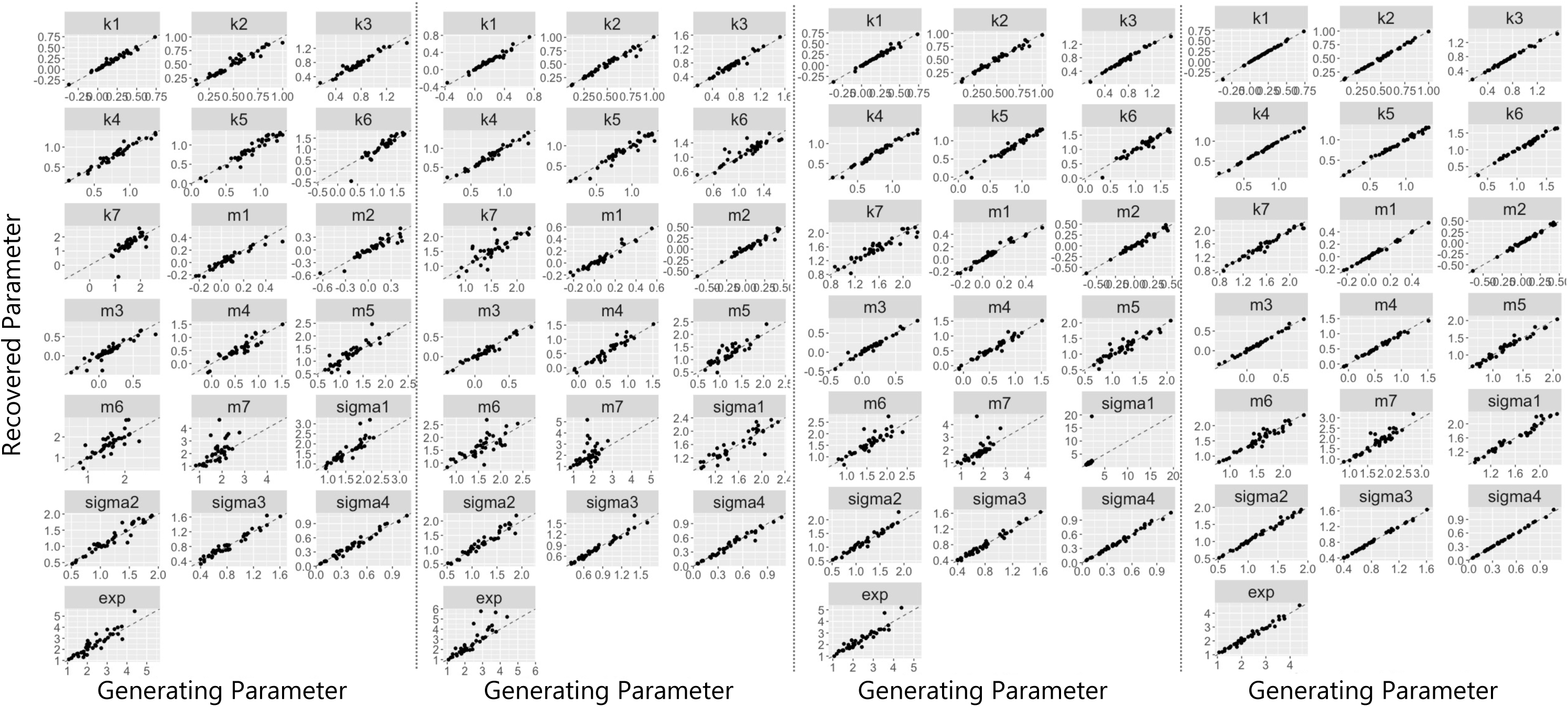
Parameter Recovery for Free-Exponent Model. We randomly generated stimulus values and simulated category and confidence responses using a set of parameters (generating parameters plotted on the x axis) according to the free-exponent model. We then fit the model to these simulated responses to determine whether we could recover the generating parameters (recovered parameters plotted on the y axis). The simulated datasets consisted of either 360 (first panel from left), 720 (second panel), 1440 (third panel) and 7200 trials (fourth panel). Sigma 1 – 4 refer to the estimated noise parameters at each intensity level (the perceived level of sensory uncertainty in the observer’s perception of the stimulus). Parameters ‘exp’ (the exponent), *k* 1 – 7 and *m* 1 – 7 are used to estimate the category/confidence boundaries according to *b* = *k* + *mσ^+-!^* (see description for scaled evidence strength models). Dotted diagonal lines indicate perfect recovery of the data-generating parameter values.

**Supplementary Figure 9.**
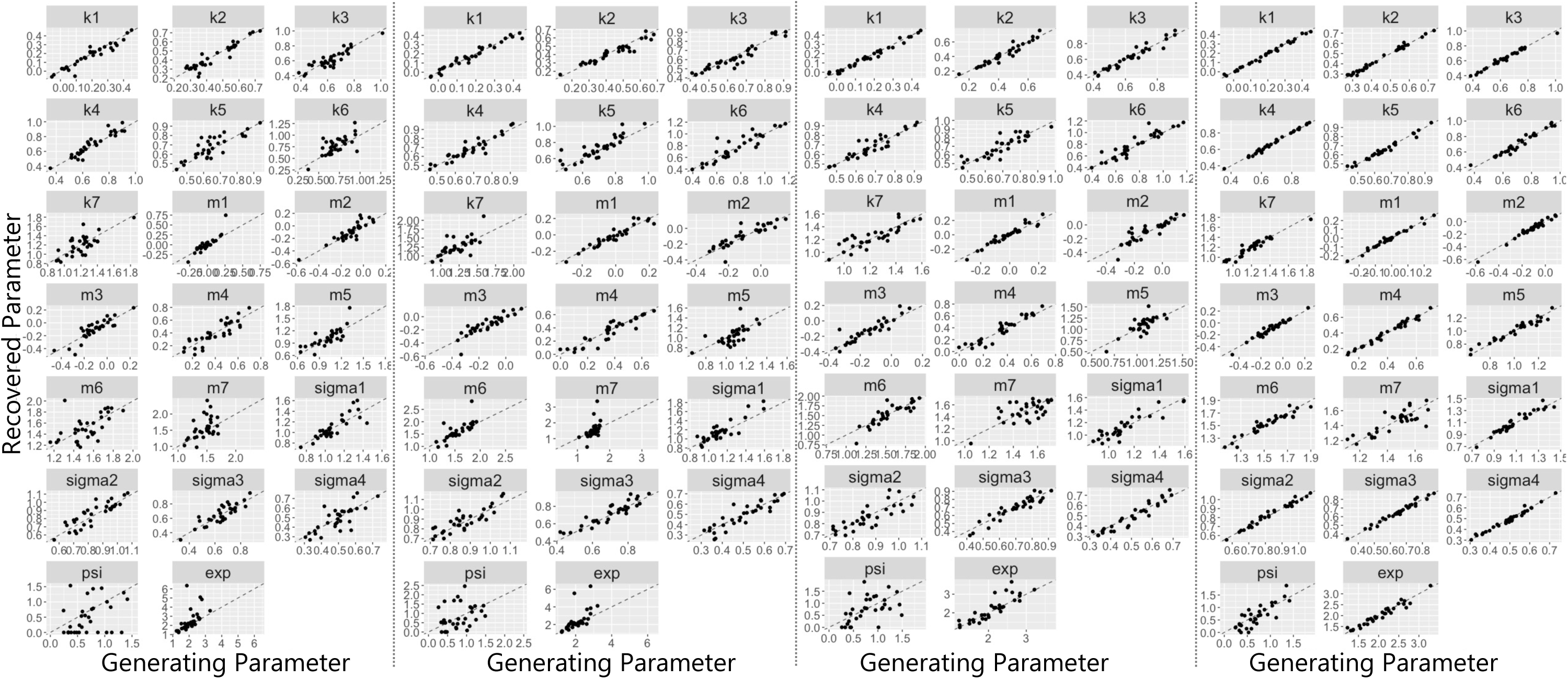
Parameter Recovery for Free-Exponent Model with Orientation Dependent Noise. We randomly generated stimulus values and simulated category and confidence responses using a set of parameters (generating parameters plotted on the x axis) according to the free-exponent model with orientation dependent noise. We then fit the model to these simulated responses to determine whether we could recover the generating parameters (recovered parameters plotted on the y axis). The simulated datasets consisted of either 360 (first panel from left), 720 (second panel), 1440 (third panel) and 7200 trials (fourth panel). Sigma 1 – 4 refer to the estimated noise parameters at each intensity level (the perceived level of sensory uncertainty in the observer’s perception of the stimulus) and ‘psi’ refers to the orientation dependent noise parameter (see Equation 2). Parameters ‘exp’ (the exponent), *k* 1 – 7 and *m* 1 – 7 are used to estimate the category/confidence boundaries according to xf*b* = *k* + *mσ^exp^* (see description for scaled evidence strength models). Dotted diagonal lines indicate perfect recovery of the data-generating parameter values.

**Supplementary Table 5.** Correlations Between Generating and Recovered Parameters for Scaled Evidence Strength Models

| Parameter | Size of Simulated Dataset |  |  |  |
| --- | --- | --- | --- | --- |
|  | 360 | 720 | 1440 | 7200 |
| Linear Model |  |  |  |  |
| k1 | 0.94 | 0.98 | 0.99 | 1.00 |
| k2 | 0.97 | 0.98 | 0.98 | 1.00 |
| k3 | 0.97 | 0.98 | 0.97 | 0.99 |
| k4 | 0.86 | 0.92 | 0.97 | 0.99 |
| k5 | 0.80 | 0.71 | 0.94 | 0.97 |
| k6 | 0.82 | 0.66 | 0.91 | 0.97 |
| k7 | 0.61 | 0.46 | 0.71 | 0.94 |
| m1 | 0.95 | 0.97 | 0.99 | 1.00 |
| m2 | 0.95 | 0.96 | 0.97 | 1.00 |
| m3 | 0.98 | 0.99 | 0.99 | 1.00 |
| m4 | 0.87 | 0.93 | 0.97 | 0.99 |
| m5 | 0.86 | 0.75 | 0.95 | 0.98 |
| m6 | 0.69 | 0.49 | 0.82 | 0.93 |
| m7 | 0.51 | 0.43 | 0.62 | 0.90 |
| Sigma 1 | 0.65 | 0.77 | 0.91 | 0.97 |
| Sigma 2 | 0.76 | 0.90 | 0.95 | 0.98 |
| Sigma 3 | 0.79 | 0.95 | 0.95 | 0.99 |
| Sigma 4 | 0.93 | 0.97 | 0.98 | 1.00 |

| Quadratic Model |  |  |  |  |
| --- | --- | --- | --- | --- |
| k1 | 0.99 | 1.00 | 1.00 | 1.00 |
| k2 | 0.98 | 0.99 | 1.00 | 1.00 |
| k3 | 0.96 | 0.99 | 0.99 | 1.00 |
| k4 | 0.96 | 0.98 | 0.99 | 1.00 |
| k5 | 0.94 | 0.94 | 0.98 | 0.99 |
| k6 | 0.96 | 0.96 | 0.97 | 0.98 |
| k7 | 0.69 | 0.86 | 0.95 | 0.97 |
| m1 | 0.97 | 0.99 | 0.99 | 1.00 |
| m2 | 0.95 | 0.98 | 0.99 | 1.00 |
| m3 | 0.96 | 0.99 | 0.99 | 1.00 |
| m4 | 0.97 | 0.98 | 0.99 | 1.00 |
| m5 | 0.89 | 0.91 | 0.97 | 0.99 |
| m6 | 0.80 | 0.82 | 0.92 | 0.98 |
| m7 | 0.45 | 0.57 | 0.78 | 0.94 |
| Sigma 1 | 0.77 | 0.88 | 0.94 | 0.99 |
| Sigma 2 | 0.90 | 0.96 | 0.97 | 0.99 |
| Sigma 3 | 0.96 | 0.99 | 0.98 | 1.00 |
| Sigma 4 | 0.98 | 0.99 | 0.99 | 1.00 |
| Free-Exponent Model |  |  |  |  |
| Exponent | 0.92 | 0.87 | 0.93 | 0.99 |
| k1 | 0.98 | 0.98 | 0.99 | 1.00 |
| k2 | 0.97 | 0.98 | 0.99 | 1.00 |
| k3 | 0.97 | 0.98 | 0.99 | 1.00 |
| k4 | 0.97 | 0.98 | 0.95 | 1.00 |
| k5 | 0.96 | 0.96 | 0.91 | 1.00 |
| k6 | 0.93 | 0.90 | 0.95 | 0.99 |
| k7 | 0.71 | 0.84 | 0.89 | 0.98 |
| m1 | 0.95 | 0.97 | 0.99 | 1.00 |
| m2 | 0.94 | 0.98 | 0.98 | 1.00 |
| m3 | 0.92 | 0.98 | 0.97 | 0.99 |
| m4 | 0.91 | 0.95 | 0.97 | 0.99 |
| m5 | 0.88 | 0.88 | 0.96 | 0.98 |
| m6 | 0.78 | 0.76 | 0.91 | 0.96 |
| m7 | 0.63 | 0.44 | 0.78 | 0.93 |
| Sigma 1 | 0.88 | 0.89 | 0.94 | 0.98 |
| Sigma 2 | 0.96 | 0.96 | 0.95 | 0.99 |
| Sigma 3 | 0.97 | 0.98 | 0.99 | 1.00 |
| Sigma 4 | 0.98 | 0.98 | 0.99 | 1.00 |

| Free-Exponent Model with Orientation Dependent Noise |  |  |  |  |
| --- | --- | --- | --- | --- |
| Exponent | 0.57 | 0.64 | 0.87 | 0.97 |
| k1 | 0.95 | 0.98 | 0.99 | 1.00 |
| k2 | 0.94 | 0.95 | 0.96 | 0.99 |
| k3 | 0.90 | 0.95 | 0.97 | 0.99 |
| k4 | 0.94 | 0.93 | 0.93 | 1.00 |
| k5 | 0.84 | 0.86 | 0.88 | 0.99 |
| k6 | 0.69 | 0.90 | 0.93 | 0.98 |
| k7 | 0.73 | 0.69 | 0.84 | 0.96 |
| m1 | 0.88 | 0.95 | 0.96 | 0.99 |
| m2 | 0.89 | 0.89 | 0.89 | 0.98 |
| m3 | 0.88 | 0.88 | 0.91 | 0.98 |
| m4 | 0.86 | 0.92 | 0.95 | 0.99 |
| m5 | 0.76 | 0.66 | 0.77 | 0.97 |
| m6 | 0.58 | 0.74 | 0.87 | 0.95 |
| m7 | 0.48 | 0.50 | 0.62 | 0.81 |
| Psi | 0.34 | 0.32 | 0.47 | 0.83 |
| Sigma 1 | 0.85 | 0.88 | 0.90 | 0.96 |
| Sigma 2 | 0.87 | 0.85 | 0.82 | 0.99 |
| Sigma 3 | 0.86 | 0.92 | 0.93 | 0.98 |
| Sigma 4 | 0.77 | 0.90 | 0.95 | 0.99 |

**Supplementary Figure 10.**
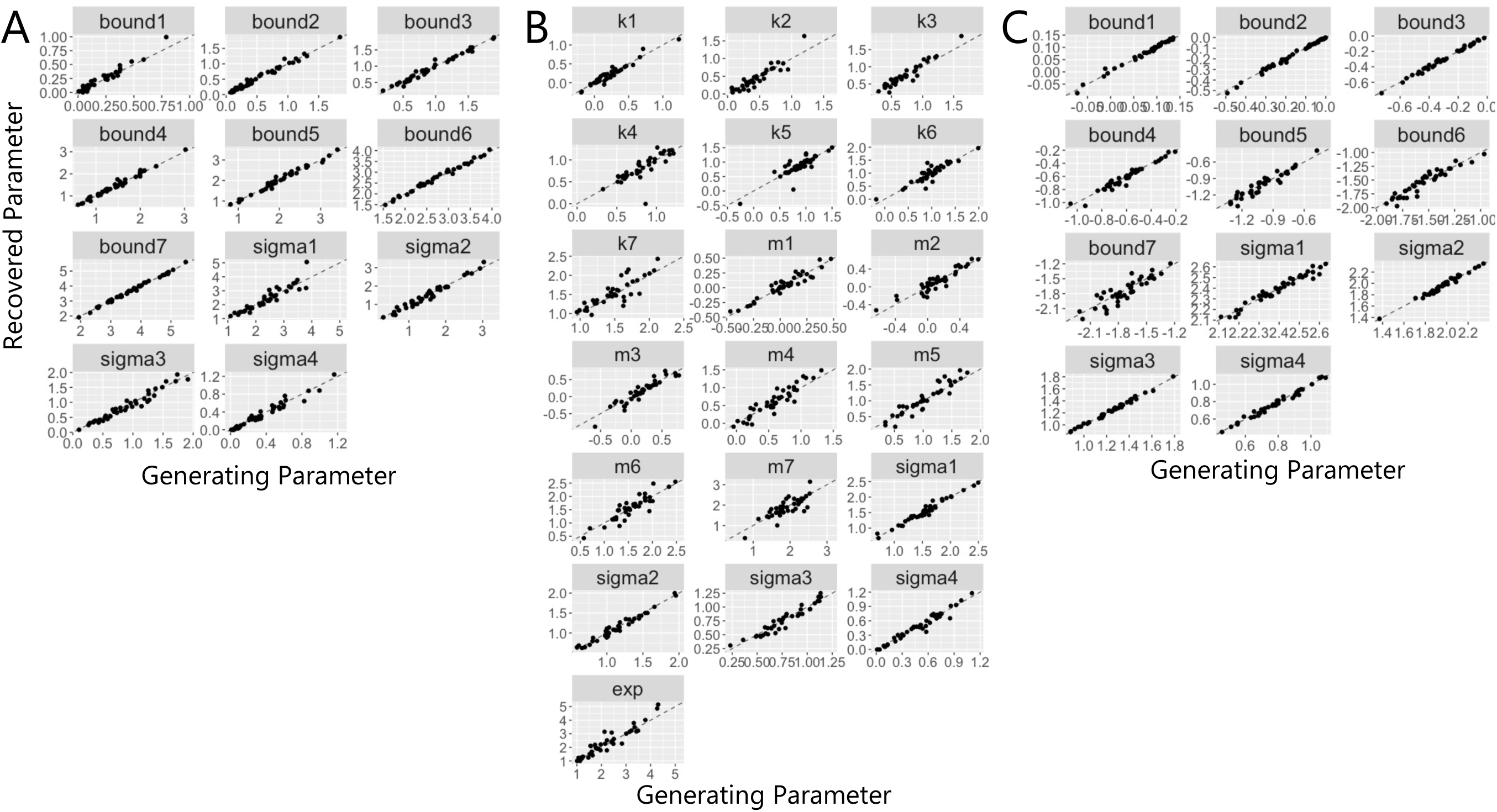
Parameter Recovery with Evenly Spaced Stimulus Values. To determine how the parameter recoveries were influenced by the sampling of stimulus values, rather than sample values from the category distributions we used values that were evenly spaced between a reasonable minimum and maximum. We simulated category and confidence responses according to (A) the fixed, (B) the free-exponent and (C) the standard Bayesian model. We used 720 trials per simulated dataset and found that the same parameters, particularly boundary 6 and boundary 7, were recovered better with the evenly spaced data (compare to column 2 in Figure 2).

**Supplementary Table 6.** Correlations Between Generating and Recovered Parameters for Models with Evenly Spaced Stimulus Values

| Distance Model |  | Free Exponent Model |  | Standard Bayesian Model |  |
| --- | --- | --- | --- | --- | --- |
| Parameter | Correlation | Parameter | Correlation | Parameter | Correlation |
| Bound 1 | 0.97 | Exponent | 0.95 | Bound 1 | 0.99 |
| Bound 2 | 0.99 | k1 | 0.96 | Bound 2 | 1.00 |
| Bound 3 | 0.99 | k2 | 0.92 | Bound 3 | 1.00 |
| Bound 4 | 0.99 | k3 | 0.96 | Bound 4 | 0.98 |
| Bound 5 | 0.99 | k4 | 0.80 | Bound 5 | 0.96 |
| Bound 6 | 0.99 | k5 | 0.86 | Bound 6 | 0.95 |
| Bound 7 | 1.00 | k6 | 0.91 | Bound 7 | 0.89 |
| Sigma 1 | 0.93 | k7 | 0.86 | Sigma 1 | 0.98 |
| Sigma 2 | 0.98 | m1 | 0.95 | Sigma 2 | 0.99 |
| Sigma 3 | 0.98 | m2 | 0.93 | Sigma 3 | 1.00 |
| Sigma 4 | 0.98 | m3 | 0.94 | Sigma 4 | 0.99 |
|  |  | m4 | 0.93 |  |  |
|  |  | m5 | 0.94 |  |  |
|  |  | m6 | 0.91 |  |  |
|  |  | m7 | 0.80 |  |  |
|  |  | Sigma 1 | 0.98 |  |  |
|  |  | Sigma 2 | 0.98 |  |  |
|  |  | Sigma 3 | 0.96 |  |  |
|  |  | Sigma 4 | 0.98 |  |  |

#### Control Study

Figure 4 shows that for the visual tasks, categorisation accuracy increased approximately linearly with increasing stimulus intensity, as expected. However, in the auditory tasks, the same pattern was not observed. That is, categorisation accuracy was closer to chance at the bottom two intensity levels and then around 75% for the top two intensity levels. To ensure that the difference in first order task performance did not account for the observed modelling results, we repeated the auditory different SDs task in 4 participants. We used different intensity levels (equivalent of -3 dB SPL, 4 dB SPL, 11dB SPL, 17 dB SPL for a 1 kHz pure tone) for the auditory task which were chosen based on the results of the main study and further piloting. Furthermore, we did not correct frequencies for equal loudness prior to adjusting stimulus intensity. This allowed for better comparison with the visual task in which orientation-based adjustments were not made to stimulus contrast. As intended, in the control study categorisation accuracy in the auditory task was approximately linearly increasing with stimulus intensity, as in the visual task (see Supplementary Figure 5).

**Supplementary Figure 11.**
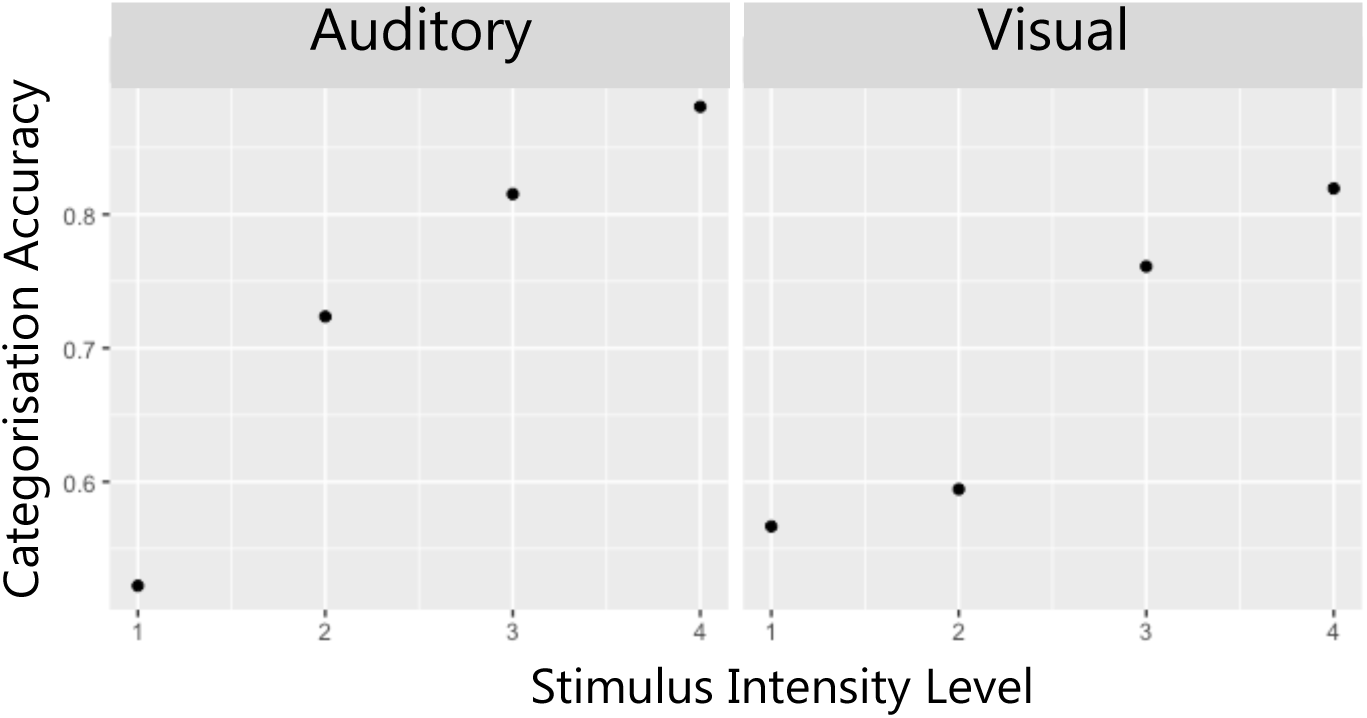
Control Experiment: Categorisation Accuracy as a Function of Stimulus Intensity. Categorisation accuracy in the control experiment increased approximately linearly with increasing stimulus intensity in the auditory task as in the visual task. Means are plotted for the same 4 participants who completed the main experiment (data used for visual task means) and the control experiment (data used for auditory task means).

#### Model Specification

All models were the same as in the main study, however, as the orientation-dependent noise parameter, *ψ*, did not reliably improve model fits (see Table 2) and did not affect the conclusions, we did not include it in any variations of the models in the control study.

### Results

In line with the results from the main experiment, the model-based analysis found that the best performing model for all subjects is one from the scaled evidence strength class (25% for linear, 50% for quadratic and 25% for free-exponent). This preference for the scaled evidence strength models is consistent with the summed *AIC* and *BIC* values which were the lowest for the free-exponent model (see Supplementary Table 7). As in the main experiment, we also wanted to determine if different parameter settings were required to account for the data across modalities. We used the same participants’ data from the visual task from the main experiment and found that the flexible settings model is the best performing model (see Supplementary Table 8) on average, according to the summed *AIC* and *BIC* scores.

At the individual level, for half of the subjects the best performing model is the flexible settings model and for the other half of the subjects the best performing model is the different noise settings model (see ‘Parameter Settings of the Free-Exponent Models Across Modalities’ in Model Specification for model descriptions). This means that for half of the subjects, the best performing model was one in which measurement noise differs across modalities, but the base positioning of the boundary parameters (without the measurement noise adjustment) stays the same. This finding suggests that when first order task performance is more similar across modalities, the metacognitive process may be influenced by the task-related variables (the boundaries) in the same way across modalities but is tuned to sensory processing differently (measurement noise).

**Supplementary Table 7.**
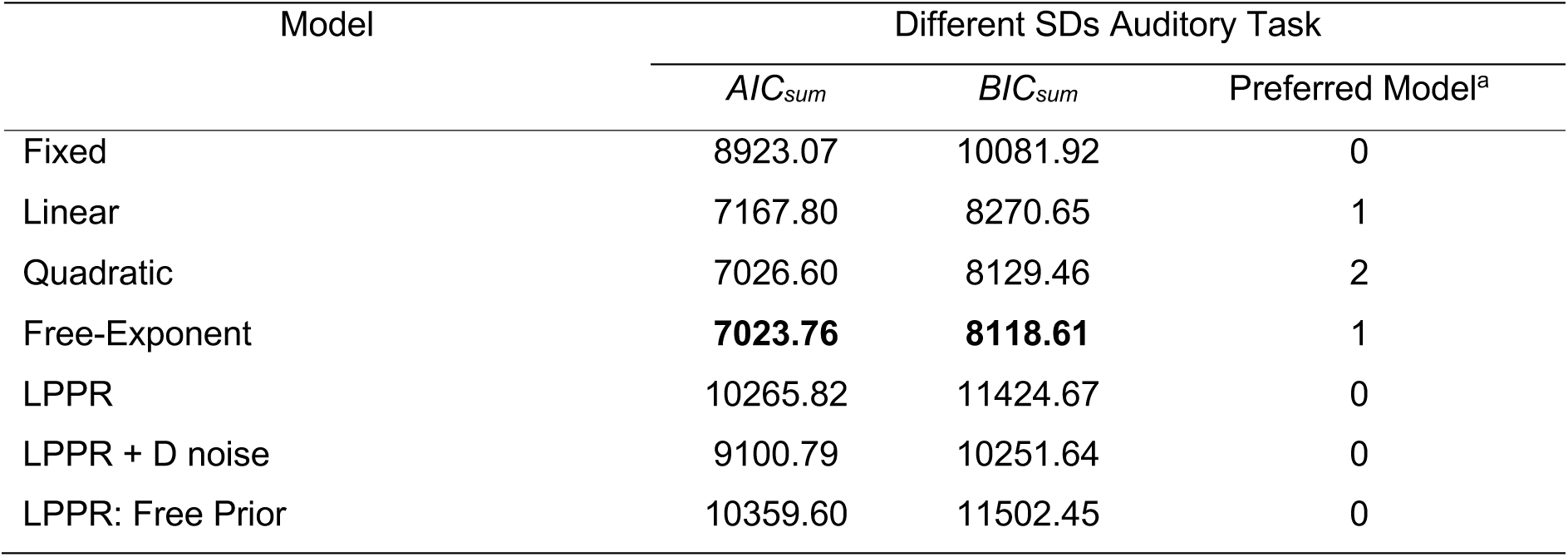
Model Fits for Control Data

**Supplementary Table 8.**
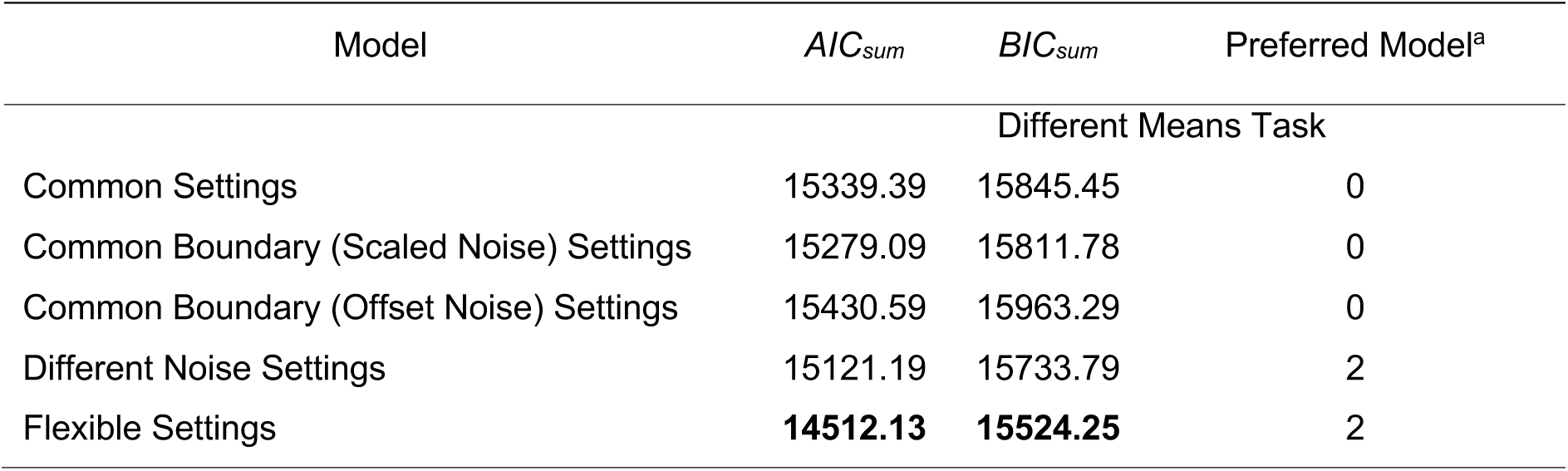
Parameter Settings Across Modalities for Control Data

**Supplementary Figure 12.**
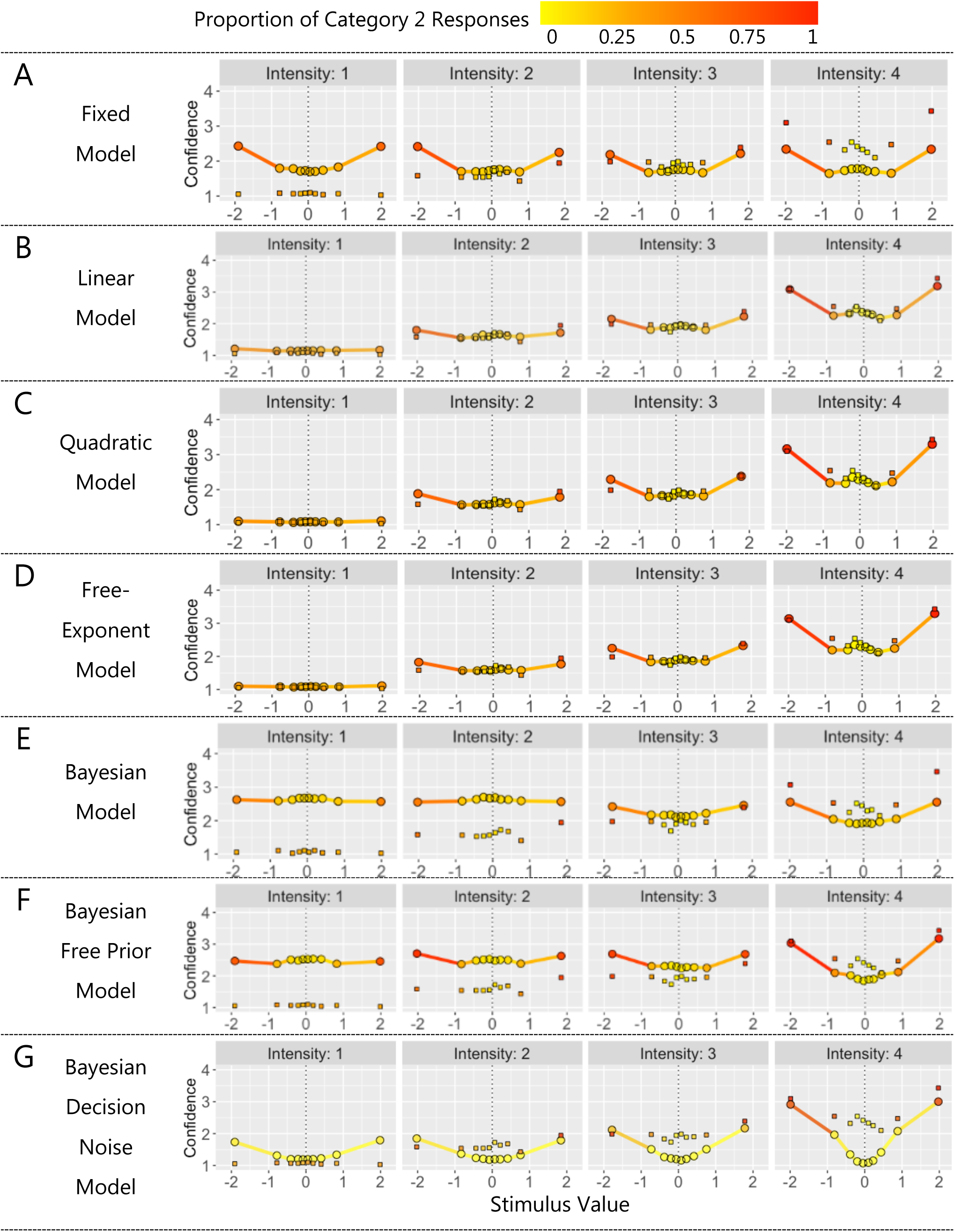
Model Comparison for Control Study. In all plots, mean confidence (y axis) and proportion of Category 2 responses (colour) for binned standardised stimulus values (x axis). Square data points show means for experimental data and solid lines and circular data points show means for model predictions.

### Alternative Visualisation of Model Fits from Main Experiment

Supplementary Figures 13-22 show the data and model predictions from the main experiment. These visualisations are an alternative to the plots shown in Figure 6, where we calculate mean ‘response’ per bin which averages over category and confidence. In the figures below, we do not collapse over category and confidence and instead show confidence on the x axes and use colour to show proportion of Category 2 responses. See figure labels for more details.

**Supplementary Figure 13.**
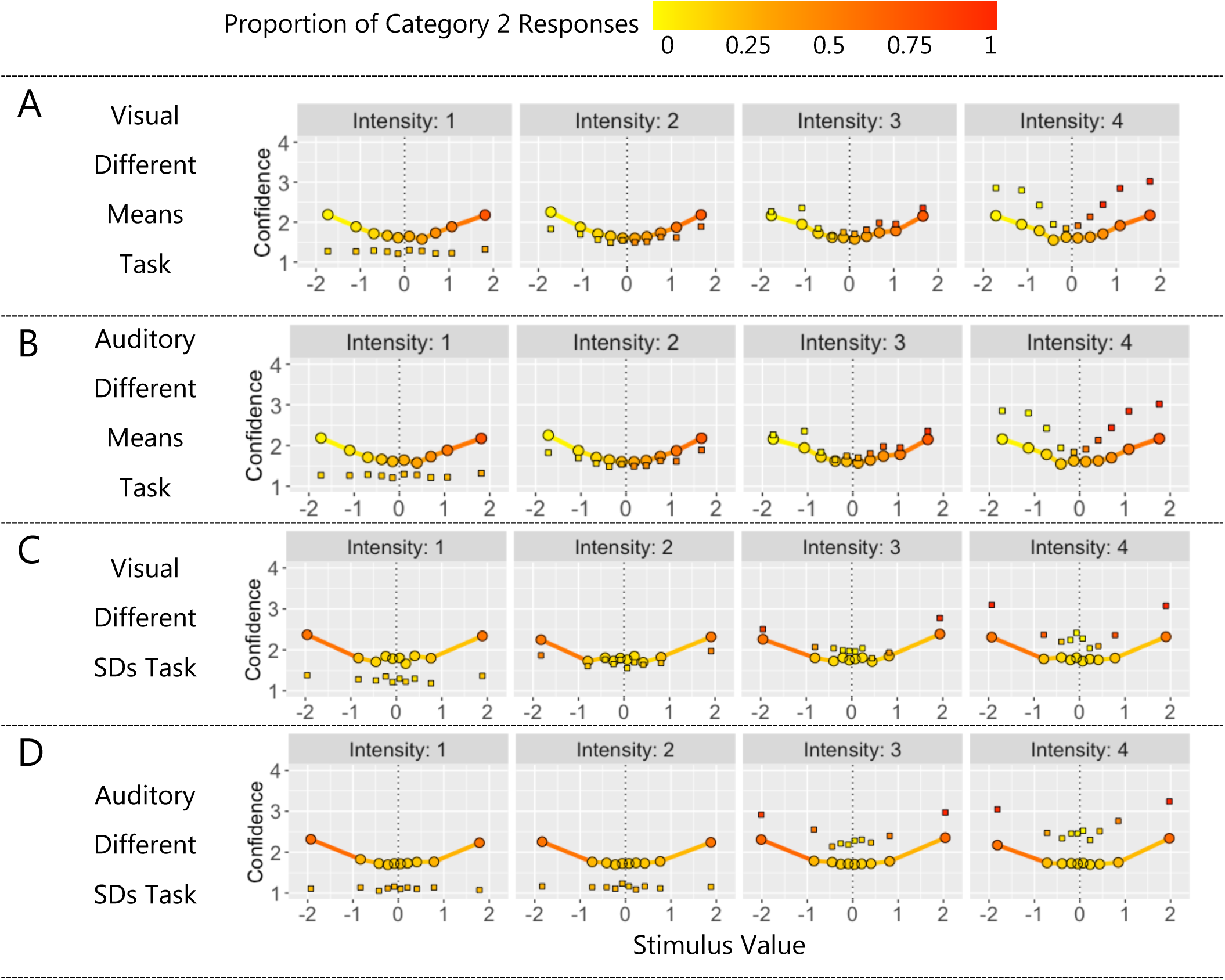
Unscaled-Evidence Strength Model: The Distance Model. In all plots, mean confidence (y axis) and proportion of Category 2 responses (colour) for binned standardised stimulus values (x axis). Square data points show means for experimental data and solid lines and circular data points show means for model predictions. Model was fit to data from each task and modality separately. Model predictions displayed for: (A) visual different means task, (B) auditory different means task, (C) visual different SDs task and (D) auditory different SDs task.

**Supplementary Figure 14.**
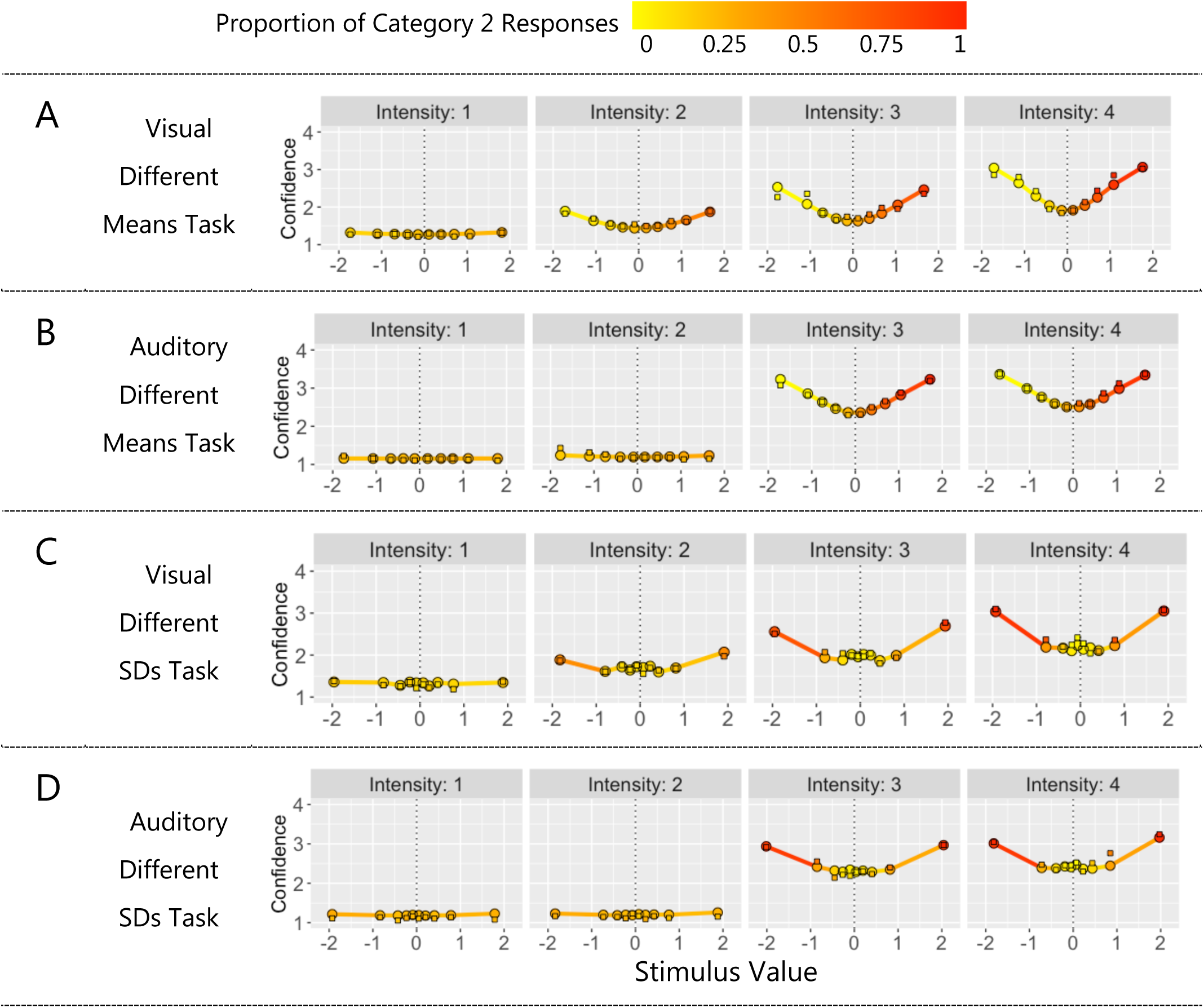
Scaled-Evidence Strength Model: The Linear Model. In all plots, mean confidence (y axis) and proportion of Category 2 responses (colour) for binned standardised stimulus values (x axis). Square data points show means for experimental data and solid lines and circular data points show means for model predictions. Model was fit to data from each task and modality separately. Model predictions displayed for: (A) visual different means task, (B) auditory different means task, (C) visual different SDs task and (D) auditory different SDs task.

**Supplementary Figure 15.**
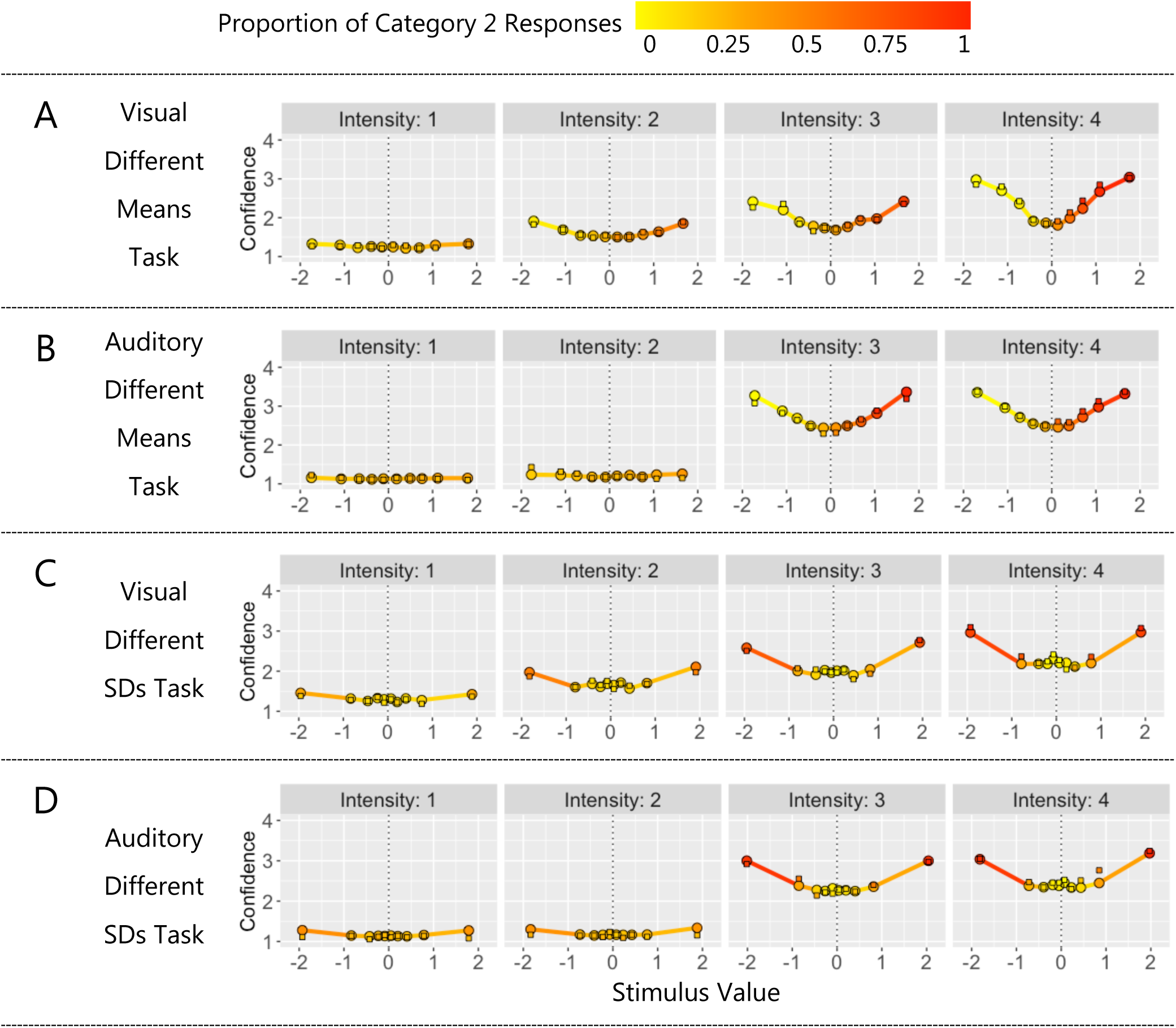
Scaled-Evidence Strength Model: The Quadratic Model. In all plots, mean confidence (y axis) and proportion of Cat 2 responses (colour) for binned standardised stimulus values (x axis). Square data points show means for experimental data and solid lines and circular data points show means for model predictions. Model was fit to data from each task and modality separately. Model predictions displayed for: (A) visual different means task, (B) auditory different means task, (C) visual different SDs task and (D) auditory different SDs task.

**Supplementary Figure 16.**
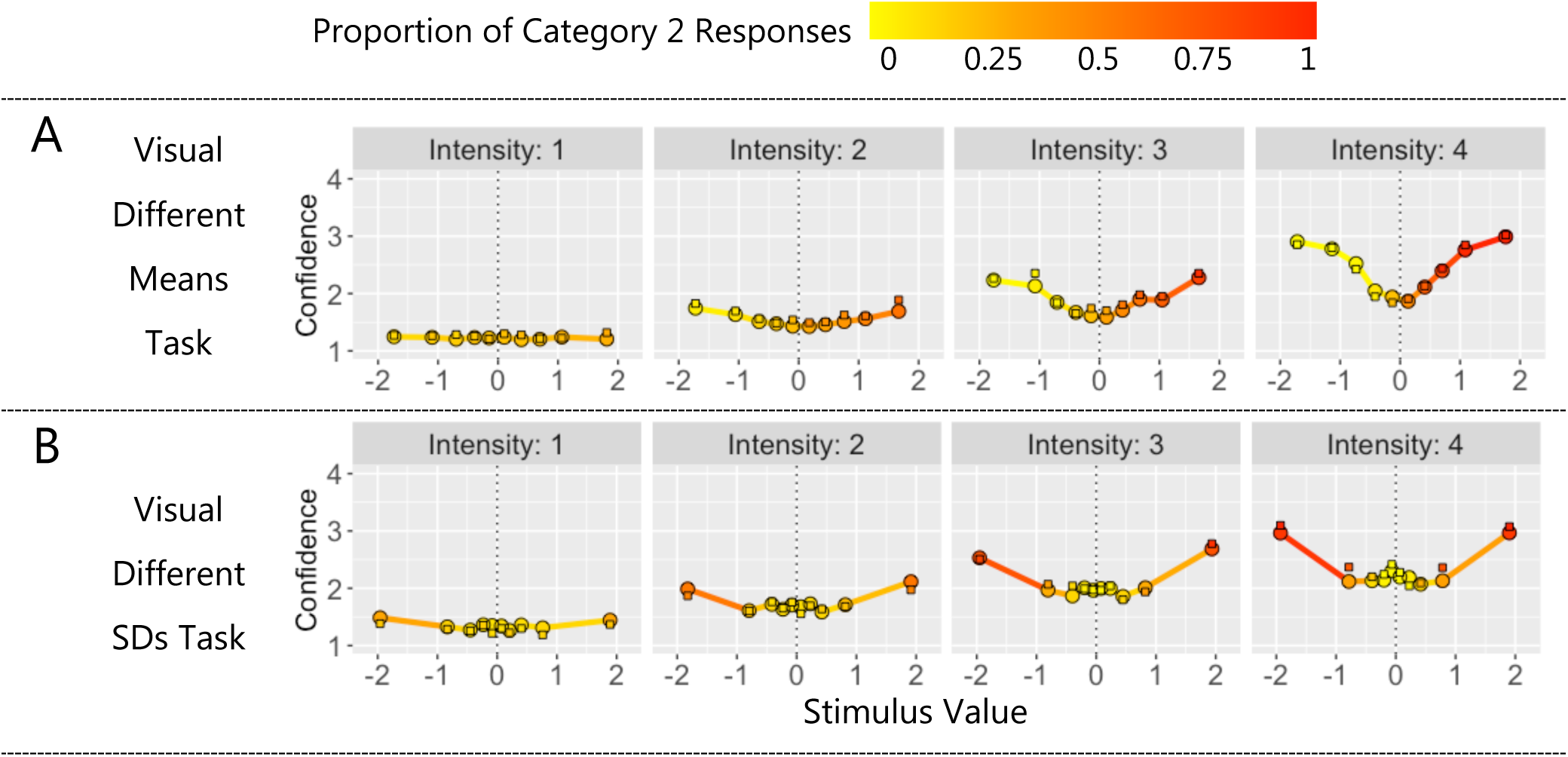
Scaled-Evidence Strength Model: The Free-Exponent Model with Orientation Dependent Noise. In all plots, mean confidence (y axis) and proportion of Category 2 responses (colour) for binned standardised stimulus values (x axis). Square data points show means for experimental data and solid lines and circular data points show means for model predictions. Model was fit to data from each task. Model predictions displayed for: (A) visual different means task and (B) visual different SDs task.

**Supplementary Figure 17.**
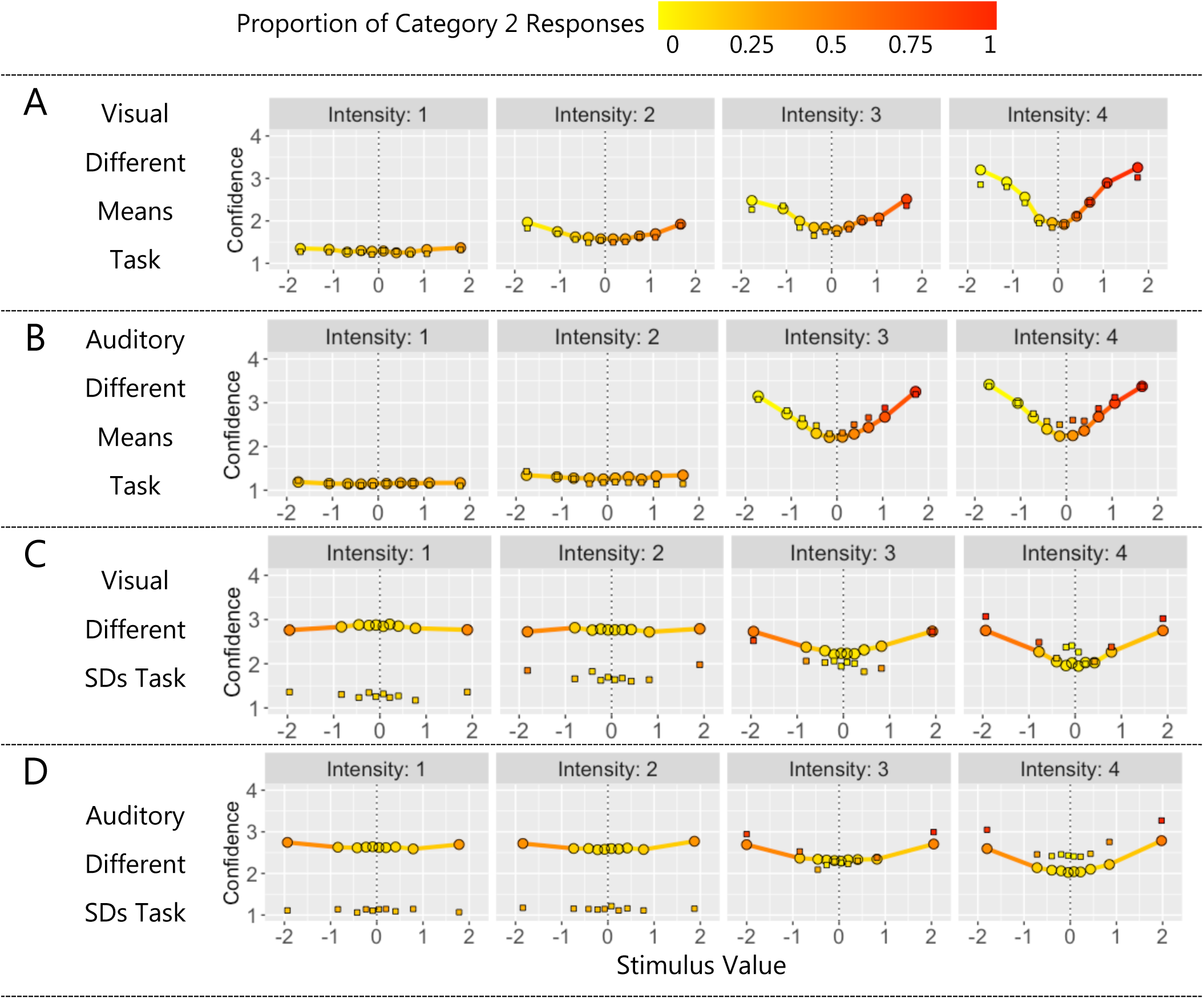
Bayesian Model: Log Posterior Probability Ratio. In all plots, mean confidence (y axis) and proportion of Category 2 responses (colour) for binned standardised stimulus values (x axis). Square data points show means for experimental data and solid lines and circular data points show means for model predictions. Model was fit to data from each task and modality separately. Model predictions displayed for: (A) visual different means task, (B) auditory different means task, (C) visual different SDs task and (D) auditory different SDs task.

**Supplementary Figure 18.**
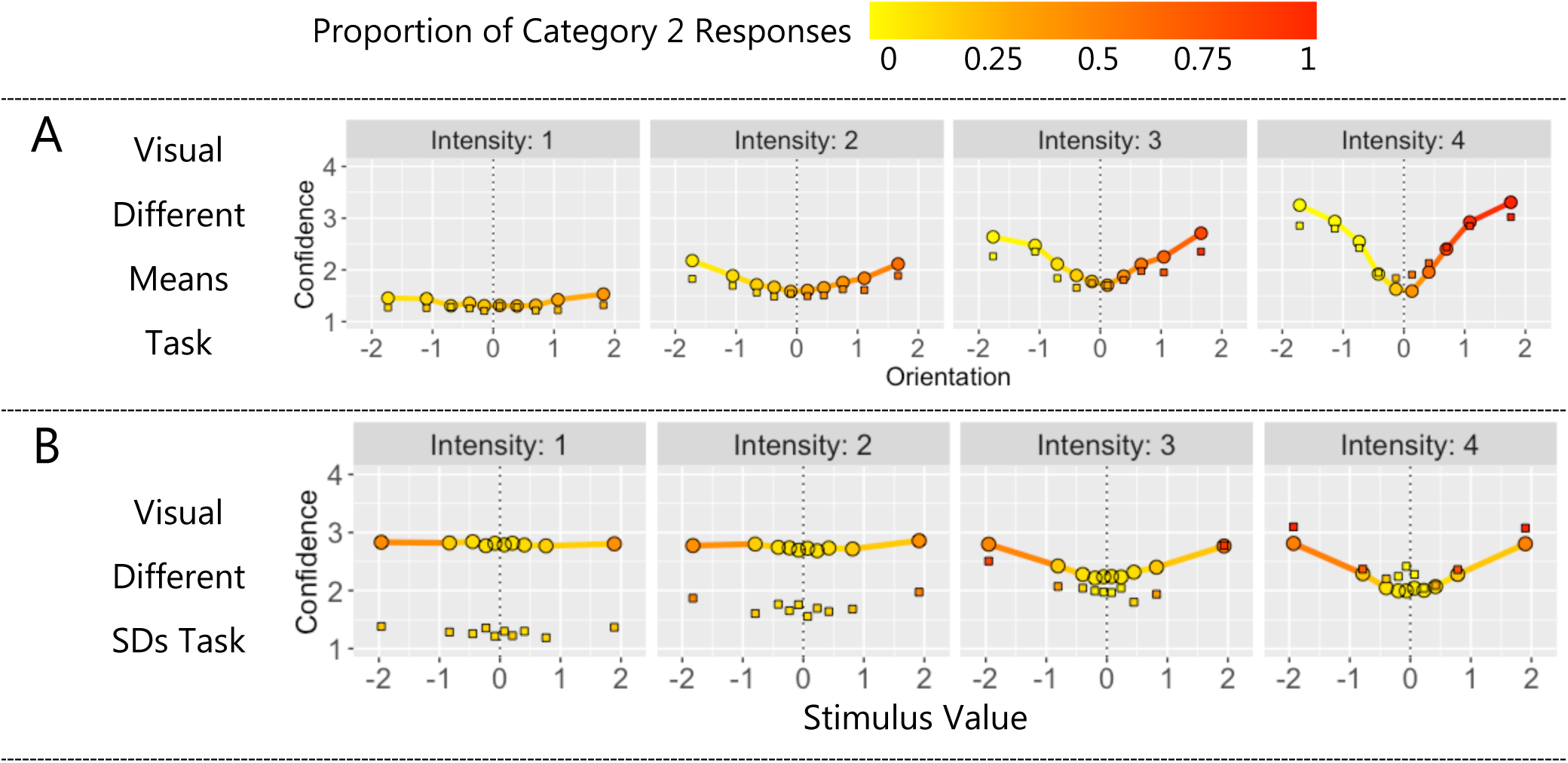
Bayesian Model: Log Posterior Probability Ratio with Orientation Dependent Noise. In all plots, mean confidence (y axis) and proportion of Category 2 responses (colour) for binned standardised stimulus values (x axis). Square data points show means for experimental data and solid lines and circular data points show means for model predictions. Model was fit to data from each task. Model predictions displayed for: (A) visual different means task and (B) visual different SDs task.

**Supplementary Figure 19.**
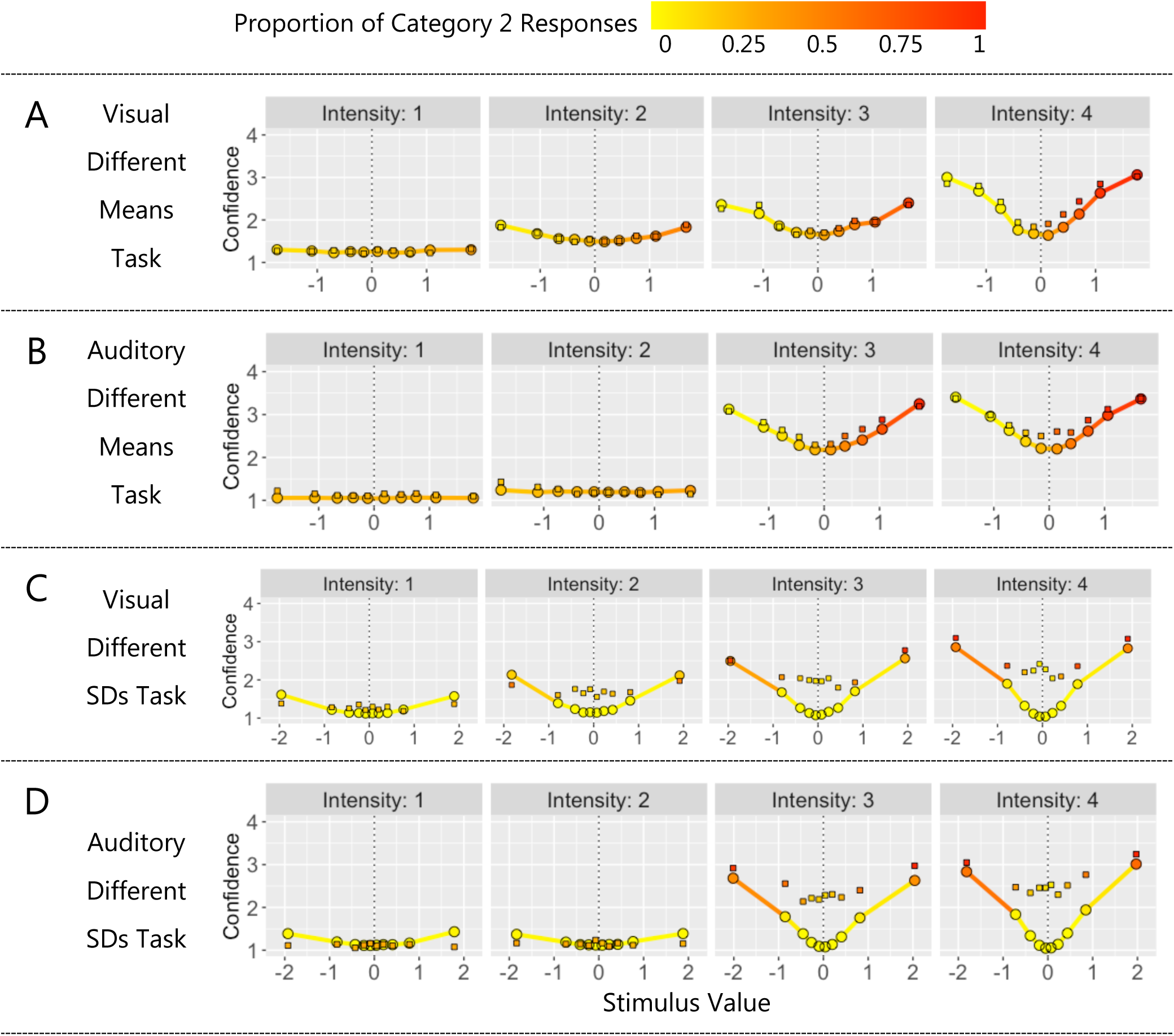
Bayesian Model: Log Posterior Probability Ratio with Decision Noise. In all plots, mean confidence (y axis) and proportion of Cat 2 responses (colour) for binned standardised stimulus values (x axis). Square data points show means for experimental data and solid lines and circular data points show means for model predictions. Model was fit to data from each task and modality separately. Model predictions displayed for: (A) visual different means task, (B) auditory different means task, (C) visual different SDs task and (D) auditory different SDs task.

**Supplementary Figure 20.**
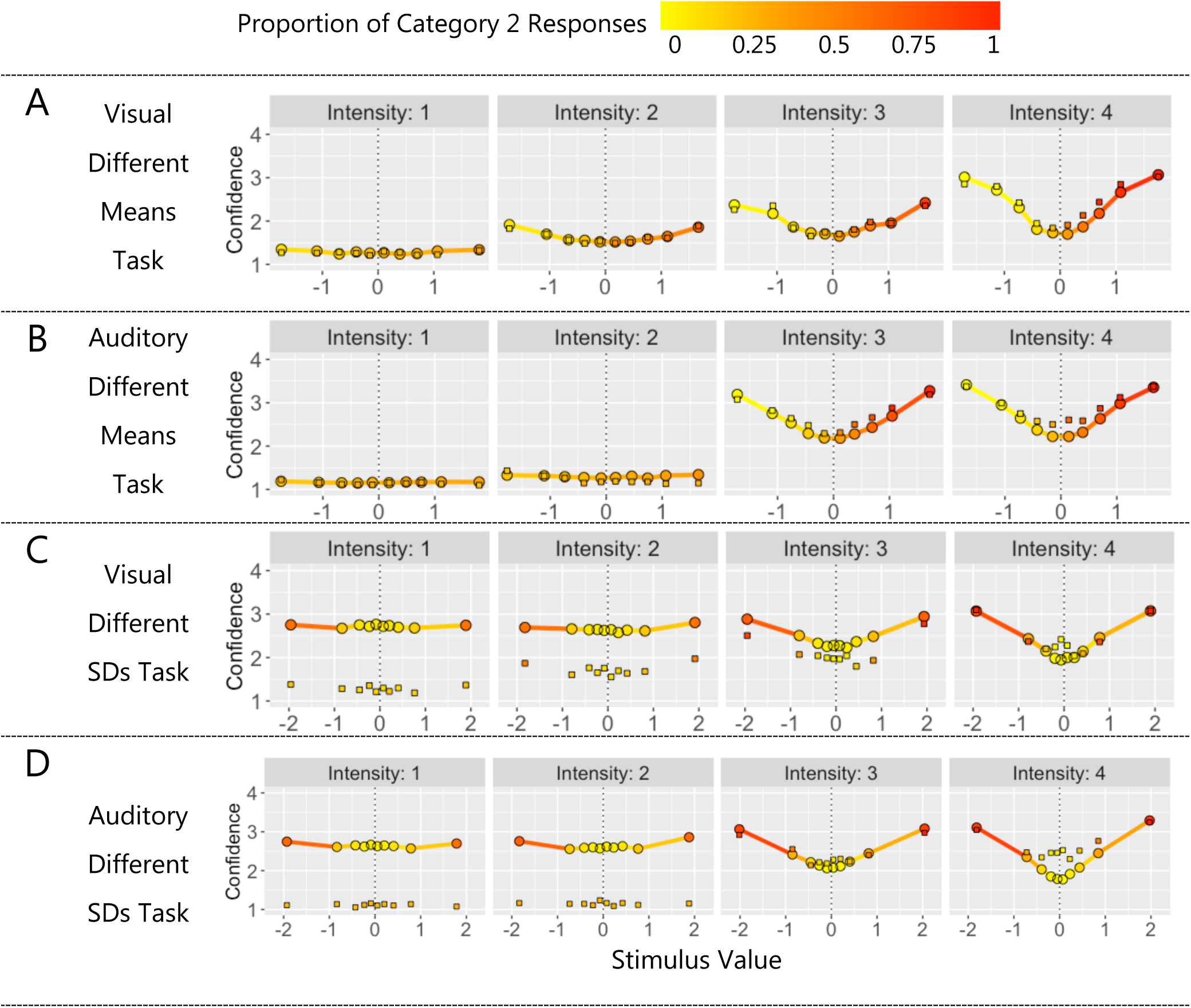
Bayesian Model: Log Posterior Probability Ratio with Free Category Distributions Parameters. In all plots, mean confidence (y axis) and proportion of Cat 2 responses (colour) for binned standardised stimulus values (x axis). Square data points show means for experimental data and solid lines and circular data points show means for model predictions. Model was fit to data from each task and modality separately. Model predictions displayed for: (A) visual different means task, (B) auditory different means task, (C) visual different SDs task and (D) auditory different SDs task.

**Supplementary Figure 21.**
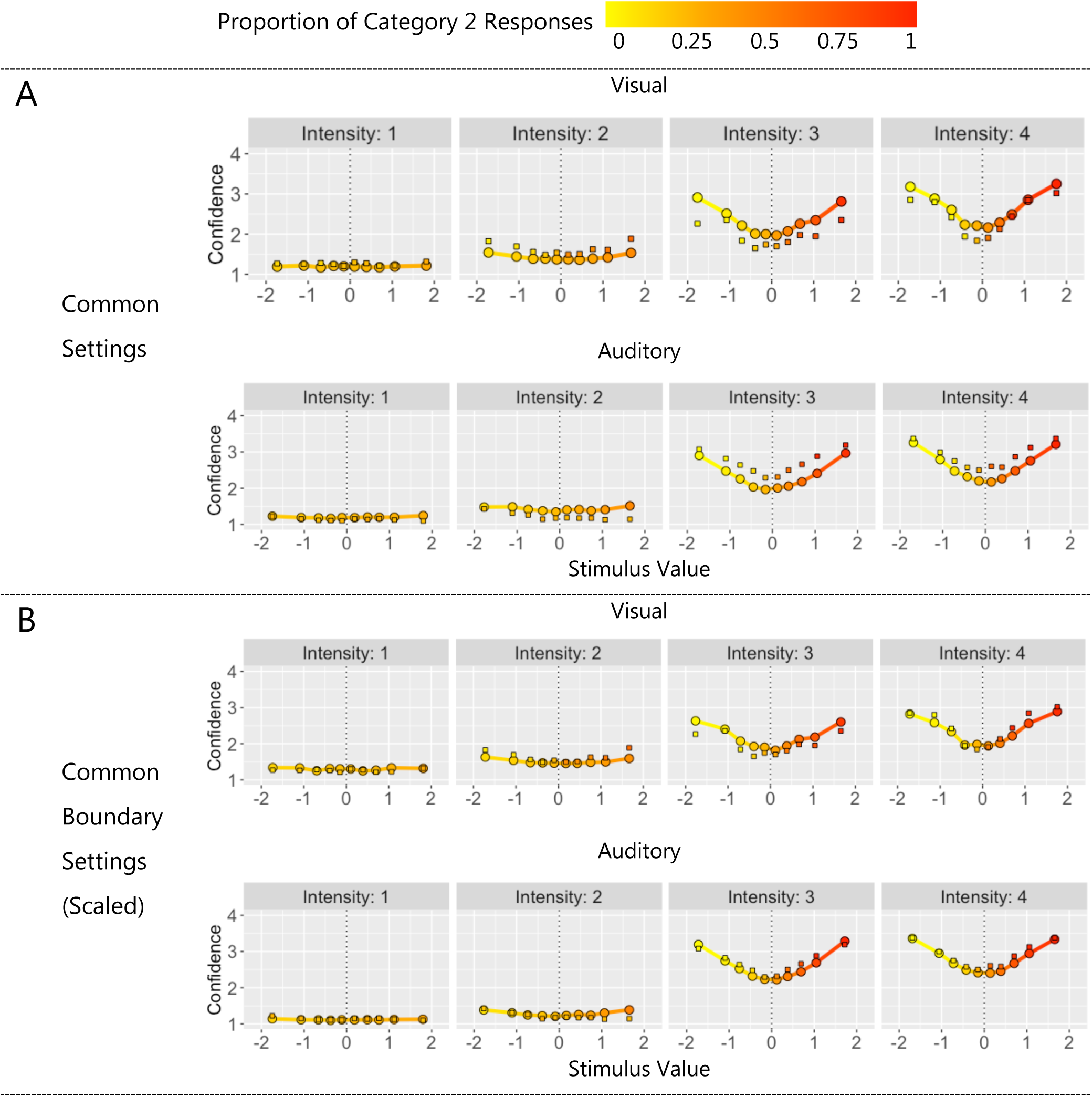
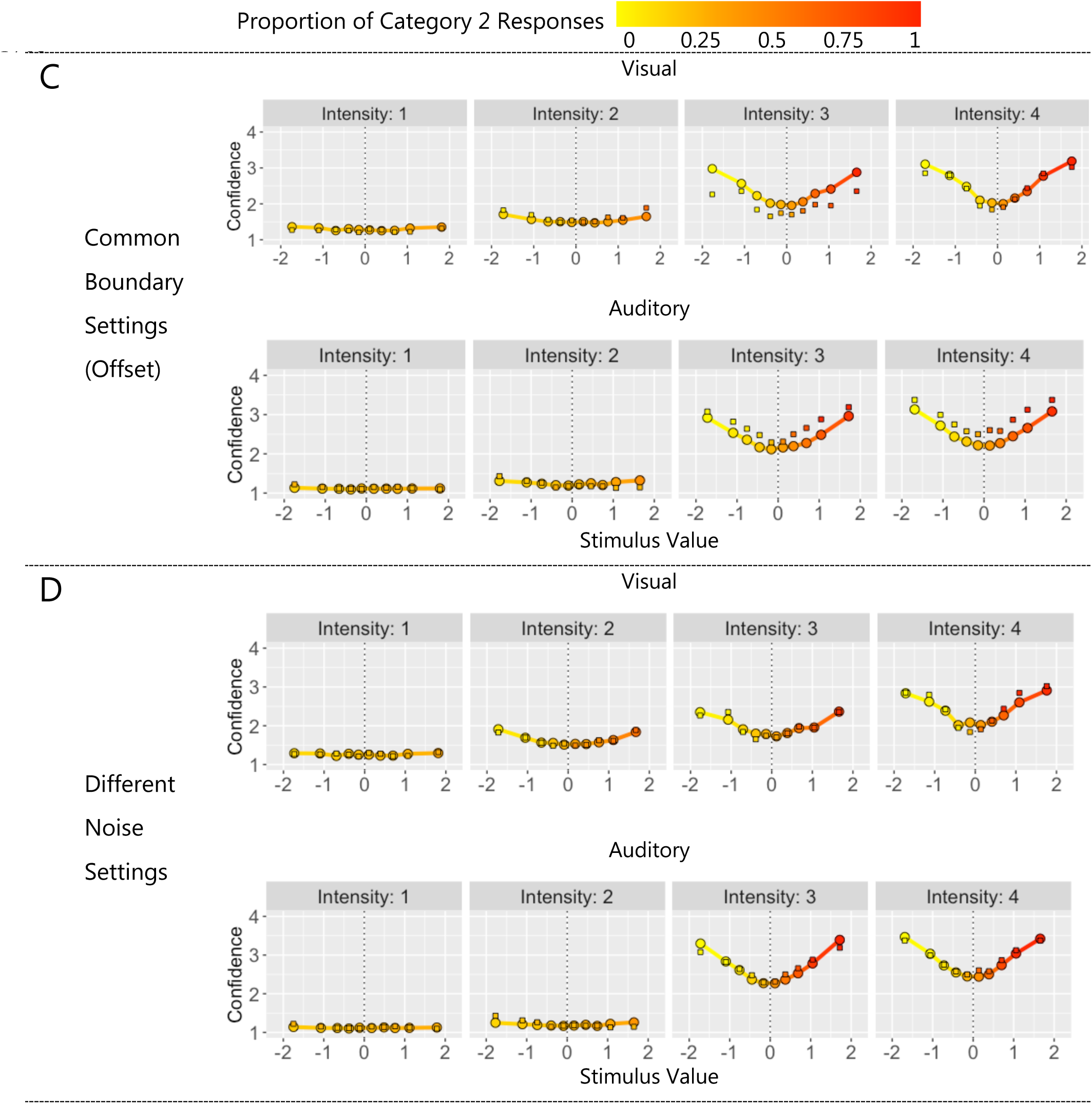
Parameter Settings Across Modalities: Different Means Task. In all plots, mean confidence (y axis) and proportion of Category 2 responses (colour) for binned standardised stimulus values (x axis). Square data points show means for experimental data and solid lines and circular data points show means for model predictions from (A) the common settings model and (B) the common boundary settings (scaled) model.

**Supplementary Figure 22.**
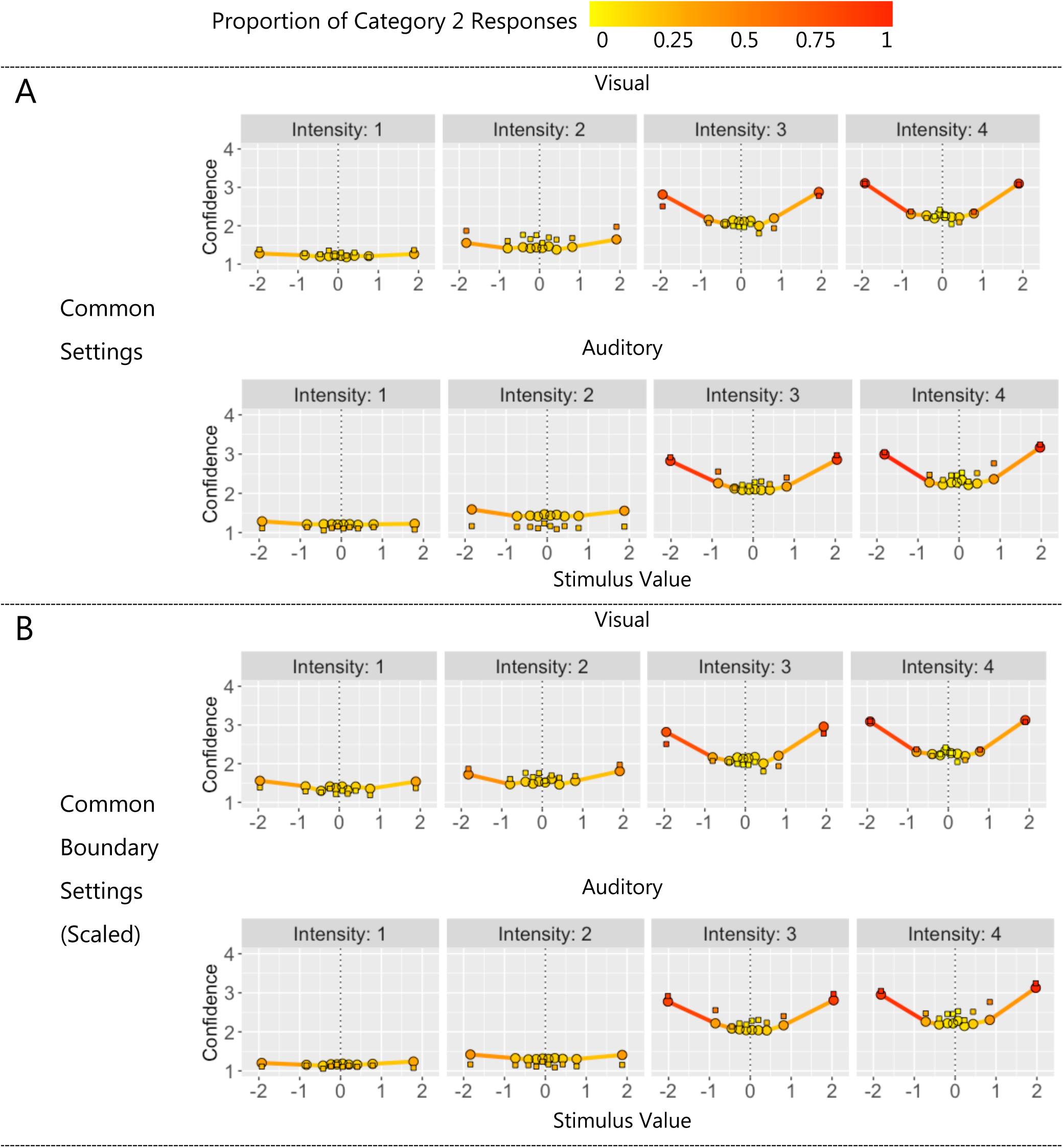
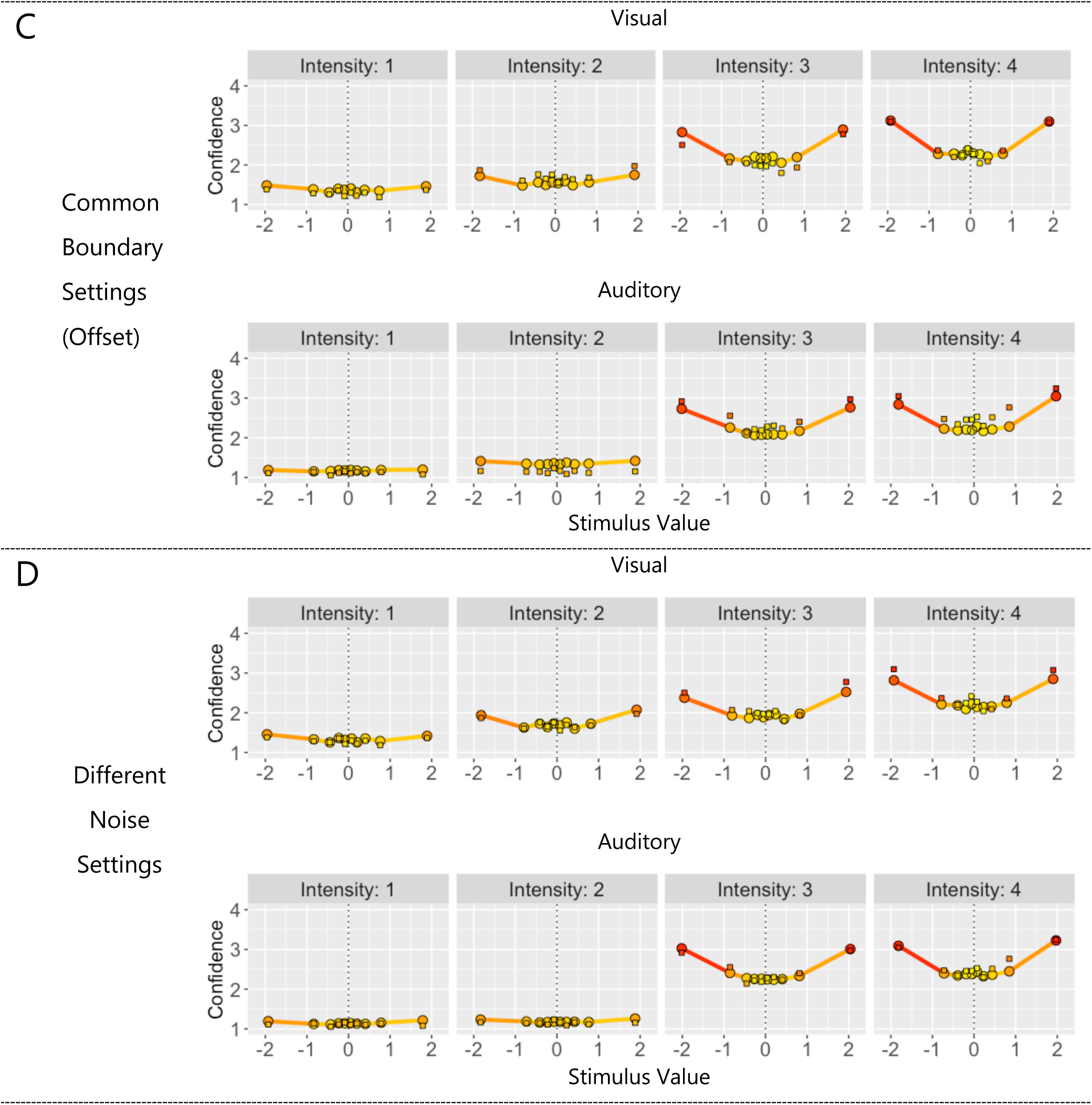
Parameter Settings Across Modalities: Different SDs Task. In all plots, mean confidence (y axis) and proportion of Category 2 responses (colour) for binned standardised stimulus values (x axis). Square data points show means for experimental data and solid lines and circular data points show means for model predictions from (C) the common settings model and (D) the common boundary settings (scaled) model.

## References

Adler, W. T., & Ma, W. J. (2018). Comparing Bayesian and non-Bayesian accounts of human confidence reports. PLOS Computational Biology, 14(11), e1006572. https://doi.org/10.1371/journal.pcbi.1006572

Aitchison, L., Bang, D., Bahrami, B., & Latham, P. E. (2015). Doubly Bayesian Analysis of Confidence in Perceptual Decision-Making. PLOS Computational Biology, 11(10), e1004519. https://doi.org/10.1371/journal.pcbi.1004519

Alais, D., & Burr, D. (2004). The Ventriloquist Effect Results from Near-Optimal Bimodal Integration. Current Biology, 14(3), 257–262. https://doi.org/10.1016/j.cub.2004.01.029

Annevirta, T., Laakkonen, E., Kinnunen, R., & Vauras, M. (2007). Developmental dynamics of metacognitive knowledge and text comprehension skill in the first primary school years. Metacognition and Learning, 2(1), 21–39. https://doi.org/10.1007/s11409-007-9005-x

Baranski, J. V., & Petrusic, W. M. (1994). The calibration and resolution of confidence in perceptual judgments. Perception & Psychophysics, 55(4), 412–428. https://doi.org/10.3758/BF03205299

Barthelme, S., & Mamassian, P. (2010). Flexible mechanisms underlie the evaluation of visual confidence. Proceedings of the National Academy of Sciences, 107(48), 20834–20839. https://doi.org/10.1073/pnas.1007704107

Bellon, E., Fias, W., & De Smedt, B. (2020). Metacognition across domains: Is the association between arithmetic and metacognitive monitoring domain- specific? PLOS ONE, 15(3), e0229932. https://doi.org/10.1371/journal.pone.0229932

Bertana, A., Chetverikov, A., van Bergen, R. S., Ling, S., & Jehee, J. F. M. (2021). Dual strategies in human confidence judgments. Journal of Vision, 21(5), 21. https://doi.org/10.1167/jov.21.5.21

Brainard, D. H. (1997). The Psychophysics Toolbox. Spatial Vision, 10(4), 433–436. https://doi.org/10.1163/156856897X00357

de Gardelle, V., Le Corre, F., & Mamassian, P. (2016). Confidence as a Common Currency between Vision and Audition. PLOS ONE, 11(1), e0147901. https://doi.org/10.1371/journal.pone.0147901

de Gardelle, V., & Mamassian, P. (2014). Does Confidence Use a Common Currency Across Two Visual Tasks? Psychological Science, 25(6), 1286– 1288. https://doi.org/10.1177/0956797614528956

Denison, R. N., Adler, W. T., Carrasco, M., & Ma, W. J. (2018). Humans incorporate attention-dependent uncertainty into perceptual decisions and confidence. Proceedings of the National Academy of Sciences, 115(43), 11090–11095. https://doi.org/10.1073/pnas.1717720115

Deroy, O., Spence, C., & Noppeney, U. (2016). Metacognition in Multisensory Perception. Trends in Cognitive Sciences, 20(10), 736–747. https://doi.org/10.1016/j.tics.2016.08.006

Donoso, M., Collins, A. G. E., & Koechlin, E. (2014). Foundations of human reasoning in the prefrontal cortex. Science, 344(6191), 1481–1486. https://doi.org/10.1126/science.1252254

Ernst, M. O., & Banks, M. S. (2002). Humans integrate visual and haptic information in a statistically optimal fashion. Nature, 415(6870), 429–433. https://doi.org/10.1038/415429a

Faivre, N., Filevich, E., Solovey, G., Kühn, S., & Blanke, O. (2018). Behavioral, Modeling, and Electrophysiological Evidence for Supramodality in Human Metacognition. Journal of Neuroscience, 38(2), 263–277. https://doi.org/10.1523/JNEUROSCI.0322-17.2017

Fitzgerald, L. M., Arvaneh, M., & Dockree, P. M. (2017). Domain-specific and domain-general processes underlying metacognitive judgments. Consciousness and Cognition, 49, 264–277. https://doi.org/10.1016/j.concog.2017.01.011

Fleck, M. S., Daselaar, S. M., Dobbins, I. G., & Cabeza, R. (2006). Role of Prefrontal and Anterior Cingulate Regions in Decision-Making Processes Shared by Memory and Nonmemory Tasks. Cerebral Cortex, 16(11), 1623–1630. https://doi.org/10.1093/cercor/bhj097

Girshick, A. R., Landy, M. S., & Simoncelli, E. P. (2011). Cardinal rules: Visual orientation perception reflects knowledge of environmental statistics. Nature Neuroscience, 14(7), 926–932. https://doi.org/10.1038/nn.2831

Hangya, B., Sanders, J. I., & Kepecs, A. (2016). A Mathematical Framework for Statistical Decision Confidence. Neural Computation, 28(9), 1840–1858. https://doi.org/10.1162/NECO_a_00864

Heereman, J., Walter, H., & Heekeren, H. R. (2015). A task-independent neural representation of subjective certainty in visual perception. Frontiers in Human Neuroscience, 9(551). https://doi.org/10.3389/fnhum.2015.00551

Hillis, J. M., Ernst, M. O., Banks, M. S., & Landy, M. S. (2002). Combining Sensory Information: Mandatory Fusion Within, but Not Between, Senses. Science, 298(5598), 1627–1630. https://doi.org/10.1126/science.1075396

Kepecs, A., Uchida, N., Zariwala, H. A., & Mainen, Z. F. (2008). Neural correlates, computation and behavioural impact of decision confidence. Nature, 455(7210), 227–231. https://doi.org/10.1038/nature07200

Komura, Y., Nikkuni, A., Hirashima, N., Uetake, T., & Miyamoto, A. (2013). Responses of pulvinar neurons reflect a subject’s confidence in visual categorization. Nature Neuroscience, 16(6), 749–755. https://doi.org/10.1038/nn.3393

Körding, K. (2007). Decision Theory: What “Should” the Nervous System Do? Science, 318(5850), 606–610. https://doi.org/10.1126/science.1142998

Koriat, A. (1993). How Do We Know That We Know? The Accessibility Model of the Feeling of Knowing. 100(4), 609–639. https://doi.org/10.1037/0033-295X.100.4.609

Lehmann, M., Hagen, J., & Ettinger, U. (2022). Unity and diversity of metacognition. Journal of Experimental Psychology: General, 151(10), 2396–2417. https://doi.org/10.1037/xge0001197

Lempert, K. M., Chen, Y. L., & Fleming, S. M. (2015). Relating Pupil Dilation and Metacognitive Confidence during Auditory Decision-Making. PLOS ONE, 10(5), e0126588. https://doi.org/10.1371/journal.pone.0126588

Li, H.-H., & Ma, W. J. (2020). Confidence reports in decision-making with multiple alternatives violate the Bayesian confidence hypothesis. Nature Communications, 11(1), 2004. https://doi.org/10.1038/s41467-020-15581-6

Lisi, M., Mongillo, G., Milne, G., Dekker, T., & Gorea, A. (2021). Discrete confidence levels revealed by sequential decisions. Nature Human Behaviour, 5(2), 273–280. https://doi.org/10.1038/s41562-020-00953-1

Locke, S. M., Landy, M. S., & Mamassian, P. (2022). Suprathreshold perceptual decisions constrain models of confidence. PLOS Computational Biology, 18(7), e1010318. https://doi.org/10.1371/journal.pcbi.1010318

Ma, W. J., & Jazayeri, M. (2014). Neural Coding of Uncertainty and Probability. Annual Review of Neuroscience, 37(1), 205–220. https://doi.org/10.1146/annurev-neuro-071013-014017

Masset, P., Ott, T., Lak, A., Hirokawa, J., & Kepecs, A. (2020). Behavior- and Modality-General Representation of Confidence in Orbitofrontal Cortex. Cell, 182(1), 112–126.e18. https://doi.org/10.1016/j.cell.2020.05.022

Mazancieux, A., Fleming, S. M., Souchay, C., & Moulin, C. J. A. (2020). Is there a G factor for metacognition? Correlations in retrospective metacognitive sensitivity across tasks. Journal of Experimental Psychology: General, 149(9), 1788–1799. https://doi.org/10.1037/xge0000746

McCurdy, L. Y., Maniscalco, B., Metcalfe, J., Liu, K. Y., de Lange, F. P., & Lau, H. (2013). Anatomical Coupling between Distinct Metacognitive Systems for Memory and Visual Perception. Journal of Neuroscience, 33(5), 1897–1906. https://doi.org/10.1523/JNEUROSCI.1890-12.2013

Morales, J., Lau, H., & Fleming, S. M. (2018). Domain-General and Domain-Specific Patterns of Activity Supporting Metacognition in Human Prefrontal Cortex. The Journal of Neuroscience, 38(14), 3534–3546. https://doi.org/10.1523/JNEUROSCI.2360-17.2018

Navajas, J., Hindocha, C., Foda, H., Keramati, M., Latham, P. E., & Bahrami, B. (2017). The idiosyncratic nature of confidence. Nature Human Behaviour, 1(11), 810–818. https://doi.org/10.1038/s41562-017-0215-1

Pelli, D. G. (1997). The VideoToolbox software for visual psychophysics: Transforming numbers into movies. 10(4), 437–442. https://doi.org/10.1163/156856897X00366

Qamar, A. T., Cotton, R. J., George, R. G., Beck, J. M., Prezhdo, E., Laudano, A., Tolias, A. S., & Ma, W. J. (2013). Trial-to-trial, uncertainty-based adjustment of decision boundaries in visual categorization. Proceedings of the National Academy of Sciences, 110(50), 20332–20337. https://doi.org/10.1073/pnas.1219756110

Rademaker, R. L., Tredway, C. H., & Tong, F. (2012). Introspective judgments predict the precision and likelihood of successful maintenance of visual working memory. Journal of Vision, 12(13), 21–21. https://doi.org/10.1167/12.13.21

Rahnev, D., Maniscalco, B., Graves, T., Huang, E., de Lange, F. P., & Lau, H. (2011). Attention induces conservative subjective biases in visual perception. Nature Neuroscience, 14(12), 1513–1515. https://doi.org/10.1038/nn.2948

Rinne, L. F., & Mazzocco, M. M. M. (2014). Knowing Right From Wrong In Mental Arithmetic Judgments: Calibration Of Confidence Predicts The Development Of Accuracy. PLoS ONE, 9(7), e98663. https://doi.org/10.1371/journal.pone.0098663

Rouault, M., McWilliams, A., Allen, M. G., & Fleming, S. M. (2018). Human Metacognition Across Domains: Insights from Individual Differences and Neuroimaging. Personality Neuroscience, 1, e17. https://doi.org/10.1017/pen.2018.16

Samaha, J., & Postle, B. R. (2017). Correlated individual differences suggest a common mechanism underlying metacognition in visual perception and visual short-term memory. 284(1867), 10. https://doi.org/10.1098/rspb.2017.2035

Sanders, J. I., Hangya, B., & Kepecs, A. (2016). Signatures of a Statistical Computation in the Human Sense of Confidence. Neuron, 90(3), 499–506. https://doi.org/10.1016/j.neuron.2016.03.025

Song, C., Kanai, R., Fleming, S. M., Weil, R. S., Schwarzkopf, D. S., & Rees, G. (2011). Relating inter-individual differences in metacognitive performance on different perceptual tasks. Consciousness and Cognition, 20(4), 1787–1792. https://doi.org/10.1016/j.concog.2010.12.011

Treisman, M. (1998). Combining Information: Probability Summation and Probability Averaging in Detection and Discrimination. 3(2), 252–265. https://doi.org/10.1037/1082-989X.3.2.252

Zylberberg, A., Roelfsema, P. R., & Sigman, M. (2014). Variance misperception explains illusions of confidence in simple perceptual decisions. Consciousness and Cognition, 27, 246–253. https://doi.org/10.1016/j.concog.2014.05.012

